# Extensive recombination, selection, and asexual blooms shape the diversity of the dominant clade of *Prochlorococcus*

**DOI:** 10.64898/2025.12.29.696893

**Authors:** Alana Papula, Gabriel Birzu, Daniel S. Fisher

## Abstract

The tiny and enormously abundant marine cyanobacterium *Prochlorococcus marinus* contains many levels of population structure, with sequenced isolates spanning four orders of magnitude of diversity. It is divided into several major clades which exhibit the classic structure of distinct species, with little gene exchange across clade boundaries. The diversity within the dominant clade, High Light II (HLII), exceeds that of many bacterial species but exhibits little global population structure beyond some small tight genomic clusters. Few ecological distinctions within HLII have been found. What causes and sustains for millions of years the extensive diversity of this globally mixed population? In this work, hundreds of single cell genomes are analyzed and used to elucidate patterns of core and flexible gene flow, delineate subtle geographical correlations, and quantitatively test several scenarios for the evolutionary history of HLII. We find that recombination connects the entire HLII clade and that genes more than a few kb apart along the genome evolve largely independently. On shorter timescales, primarily-asexual blooms occur, but the clonal backbones of the resulting closely-related cell clusters are steadily eroded by recombination with the rest of HLII. Although a small number of genes have ocean-specific alleles, there is no evidence for genetically-stable ecotypes within the clade. We conjecture that spatiotemporally variable selection drives conditionally-beneficial variants of genomic segments to rise to high abundance while horizontally-transferring throughout the population.

## Introduction

The diversity of bacterial species is well known, but only recently has the surprising extent of diversity *within* species become apparent. How is such intra-species diversity generated and maintained? How can this diversity best be characterized? Historically, bacterial species have been classified using 16S ribosomal RNA. Large-scale whole-genome sequencing enables learning far more. At the whole genome level, is intra-species bacterial diversity mostly asexual with occasional horizontal gene transfers, or are some bacterial populations better characterized as *quasi-sexual*, with individual genomes roughly a random grab-bag of rapidly exchanging genetic elements (*1*, *2*)? At the genic level, how varied are core genes that are present in all members of the species? Do their variants reflect whole genome structure – as oft assumed for 16S rRNA? How does flexible gene content vary, and on what timescales?

Most importantly, what are the evolutionary and ecological processes that shape intra-species diversity? For bacteria that live in complex environments with complex geographical structure, such as gut bacteria, it is plausible – although far from clear – that there are a great number of ecological niches with slow geographical mixing. What about bacteria that live in relatively simple environments which mix rapidly? *Prochlorococcus marinus* is an extreme example whose diversity presents a great puzzle. It also presents a great opportunity: other than some human pathogens, it is one of the few bacterial species about which geographical mixing processes are quantitatively known.

*Prochlorococcus* is a tiny cyanobacterium which dominates photosynthesis in nutrient poor tropical oceans. Alone, it is responsible for fixing about as much carbon as is emitted by activities of the human race (*3*). To understand the robustness and response to climate change of the oceans requires understanding the current diversity of *Prochlorococcus* and the species’ evolutionary history over tens of millions years. The surface layers of the ocean are mixed globally in tens of years (*4*) and *Prochlorococcus* populations are globally connected (although likely were not during ice ages). Thus, potentially, all individuals can directly compete and a unilaterally-beneficial genetic change would rapidly sweep globally. Due to its enormous population size, ∼ 10^27^ cells (*5*), even complex genetic changes arise all the time. The typical generation time is on the order of one day (*6*), so that a genetic change with a 1% fitness advantage could take over in tens of years. Such sweeps would eliminate genetic diversity, but the diversity of *Prochlorococcus* spans a huge spectrum of time and genetic scales (*3*, *7*, *8*). In such a well-mixed population living in a seemingly simple and relatively uniform environment, why doesn’t “survival of the fittest” continually eliminate most of the diversity?

Several qualitative explanations for intra-species diversity are often given. The simplest (based on population genetics) are “neutral” explanations with demographic and geographic fluctuations driving the population dynamics (*9*–*11*). However, due to the large population size and absence of tight bottlenecks, the time required for mutations to reach observable frequencies through drift is absurdly long. Thus the observed genomic diversity must be driven by differential selection. But all ubiquitously-beneficial allelic variants will have already swept through this old, well-mixed population; thus observed genetic differences can be only *conditionally* beneficial, depending on environmental conditions, ecological interactions, and epistatic interactions with other genetic differences.

The traditional ecological explanation for intra-species bacterial diversity is the existence of discrete ecological niches, each occupied by a distinct *“ecotype”* : an evolutionarily-stable group of ecologically-similar bacteria. Key to the ecological explanation of diversity is that adaptations that are advantageous in a particular ecological niche can sweep through an ecotype, eliminating diversity within it, but not spreading to other ecotypes, with recombination across ecotype boundaries rare due to differing selection pressures and barriers imposed by genetic differences (*12*). While some ecotypes may go extinct and others may branch to yield new ecotypes, crucially, ecotypes are assumed to remain distinct and not blend in to the rest of the population via gene flow. However, explaining finer and finer scale genetic diversity would require hypothesizing finer and finer scale niches. At what point does the delineation of ecotypes – even in principle – fail to be a useful framework? Surely before the yoctoniche level, where each distinct genome occupies its own niche! We will argue that a *persistent* fine-scale ecotype structure is not supported by the *Prochlorococcus* data.

How might the ecotype hypothesis be tested? Cells belonging to the same ecotype should share alleles at certain ecologically-relevant genes that together enable them to adapt to their specific ecological niche; thus the presence of certain alleles should be predictive of the presence of other alleles at some set of genes scattered along the genome. Ecotype-determining genes would be *strongly linked.* In populations undergoing minimal recombination, linked alleles could result from genome sweeps within a niche (as proposed for *Prochlorococcus* in Kashtan 2014 (*13*)). In populations with extensive recombination, ecological specialization could perhaps be maintained through epistatic interactions among a smaller number of genes, with other genes apparently freely mixing across ecotypes. Whole-genome linkage information is therefore essential for testing whether diversity can be explained by ecotypes and for analyzing the crucial role of gene exchange. A large number of whole genomes are needed. Isolate strains can be used to study readily culturable bacteria, but for difficult to culture organisms, such as *Prochlorococcus*, this is not viable.

Using the conventional 16S definition, all *Prochlorococcus* comprise a single species. Several major clades – High Light (HL) I, II, and Low Light (LL) I, IV, etc.– were identified initially, from physiological and ecological differences (*3*, *14*). The criterion used to define distinct species for sexual organisms is lack of inter-species genetic exchange; by this definition, these major clades (as we show) are distinct species. Finer scale groupings and trees of *Prochlorococcus* cells were made based on ITS, a higher diversity spacer region in ribosomal RNA which enables resolution of finer structure (*3*, *13*, *15*–*18*), but it has not been shown whether structure at the ITS correlates with either ecological differences, or other genomic differences, at scales typical of divergences within *Prochlorococcus* clades.

Here, we focus on HLII, the most abundant major of *Prochlorococcus*, itself more diverse than many bacterial species. The puzzle of diversity is most extreme within this clade. Investigations of whether ITS clusters within HLII correlate with environmental variables – which would indicate possible ecotypes – have yielded minimal correlations (in contrast, some have been found within HLI) (*18*) (Figure S11). The difficulty of culturing *Prochlorococcus* has impeded whole-genome studies, but the 85 lab cultures, which span the diversity of *Prochlorococcus*, have yielded 17 HLII “reference” genomes (*8*, *19*, *20*). Metagenomics is widely used to study bacterial diversity but is limited by read length, and the “gold-standard” of assembly fails with extensive coexisting intra-species diversity and cannot provide whole-genome linkage information (*21*, *22*). Single-cell genomes combine the advantages of both isolates and metagenomes, as they are not biased towards lab growth and provide crucial information on whole-genome scales (*23*). We use single cell amplified HLII genomes (SAGs) from two complementary datasets, one of 489 *Prochlorococcus* cells of which 167 are HLII cells quasi-uniformly sampled (Figure 1 (b)): the *quasi-random* dataset (*24*) (SM 1. A.). The other dataset contains 90 closely-related HLII cells from deep sequencing of an ITS-based cluster of cells from one location (Figure 1 (c)): the *deep clustered dataset* (*13*). Although the SAGs are far from complete – typically 50-90 % – the high degree of synteny characterizing HLII genomes makes it possible to evaluate genome-scale linkage (Figure S7) and the low error rate (SM 1. A.) enables reliable identification of very small differences.

**Figure 1:**
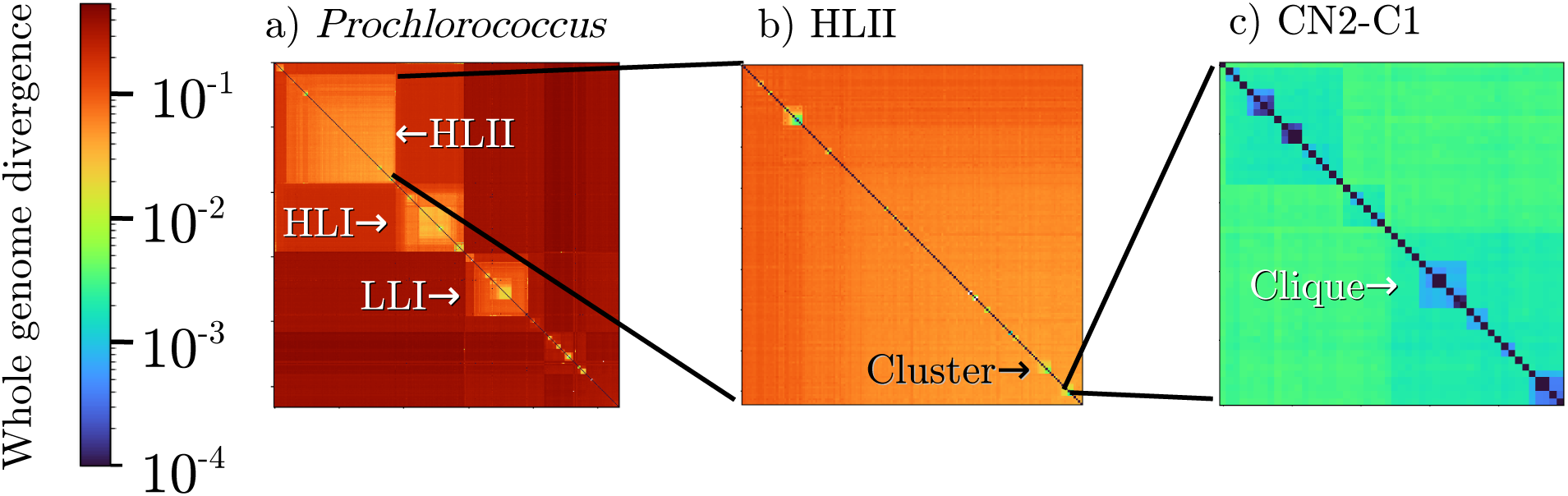
Diversity of *Prochlorococcus* revealed in whole-genome pairwise divergence matrices between Single Cell Amplified Genomes (SAGs) ranges over four orders of magnitude (*13*, *24*). (a) At the highest level are major clades: High Light (HL) and Low Light (LL) groups HLI, HLII, LLI, etc., each sufficiently diverse and genetically isolated to be considered separate species. (b) Within HLII most cells are uniformly diverged, but multiple small tight clusters of more closely-related cells exist. (c) One deeply sequenced cluster, CN2-C1, contains several cliques of 3-7 cells that are identical along the majority of their genes with clonal-backbone divergences as low as 10*^−^*^4½^. (a) shows all SAGs sequenced by Berube et al. (*24*), (b) shows just the quasi-randomly sampled HLII SAGs from this dataset, (c) shows C1, one of 5 deeply-sampled clusters in a set of HLII cells sequenced by Kashtan et al. (*13*). In (a) and (b) mean core exome divergence is used as a proxy for whole-genome divergence; in (c) clonal backbone divergence is used, roughly a factor of 4 less than the mean core exome divergence of these cells (C1 has a diameter of 0.4% using clonal backbone divergence and 1.6% using mean exome diameter).

### Many levels of HLII whole-genome diversity

The multiple levels of *Prochlorococcus* genomic diversity can be visualized in whole genome distance matrices – defined here as the average core gene nucleotide divergence between pairs of cells (Figure 1). At the highest phylogenetic level (a), the major clades (“species”) are clearly distinguished, but the quasirandom sample of HLII cells (b) has little structure. Most pairs of HLII cells are comparably diverged (Figure S11), but several *tight clusters*, each comprising a few relatively closely-related HLII cells, are found. The Kashtan 2014 dataset contains deeper sequencing of five tight clusters named CN2-C1 to CN2-C5 (henceforth simply C1, etc). These were identified by similar ITS (as part of a larger ITS-based cluster denoted CN2) but are comparably diverged from each other as typical HLII cells at the whole genome level (Figure S14). Seven SAGs from the quasi-random dataset fall into one of these deeply-sampled clusters, including three clustering with C1, the cluster with the most (51) sequenced cells (Figure 1 (c)). Within C1-C5 are finer-scale structure, including six *cliques* of 3-7 cells in which all cells are identical at over half of their genes.

HLII has higher nucleotide diversity than most well-studied bacterial species, which are typically diverged 1-3% (*3*, *25*, *26*)). Typical core genome nucleotide divergences between HLII cell pairs fall between 5-10% (typical HLII amino acid divergences are 4-9%), but the tight clusters have a range of nucleotide divergence diameters from 0.1 - 4%. At twofold and fourfold degenerate synonymous sites, HLII divergences average 10% and 20%, respectively, and only 2.6% of fourfold degenerate sites are monomorphic. The narrow distribution of most whole genome divergences makes phylogenies difficult to construct, producing a star-like tree with some clusters of leaves. Although a large number of “defining” SNPs might be expected to aid in tree construction, most mutations have occurred multiple times – homoplasy. In any case, we shall see that whole-genome “phylogenies” are not meaningful. Even at the level of individual genes, phylogenies are highly non-parsimonious and uncertain. We therefore primarily use statistical information, such as SNP statistics and pairwise divergences, rather than processing the data with phylogenetic or other methods that introduce unjustifiable assumptions.

Synonymous mutations provide a rough “clock” to relate a cell pair’s genomic divergence at a particular region of the genome to the time since the pair’s common ancestor at that region. The mutation rate of a HLII lab strain was measured to be 10^−9½^ mutations per bp per generation (*10*). With roughly one generation per day and the proportionality factor observed between synonymous and all-nucleotide divergences (SM 3. A.) this translates to mutational genomic differences accumulating at roughly 10^−7^ per year, with perhaps an order of magnitude uncertainty. Thus typical HLII cells diverged 10^5½−6½^ years ago, and the closest pair of cells sequenced around ∼ 10^2−3^ years ago – already more than the global surface-ocean mixing time.

The structure of the diversity of HLII raises many questions. How can such high diversity be maintained in this seemingly well-mixed population? What gives rise to the largely-uniform structure of HLII at the whole genome level? Are tight clusters within HLII stable and long-lived, or do their frequencies or even existence change over time? Are these tight clusters (or other groupings of cells) genetically isolated or does recombination erode the differences between them? Do they correspond to distinct ecological niches? Linkage information can be leveraged to ask these and more specific questions including correlations between core genes along the genomes, and the presence or absence of flexible genes.

### Disentangling mutation and recombination

Diversity is generated by mutation on clonally inherited genomes, homologous recombination of core genes, and horizontal gene transfer of flexible genes. It is difficult to distinguish between the contributions of these processes in highly-diverged cells; however, the closely-related cells from the deep clustered dataset provide opportunity to disentangle and quantitatively compare these processes’ effects. For this analysis, we use the multiple sequence alignment of Kashtan et al., aligned using a composite reference comprised of 12 long overlapping contigs from C1 SAGs (*13*).

The most closely-related cell pairs in C1, comprising the close cliques, have divergences from ∼ 10^−4½^− 10^−3^ along much of their genome, but long low-divergence regions are interspersed with occasional short segments with much higher divergences (Figure 2 (a)). These high-divergence regions account for more than half of the genome-wide divergence of closely-related cell pairs, despite only covering a few percent of the genome (Figures S23, S25). This structure suggests that high-divergence segments are homologous recombination events atop a low-divergence clonal backbone.

**Figure 2:**
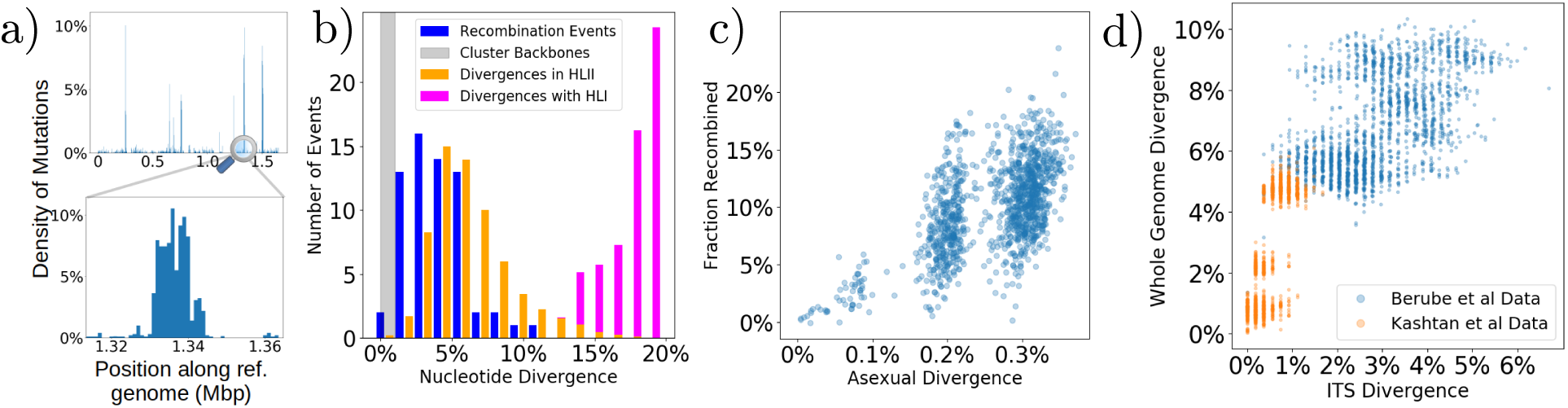
Recombination overwriting clonal backbones: (a) Alignments of closely-related cell pairs: low mutational-divergence clonal-backbone regions are interspersed with high-divergence recombination events. (b) Donors of recombination events into C1 span the diversity of HLII; the divergence distribution of recombined events (blue) is similar to available divergences within HLII (orange) at co-located genomic regions. Divergences with HLI (pink) are larger, with little recombination between clades, although one 35% divergence recombination event was found (not shown). (c) Fraction of genome recombined of closely-related cell pairs (in C1) is proportional to pair’s clonal backbone divergence. Extrapolating suggests that over typical HLII divergence timescales, genomes have been completely overwritten by recombination. (d) Divergences of small genomic regions, such as ITS (shown here) or single genes, are poor predictors of whole genome divergence for typical HLII cell pairs. The CN2 group (orange) forms a 2% diameter ITS cluster and comprises 5 tight genome-level clusters C1-C5 (*13*) which together span a significant portion of the diversity of HLII (blue) at whole genome level. At genome divergences of ∼4%, correlation with ITS divergence is lost, consistent with extrapolation from (c).

To test whether the low-divergence regions represent a clonal backbone (or clonal “frame”), we examine features on short and long genomic length scales. For each cell pair, the fraction of mutually-covered kb segments that are identical gives an estimate of the mean backbone divergence. The distribution of SNPs per kb in the low-divergence regions compared well with a Poisson distribution with this mean (Figures S21, S22), consistent with each cell pair sharing a single clonal backbone. A further test uses long identical segments, up to ∼100 kb in some cell pairs. The length distributions showed long exponential tails with a length scale that varied across pairs (Figure S20), consistent with the prediction from the same Poisson mutation process. We conclude that low-divergence regions between cell pairs in tight clusters such as C1 represent clonal genomic backbone, and are not the result of recent recombination (*27*).

For a clonal backbone to be found in more than one SAG, cells sharing it must have descended from a primarily asexual *bloom*, a selection-driven expansion whose growth outpaces recombination. The bloom must have been large enough, persistent enough, and recent enough for the backbone to still be recognizable. Punctuating the low-divergence clonal backbones of pairs from C1 are recombination events manifesting as discrete, distinct, high-divergence regions. We identify and characterize these within the close cell cliques, and, more conservatively, across C1 (SM 5. A.). Putative recombination events were manually validated using genome alignments and putative recombination breakpoints identified. Many short high-divergence regions that could not be confidently distinguished from alignment errors were conservatively disregarded. This method identified 65 distinct high-divergence homologous recombination events in the 51 C1 cells. Transferred regions ranged from 500 bp to 59 kb, with a median of 7.5 kb (Figure S27) (these are similar lengths to tycheposons, transposons found in *Prochlorococcus* (*28*) believed associated with recombination). Donors of recombination events spanned the divergence of the entire

HLII clade. Typical nucleotide divergence between donor and recipients was 2-8% (4-12% synonymous divergence)(Figure S26), although with proportionately fewer events at high divergences (Figure 2 (b)). These are similar to divergences of recombination events found in studies of some other bacterial species (*29*, *30*).

Not surprisingly, the flow of genetic material goes both into the tight clusters and out of them. Alleles common to one of the C1-C5 clusters were found in HLII cells at ∼13% of core genes examined (SM 5. A.). This raises a key question: in spite of some exchange between them, are the deeply sampled clusters substantially genetically isolated from the rest of the population? Are even HLI and HLII largely isolated?

At very large genomic divergences, homologous recombination can be inhibited by mechanistic obstacles (*31*, *32*). Even mechanistically-successful events may be inhibited by functional incompatibilities – a putative source for maintenance of ecotypes. At what scale is extensive recombination limited in *Prochlorococcus*One event directly identified in C1 was from outside HLII, and in the broader quasirandom dataset, much more evidence of high-divergence events is seen: 59 core genes have two or more well-separated non-singleton alleles that are at least as diverged as HLI is from HLII, and over 100 core genes have singletons that are similarly diverged. Nevertheless, the genetic exchange between HLI and HLII is very limited and identifying them as separate species is justified. Is this true on finer scales of differences?

At the opposite extreme, there are some genes with cells sharing closely related alleles from a partial genic sweep. With cells that form tight clusters at the whole genome excluded, there are clusters of identical alleles in ≥5 cells at 6% of core genes. Evidence for partial sweeps 10^2½^ times more recent than typical HLII divergence times is also found: for example, a core gene where 23 HLII SAGs not in tight clusters are identical over 180bp, and a 2239 bp segment identical in two distantly related cells, both suggesting events 10^3−4^ years ago, However, no well-covered core genes have diameter less than 5%: there is no evidence of full genic sweeps on time scales substantially less than the age of HLII. This is consistent with the expectation that any genetic change that is unconditionally beneficial will have swept even further in the past. It is likely that partial sweeps are temporary, reverting or being overwritten as conditions and genomic contexts change even slightly.

We now show that the overall effects of genetic exchange within HLII build up steadily over evolutionary time. Relying on directly identifying recombination events would lead to large underestimates in the fraction of genomes recombined. Thus we measure for each cell pair in C1 the extent of recombination by the number of kb segments with divergences that could not be explained by the asexual Poisson model (similar to methods in e.g. (*13*, *25*, *30*, *33*)). The estimated recombined genome fraction between a pair grows roughly linearly with the inferred clonal backbone divergence (Figure 2 (c)). This suggests an *apparent* rate of core-genome homologous recombination *into* C1 of about one gene per 10^−4½^ asexual divergence (∼ 10^2−3^ years). Using this, the extent of flow of genomic material *out* of C1-C5 can be estimated assuming there is indeed free recombination across HLII. This yields estimates for the total number of C1-C5 alleles found in unrelated cells roughly consistent with how long these clusters appear to have existed, their abundance, and the rate at which genomes are overwritten by recombination (SM 5. B.).

Quantitative inferences of *actual* recombination rates are very challenging, as observed “apparent” rates are “successful”: in order for a recombination event to be observed, it must not have been deleterious and, indeed, likely conferred some fitness advantage, especially if found in more than one cell and thus rose in abundance. A conventional comparison is the ratio of apparent recombination initiation rate (the number of recombination breakpoints per bp per generation) to mutation rate (denoted *<u>^ρ^</u>*), is here ∼ 10^−2^ (Figure S23). This is at the low end of the range found in other bacteria, from 10^−2^ (Bacteroides vulgatus between clades; Bacteriodes eggerthii) (*30*) to 10 (thermophilic Synechococcus) (*1*, *2*). An alternative measure, the ratio of the number of SNPs per cell pair contributed by high divergence recombination events to the number of SNPs resulting from mutation on a clonal backbone (*25*), is here about two. Extrapolating the trends in Figure 2 (c) suggests that the entire genome is overwritten by recombination by the time cells are 3% diverged. At 5-12% divergence, typical HLII pairs should thus share no clonal backbone. We turn to statistical methods to investigate effects of recombination in cell groups that share little or no residual clonal background.

### HLII as a quasi-sexual population

The impact of recombination on HLII is large. Does this erase memory of the clonal backbones of old asexual blooms? Does this erase the memory of structure that might be associated with distinct ecotypes? More concretely, are there important residual linkages that survive extensive exchange of most of the genome? To address these questions, we analyze correlations along the genome, focusing on correlations between cell pair divergences among core genes. Without recombination, the entire genome shares a single asexual tree, and cell pair divergences at most genes should be highly correlated (Figure 3 (a), (b)). For primarily asexual ancestry, the correlation between genes will be affected by the shape of the phylogeny (at the extreme of star-like phylogenies, no correlations) and be lower if multiple mutations have occurred at the same site in the history of the sample, but – crucially – the correlations between genes will not decrease with separation between them. Conversely, if recombination played a large role in the population’s history, there would be distinct local phylogenies at different parts of the genome (Figure 3 (c), (d)). The probability that a recombination event has occurred between two genes, altering the local phylogeny, scales with the separation between the genes and the correlation of cell pair divergences decreases. However, on scales longer than recombining segments, this decay will saturate at a level that reflects the extent of memory of some asexual history of the population.

**Figure 3:**
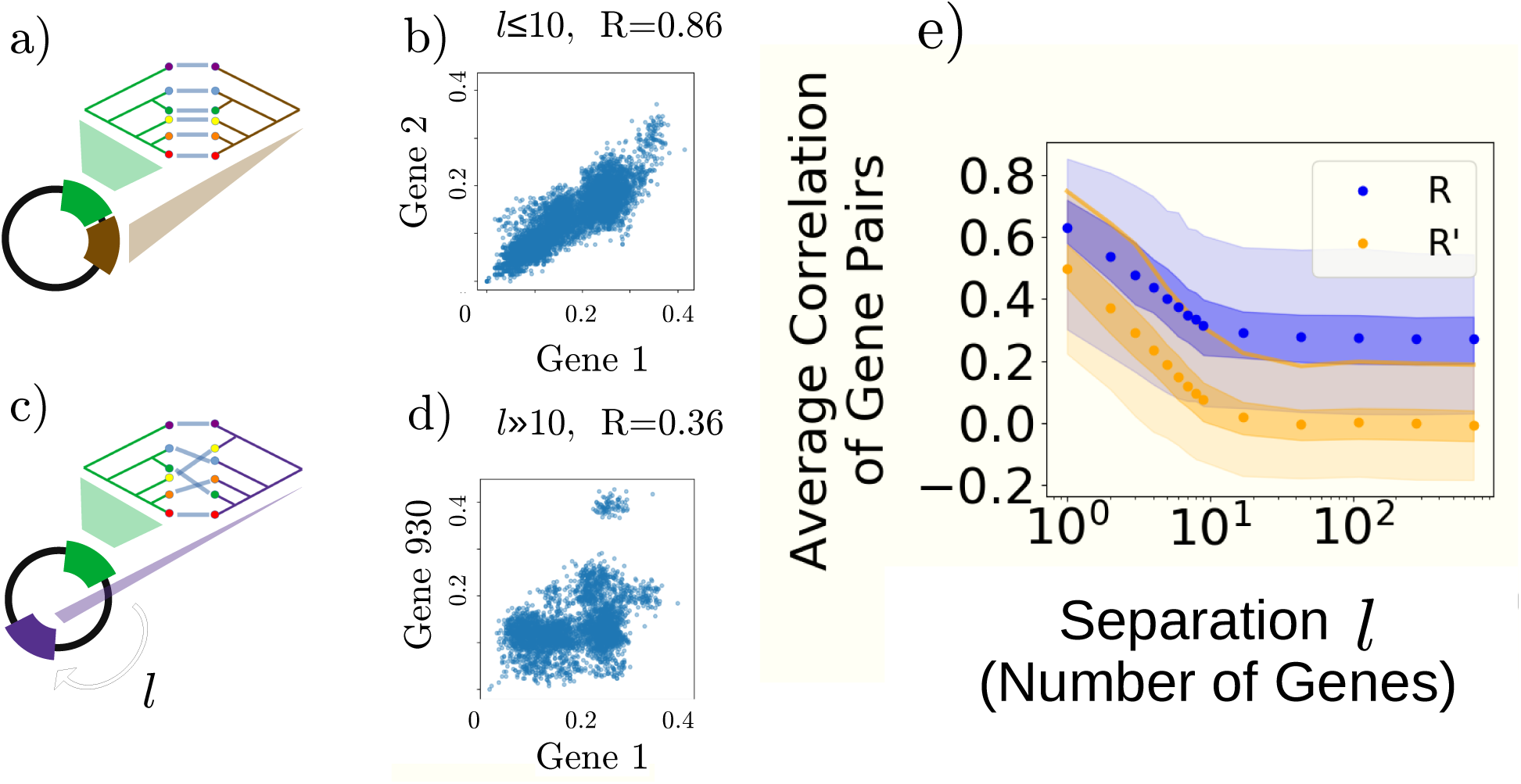
Linkage correlations along core HLII genome probed by pairwise synonymous divergences of genes: (a) Caricature of similar phylogenetic trees of two nearby genes. (b) Cell pair divergences for a highly-correlated pair of nearby genes. (c) Caricature of very different phylogenetic trees of two far-apart genes. (d) Cell pair divergences for a weakly correlated far-apart pair of genes. (e) Average Pearson correlation coefficients, *R* (blue), of cell-pair synonymous divergences at gene pairs, binned by distances, showing decay over ∼10-gene separation, then saturation, with shadings 1 and 2-sigma ranges. Average correlations, *R̃* (orange), of cell-pair genic divergences normalized by whole-genome divergences: *R̃* decays nearly to zero showing that the residual correlations in *R* (blue) are largely due to population structure or weak epistasis rather than to a small number of anomalously highly-linked genes (SM 6. B., Figure S37). This is supported by *R* decaying to ≈0.04 for a group of HLII cells with roughly uniform whole genome divergences (Figure S36).

To analyze correlations among widely diverged groups of cells for which alignments can be problematic, we focus on well-conserved core genes. Orthologous gene groups were constructed by BLASTing SAG ORFs (called and annotated by JGI’s Integrated Microbial Genomes (IMG) system (*34*, *35*)) to a set of conveniently defined “HLII core genes” (SM 1. B.) (this is facilitated by the lack of duplicate genes on the streamlined HLII genome (*7*)). Attempts to construct phylogenies of the core genes yield many ambiguities caused by the homoplasy and star-like character of many genes (SM 10. A.). Trying to correlate such phylogenies is fraught with problems. We thus use the genetic divergences directly.

For each gene, synonymous nucleotide divergences were computed between all cell pairs in the quasirandom set for which that gene was covered (typically in ∼70% of cells). The Pearson correlation coefficient, *R*, between these divergences was computed for all pairs of genes. Cell pair divergences are highly correlated for nearby genes, but decrease from an average *R* ≈ 0.6 to *R* ≈ 0.3 at *>*10 genes separation (Figure 3 (e)). Linkage decay at short distances is probed by correlations between pairs of SNPs: ⟨*r*^2^⟩, *σ*^2^, and divergence-divergence correlations (closely related to *R*; see SM 6.) (Figures S29-S31) all show linkage decay within genes by close to a factor of two in excess of the random null (and saturation at *>*10 genes separation). However, within the whole of HLII, there is a substantial residual correlation at large separations. Is this due to memory of asexual history? To weak genome-wide epistatic interactions? Or to a small minority of highly-linked genes – potentially indicative of distinct ecotypes? To distinguish these scenarios, cell pair genic-divergences were normalized by their average core genome divergence, and the correlations of these normalized divergences, *R̃*, computed. If correlations were dominated by a small number of highly linked groups of genes, *R̃* at large separations would be bimodal with a nonzero mean (SM 6. B., Figure S37). Instead, *R̃* decays to zero at large separations, suggesting that nearly all residual correlations are captured by whole genome population structure. Stronger evidence is found in the *distribution* of *R* values for well-separated genes, which is unimodal with no sign of multiple strongly linked outlier pairs (Figures S34 and S35). As the spectrum of whole-genome divergences dominates linkage measures, we also study a *quasi-equidistant subset* (Figure S12) of 68 cells with pairwise divergences 4.4% to 7.4% (mean 5.8%, std 0.5%) for which the average *R* decreased to 0.04 at long separations (Figure S36).

The lack of substantial linkage in the quasi-equidistant set of cells on genomic length scales suggests that this population is better characterized as *quasi-sexual*, with each cell consisting of an allele at each gene chosen randomly and, if far apart, independently. In the whole of HLII, there are frequent changes of phylogeny along the genome but enough variation at the whole-genome level for alleles of core genes to be weakly correlated (*25*). Nevertheless, many aspects of the large majority of HLII cells are, to a reasonable approximation, quasi-sexual. However, within the population there are several tight clusters of cells that share a substantial clonal backbone. What this structure suggests about the evolutionary history of HLII is a key question we shall return to.

#### Flexible genes

Is it possible that there are clearly delineated subpopulations characterized by their flexible gene content, with the core genes adjusting – in unrecognizable ways – to work best with key ecotype-defining flexible genes? After removing the “HLII core” genes, ORFs are BLASTed all against all and groups of orthologous genes constructed based on nucleotide identity and coverage cutoffs (SM 1. B.). Most flexible genes are singletons, suggesting much hidden diversity of the gene repertoire. But some are almost “core,” found in a substantial fraction or even most cells. We thus consider as “flexible” only genes found in ≤ 34 of the quasi-random SAGs (roughly corresponding to a trough in the distribution of cells per gene) (Figure S8).

Many flexible genes are found in islands, hyper-variable regions of the genome a few tens of kb in length, but others are scattered throughout the core genome. As the small genomes of *Prochlorococcus* – 1.7 Mb for HLII – suggest genome length is under selective pressure, the number of flexible genes in a cell typically ∼600 occupying ∼ <u>^1^</u> of the genome – is likely also. Thus the flexible genes that are present at least the non-singletons – are likely positively selected for. Typical quasi-random SAG pairs share 1 to 25 flexible genes (Figures S9, S10). Flexible gene turnover is very fast, yet genes that occur in multiple cells typically have divergences similar to HLII core genomes and thus are likely to have existed in HLII for much of its history. However, a few percent of flexible genes exhibit clusters and divergences at least twice as high, suggesting that they – or a close relative – was transferred into HLII multiple times from some more-diverged population. At the opposite extreme, a few percent of flexible genes have very low divergence (sometimes lower than 1%, even for typically-diverged HLII cells) across all cells in which they are found: these are evidence of relatively recent partial sweeps (Figure S38).

More quantitative information on the dynamics of flexible gene turnover is evident in closely-related cells. In C1, flexible genes were lost and gained about ten times as fast as core genes recombined (Figure S10) (possibly mediated by mobile genetic elements (*28*)). By ∼ 10^−2½^ divergence, typical within *C*1, the flexible genes of two cells are completely different. This is consistent with previous work showing that flexible gene content does not reflect whole genome divergences, even among closely-related cells (*36* –*38*). The lack of persistent co-occurrence of groups of flexible genes rules out the scenario that they could be forming ecotypes and sustaining the diversity of the core genome, even for the relatively “fast” dynamics and limited persistence of blooms such as C1.

#### Marker genes, metagenomes, and references

The quasi sexual nature of the HLII population suggests that at phylogenetic levels above the tight clusters such as C1, conventional methods for analyzing diversity from metagenome or amplicon data are very problematic. Single genes cannot act as a proxy for whole genomes for typical HLII divergences(*39*), though this has been attempted (*13*, *38*, *40*). ITS divergences above 1% are poor predictors of whole genome divergence, as seen in Figure 2 (d) and Figure S39 (b). In particular, the CN2 cluster identified by Kashtan et al (*13*) based on 2% ITS diameter does not cluster at the whole genome level. However, within CN2, the tight-clusters C1-C5 with ITS diameters *<*1% do cluster at the whole genome level, although not perfectly. Attempts to estimate the number of putative “backbone subpopulations” within HLII from 1381 ITS sequences from the Atlantic have been extrapolated to estimate the existence of thousands (*13*). However, most subpopulations with substantial clonal backbones must be buried deep within the overall diversity: inferring anything about them must rely on a theoretical framework which does not yet exist.

Another consequence of the widespread genomic mixing is that an oft-used way of identifying sub-populations is also highly problematic: metagenome read recruitment to reference genomes cannot be used to infer the abundance of HLII reference strains, as attempted in (*19*, *41*, *42*). To infer putative “strain” abundances reliably would require a high fraction of genomes from a sample to have low enough divergences from one of the references to share significant clonal backbone, comparable to the tight clusters (SM 10. B.). Tens of HLII reference strains exist. However, only 37 of the 167 quasi-random HLII SAGs have a close enough relative in the dataset. If more and more SAGs are sampled, at some level each is sure to have one – or eventually many – relatives close enough to share a clonal backbone. The statistical properties of such intermediate scale structure – at the level of C1 and other observed tight clusters and sub-clusters – is an open question. Investigating this, as well as more information on finer scales of structure such as the cliques in Figure 1, will require sophisticated ways of combining SAG and meta-genomic data. Another route is to focus more on local samples or subsamples, as done by Kashtan et al (*13*, *37*). This raises the general question of geographical structure of the HLII population to which we now turn.

### Geographical population structure and blooms

Although on phylogenetic scales of the whole clade, HLII resembles a quasi-sexual population with seemingly few barriers to genic level mixing, is this true on geographical scales of the whole Earth? Alternatively, are there geographical correlates that might reflect ecological or epistatic differences, or barriers to mixing? As there is not much direct evidence for ecological distinctions within HLII, we investigate geographical differences and similarities among cells sampled from different locations – at least a rough proxy for different environments. The Atlantic and Pacific oceans have different limiting resources – most clearly iron in the Pacific and phosphorous in the Atlantic (*37*, *43*, *44*) – which might be reflected in geographical structure of genomes. The quasi-random cells comprise 40 from BATS near Bermuda in the Atlantic; 60 from HOT, near Hawaii in the Pacific; 42 other Atlantic; and 23 other Pacific cells.

We first look at the tight cluster level. All the deep-clustered cells were from BATS as were the 7 C1-C5 cells in the quasi-random dataset. The closest pair of cells across oceans is 1-2% (depending on the measure) diverged (Figure S47). Thus there is so far no evidence for successful global mixing of HLII *Prochlorococcus* cells faster than 10^4½−5½^ years. Mixing could be inhibited by even small ecological barriers: e.g. if cells from the Atlantic were 1% less fit than the local *Prochlorococcus* population around southern Africa, all Atlantic cells would die out during passage. Yet such tiny growth-rate differences averaged over 10^4^ generations would hardly constitute different “niches”! If the selective disadvantage were ten-fold less, 0.1%, one in 10^4^ cells would survive. If these Atlantic cells were perfectly neutral in the Pacific, their abundance would be too low to observe, but if even a much tinier amount better than average in the Pacific, their descendants could rise to a readily observable fraction before genomic mixing obliterated their distinction from the rest of the population. Alternately, if only some of their *genes* were better than local variants, these could transfer around and prosper. This leads to the question of geographical mixing at genic, rather than whole-cell, level.

Alleles of core genes, as well as presence or absence of flexible genes, were compared for cell pairs from the same versus different oceans, in search of ocean-specific genes or groups of genes which could reflect ocean-specific ecotypes. Distinct patterns of core-gene allele sharing across oceans were found for closely-related versus typically-diverged cells. For each cell in the quasi-random set, the fraction of core genes at which the cell’s closest relative was from the same ocean was computed. The results are consistent with a null in which alleles were randomly assigned across oceans (Figure 4 (a), blue and green curves). However, the cells that were members of a tight cluster had many more core genes at which their closest relative was in the same ocean (Figure 4 (a), red curve). Focusing on genes rather than cells shows that alleles of core genes diverged less than 0.5% were almost always from cells from the same ocean (Figure 4 (b)). Nevertheless, identical alleles of *>*100 core genes are found across oceans (at roughly the same rate at which they are found in the Pacific, although the Atlantic – in which the genomic diversity is somewhat lower overall – is enriched by 2-3 fold for genes with pairs of identical alleles). This implies that there is substantial global mixing at the level of genes in 10^3½−4½^ years, a factor of ten faster than evidence for mixing of whole cells. By 1% divergence, alleles appear randomly mixed between the Atlantic and Pacific. Consistent with this, core gene alleles that match the clonal backbone of C1 are found at similar frequency in Pacific cells as in non-C1 Atlantic cells (Figure S28). Thus C1 alleles have been steadily mixing globally despite the absence of C1 cells found in the Pacific (SM 5. B.). The mixing of genes from C1-C5 into other HLII cells, as stated previously, is quite extensive: about half the HLII cells contained at least one C1-C5 allele.

**Figure 4:**
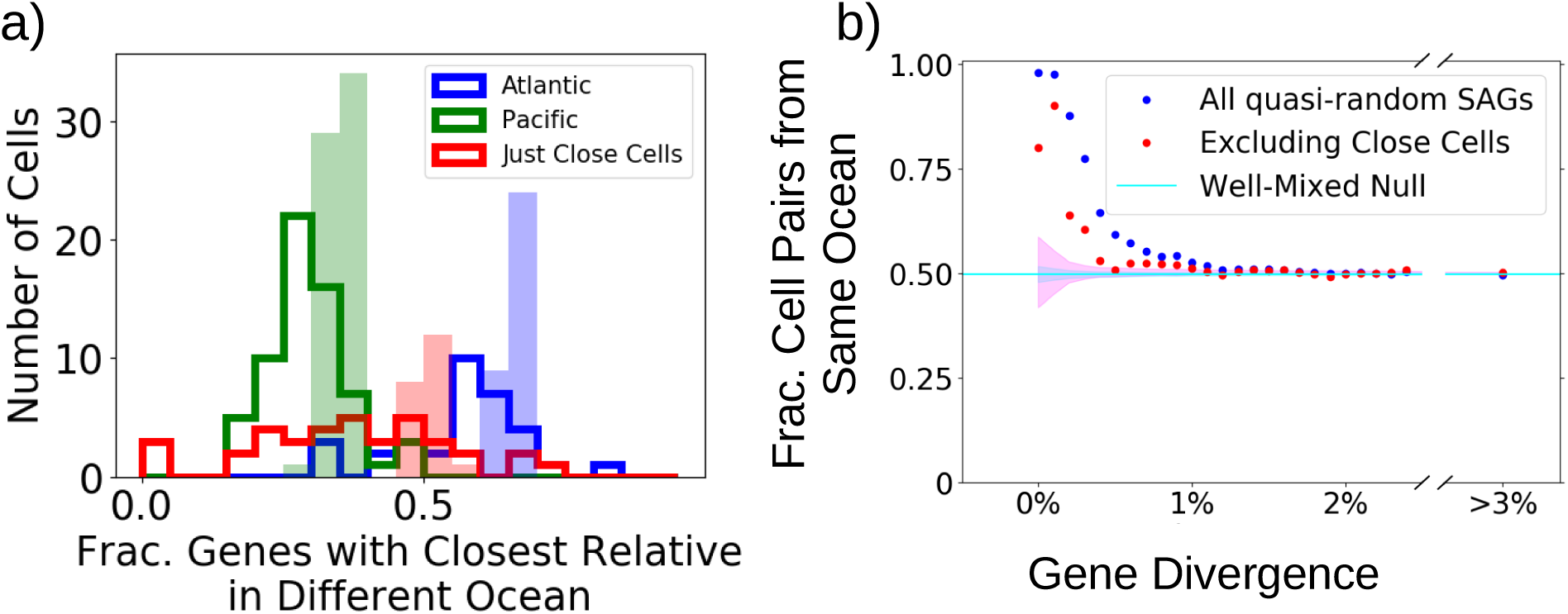
Mixing of genes between Atlantic and Pacific oceans shows little evidence of ocean-specific ecotypes: alleles of core genes mix globally at divergences more than 0.5%. For each core gene in each SAG from the quasi-random dataset, the closest allele from any other SAG from the quasi-random dataset is found and its ocean identified. (a) For each *cell*, the fraction of its core genes with closest relative in the other ocean is histogrammed over cells excluding tight clusters (dark outlined histograms: blue, Atlantic, and green, Pacific) compared to null expectation of sampling uniformly across oceans (light, solid histograms). Contrast with cells in tight clusters (red) showing less cross-ocean genic mixing (SM 9.) (b) For each *gene*, the fraction of cell pairs that are from the same ocean as function of the pair’s nucleotide divergence at that gene, averaged over core genes. All quasi-random cells (blue); excluding close cell-pairs (red). Alleles diverged *<*0.5% are typically from same ocean even though oceans mix ∼ 10^3^ times faster. At higher divergence, data consistent with random mixing (cyan line). Genes less than median length are excluded from the lowest divergence bin (often identical by chance).

In spite of the general mixing of alleles between oceans, we found several anecdotal examples of core genes with ocean-specific alleles (divergences of cell pairs from one ocean significantly lower than inter-ocean cell pairs) – although the localization was incomplete (even in the genes with clearest ocean clustering, at least ∼10 cells from the quasi-random dataset clustered with the “wrong” ocean). The genes with substantially ocean-correlated alleles included iron, phosphate, and phosphatase-related core genes (enriched by factors of 3 to 8), potentially associated with genic adaptations to known environmental distinctions. However, only about 50 core genes have alleles with substantially lower intra-ocean divergence compared to inter-ocean divergence, most of which are not readily ecologically interpretable (Figures S43 and S44).

Flexible genes showed different patterns of inter-ocean diversity than core genes, tending to be more correlated with sampling location. A few percent of flexible genes (mainly in the Atlantic) show evidence of recent gene sweeps (SM 7. B.). At the cell level, cell pairs that are not closely-related at the core genome but share twenty or more flexible genes are almost always from the same ocean (Figure S45). This suggests that in certain locations, some flexible genes have risen to high frequency via horizontal gene transfer – unlinked from any clonal backbone – but do not confer a selective advantage in other locations. This provides evidence for some ecological distinctions that are not correlated with clonal backbones and do not present barriers to horizontal gene flow. However, typical flexible genes appeared randomly distributed across the Atlantic and Pacific oceans (Figure S42). Of the 421 flexible genes with intermediate frequency (found in 4-34 of the quasi-random SAGs), 40 were found exclusively in either the Atlantic or Pacific. These ocean-specific genes were enriched for iron III transport genes 10-fold and phosphate-related genes 6-fold. These overlapped with ocean-specific genes identified in Kashtan 2017 (*37*). Other flexible genes associated with particular oceans also likely reflect adaptations to different environmental conditions(*45*).

#### Asexual blooms

The extensive geographical mixing of genes decreases the overall genomic scale linkage, but there appear to be residual effects of the blooms on timescales over which the whole genomes are overwritten by recombination. At the BATS site, 32 of the 40 quasi-random HLII cells, and all cells in the deep-clustered sample, are in an intermediate-divergence cluster which roughly corresponds to the quasi-equidistant set plus the tight clusters that overlap it (Figure S13). This suggests residual evidence of a persistent sub-population half as old as HLII that rose to high abundance: an *ancient bloom* that spread globally. The genes of this ancient bloom have been generically mixing with other cells ever since, but not quite enough to eliminate all memory of its history, especially in the North Atlantic. More recently, tight clusters such as C1 rose to substantial abundance (and even more recently, tighter cliques within these), and similarly started mixing with other cells. All of these must have initially been primarily asexual blooms and occurred at times that span from the ancient bloom to a thousandth as long ago, as seen in Figure S18 (SM 4. B.). This rough scenario for *some* of the evolutionary dynamics leads us to explore the evolutionary history of HLII by less anecdotal approaches.

### Neutral markers and genetic hitchhiking

Genomic data provides only a snapshot of diversity. To glean information about the underlying evolutionary dynamics, one needs models with clear assumptions and distinguishable predictions. For this, the statistics of synonymous variation are invaluable. Oft-studied is the synonymous site frequency spectrum (SFS), *ρ*(*f*), the distribution over sites of the fraction of cells with a particular variant at the site: its “frequency” *f*. We show (SM 3. B., 3. C.) that synonymous sites are almost all effectively neutral in HLII (Figure S16 and S17). However, there is a large mutational bias making A/T alleles four times as common as G/C alleles at synonymous sites. The polarized SFS of fourfold degenerate sites in Figure 5 (a) shows this asymmetry (although a slight excess of sites with 100% G/C suggest a small fraction of synonymous sites are under selection). What evolutionary dynamics give the SFS this shape? As the gene-level diversity is mixed globally relatively rapidly, treating HLII as well mixed is a reasonable approximation – far more so than for other studied bacterial species – and this enables comparison with theory (*46*).

**Figure 5:**
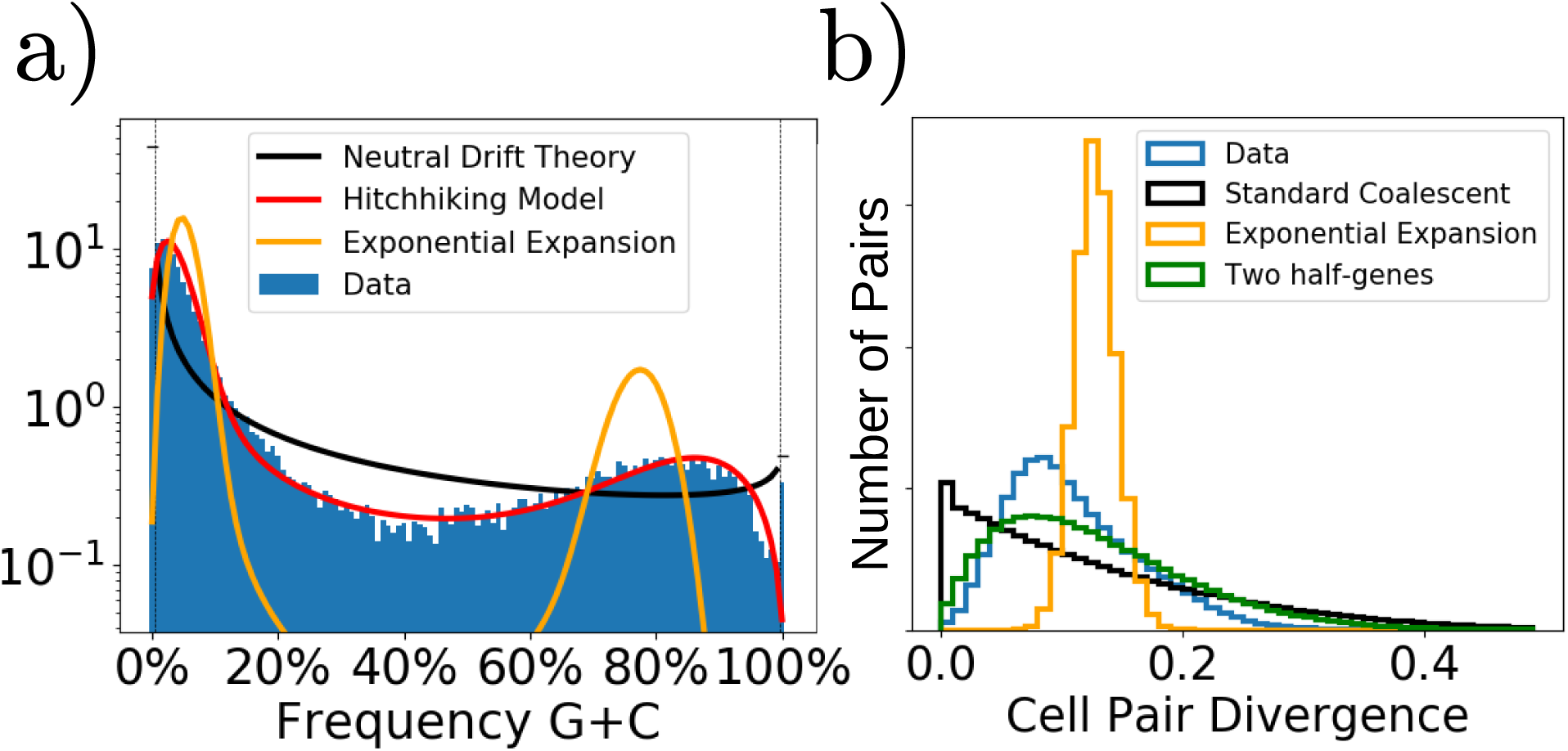
Statistics of synonymous nucleotide diversity compared with models of evolutionary history. (a) Polarized synonymous site frequency spectrum of fourfold degenerate sites, lumping A&T and G&C together and down-sampled to 120 quasi-random cells. Data (blue histogram) compared with: neutral drift-like model (black, with dashed lines delineating monomorphic sites); a uniform expansion (orange); and a hitchhiking model in which neutral sites undergo rapid changes in frequency due to dynamics driven by linkage to selected elements elsewhere on the genome (red). Theory curves all use only the same two parameters: the average G/C frequency and the heterozygosity 2⟨*f_GC_*(1 − *f_GC_*)⟩ which is the probability two cells are different at a site; no other fitting is done. (b) Distribution of cell-pair synonymous genic divergences across all genes and all pairs of quasi-random cells. Data (blue) are inconsistent with a constant probability of coalescence (black) (e.g. from conventional asexual drift) and a single radiative expansion (orange), but similar to a model of two half-genes with independent phylogenetic histories (green), approximating the effects of recombination within genes which causes the observed linkage decay (SM 4. F.).

Demographic fluctuations (drift) are negligible for huge populations: the dynamics of *Prochlorococcus* must be driven by differential temporally-varying selection. However, if selection varies rapidly enough to cause only relative abundance changes of less than 1*/*(sample size)∼ 1%, this would manifest as pseudo-drift. In Figure 5 (a), different evolutionary models for the SFS are compared with the data. The only parameters are the asymmetry of mutation rates (determined by the average frequency ⟨*f_GC_*⟩), and the product of the mean coalescent time (*T_c_*) with the mutation rate, inferred from the heterozygosity (*π* = 2⟨*f_GC_*(1−*f_GC_*)⟩). The pseudo-drift prediction (black) is qualitatively wrong: indeed, drift cannot cause two peaks at intermediate frequencies. An alternate model, a single slow expansion (*47*), gives two peaks (orange) but the Luria-Delbrü ck spectrum is squeezed due to almost-deterministic mutations that occurred after the population was sufficiently large, yielding far too few mostly-G/C sites.

A more promising model is genetic *hitchhiking*. If a synonymous variant is lucky enough to be on a genome that has just acquired a currently beneficial genetic change – a combination of mutations, or a core-gene allele or flexible gene transferred onto its genome – it will rise in abundance. If it is very close to the driving element, it can be transferred around with it and hitchhike to some substantial fraction, *h*, of the population limited by how long it is linked to the transferred segment. The simplest model is local hitchhiking with the probability a site of interest is transferred with a driving element decaying with their separation. This gives a distribution of hitching factors *w*(*h*) ≈ 2*H/h*. With the rate of hitchhiking *H* = 1*/T_c_* and no extra parameters, the predicted SFS in evolutionary steady-state is shown in Figure 5 (a), red curve. It is remarkably close to the data.

In addition to genic-scale hitchhiking that affects only sites within a few kb of the driving element, there are also events in which a whole genome increases in abundance: the largely asexual blooms in HLII must each have resulted from *some* single ancestor being driven to substantial abundance. Ecological or geographical barriers, as well as susceptibility to phages, are likely to limit the magnitude of genome-wide hitchhiking which must persist for substantial time to affect the SFS. The apparent rate of occurrence of genome-wide hitchhiking in the past – conditional upon survival to the present – can be inferred by the number of cells that belong to some tight cluster as a function of the diameter of the cluster. As shown in Figures S18, S14 (SM 4. F.), the *apparent* rate is roughly constant back to 4% divergence. This supports the evolutionary steady-state hypothesis. However, the rate of occurrence of the inferred genome-scale events is less than expected from the inferred overall hitchhiking rate, *H*, and there are too few large events involving a substantial fraction of the population. The dearth of genome-scale hitchhiking is *a priori* surprising. *Genic* scale selected elements that transfer around slowly enough as they rise in abundance would cause the whole genome to hitchhike. Even if quite rare, these could dominate over more rapidly transferring selected elements as they can affect hundreds of times as many sites. That they do not may be due to most primarily-asexual blooms decaying rapidly. In any case, it appears that since the putative ancient bloom, genome-wide hitchhiking likely only contributes a modest rate of small *h* events which do not change the SFS by much except at low frequencies which the limited sample size cannot resolve.

Although the excellent agreement of the simplest hitchhiking model with the observed SFS may be fortuitous, it does strongly suggest that the basic picture of sporadic hitchhiking events drove the evolutionary history of HLII. Crucially, this must have involved many hitchhiking events with *h* much larger than the ≲5% frequency of the observed tight genomic clusters. That most HLII cell pairs are diverged at least 4% suggests that the last genome-wide bloom to reach – and sustain – high abundance occurred long in the past. The dominance since then by undetected hitchhiking events – most likely genic-scale is supported by analysis of the SFS for the group of quasi-equidistant cells that are part of the residue of that ancient bloom. Although tight clusters are excluded from this group so that it contains no direct evidence of genome-scale hitchhiking, again, as shown in Figure S18 (SM 4. B.), its SFS is fit remarkably well by the simple hitchhiking model.

Evolutionary histories on genic scales are reflected in distributions of cell pair genic divergences, which provide complementary information to the SFS. In Figure 5 (b), the distribution of pairwise synonymous divergences across cells and core genes is compared to simulated sampling from several predicted forms. The data show a suppression at low divergences inconsistent with standard asexual coalescence (black curve), but are broader than the distribution for the star-like phylogenies from an exponential expansion (orange curve). The data are more consistent with a model (green curve) in which each gene is comprised of two independent half-genes caricaturing the decay of *intra-gene* linkage correlations (Figures S29-S31)(SM 4. F.).

Hitchhiking affects not only single-site properties, but also linkage between different sites on the genomes. If two sites are both close enough to be transferred around with the driving element, they will both hitchhike to a larger frequency, making them strongly linked. However, if they are somewhat further apart it is more probable for one site to be transferred in a recombination event without the other, resulting in linkage decay. In some bacterial populations (*1*), linkage is observed to decay so fast with separation that this is a likely scenario. However, for HLII there are certainly events that the whole genome can hitchhike on and these will cause genome-scale linkage. Nevertheless, recombination breakpoints of *successful* transfer events appear to have occurred within most core genes often enough to make the linkage decay on genic-scales. It is possible that many of these reflect the ends – varying during a sweep of transferred segments carrying selected elements that drive hitchhiking, rather than close-to-neutral transfers between hitchhiking events. If that is indeed the case, then the dominant processes that unlink synonymous sites within genes and between nearby genes may be due to these same hitchhiking events with the scale for loss of linkage then related to typical lengths of transferred segments.

## Discussion

The analyses we have carried out suggest a plausible scenario for the evolutionary dynamics of *Prochlorococcus* HLII which is illustrated in Figure 6. Occasionally, one bacterium has a genetic change – from homologous recombination, transfer of a flexible gene, or a combination of mutations – that gives its descendants a sufficient selective advantage, averaged over time and over the ever-expanding area of the ocean over which they are advected – to drive them to rise to a substantial fraction of the HLII population. This expansion is limited in abundance or duration, either by ecological and spatiotemporal variations – including susceptibility of its now-large population to phages – or by the genetic element that drove the expansion transferring around to other genomes and driving their expansions at the cost of its original hosts. At least some of these expansions are fast enough relative to homologous recombination – albeit very slow in generations or years – that most of the clonal backbone of the lucky ancestor is driven to substantial abundance in such a largely-asexual bloom. However, during its expansion and for as long as it persists, the cells in the bloom gradually exchange their core genomes – and faster their flexible gene content – with the rest of the HLII population. Thereby, they gradually blend in and eventually become almost unrecognizable. This genetic mixing is both ways: alleles from the bloom continually spread into the rest of the population on global scales even when the bloom is confined geographically.

**Figure 6:**
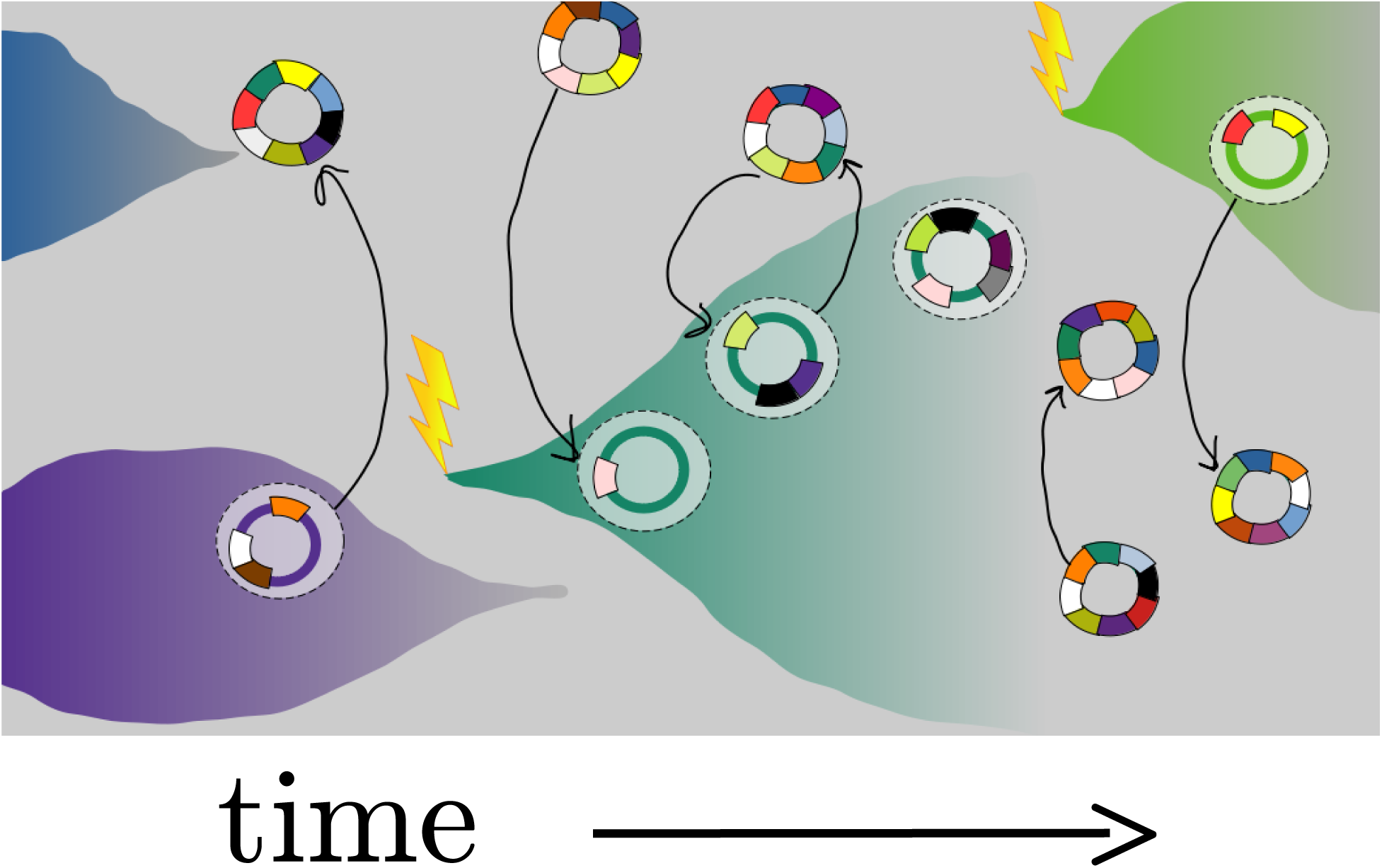
Caricature of evolutionary history of HLII characterized by overlapping selection-driven expansions of genetic variants in primarily-asexual “blooms” (blue, purple, turquoise, and green clones), from within a sea of quasi-uniformly diverged HLII cells (multi-colored genomes). Each bloom begins as a clonal expansion whose descendants share a clonal backbone that rises in abundance over timescales long compared to ocean mixing, limited by subtle ecological variations and transfers of driving genetic elements to other genomes. As blooms expand and persist, they undergo homologous core-genome recombination with cells outside the bloom, steadily eroding their clonal backbones and concomitantly “infecting” other cells with their genomic material; by the time a cell pair’s backbone has diverged ∼0.3% by accumulation of mutations, about 10% of the core genome has been overwritten by recombination and eventually the bloom blends into sea of typical HLII cells.

Tight genomic clusters such as C1-C5 and others in the quasi-random set (SM 2. B.) are the result of largely-asexual blooms, and the cliques within these reflect more recent and more limited blooms leading to the fine-scale structure of whole-genome divergences seen in Figures 2 (c) and S15. Quantitative analysis of the clustering (SM 2. B., 4. F.) is consistent with these asexual blooms having occurred at a roughly constant rate back to ∼4% divergence. The simplest scenario on longer timescales is that the collection of cells that includes the quasi-equidistant subgroup represents a very large and persistent ancient bloom which has been largely overwritten by recombination, but which is still discernible as a peak in the whole genome divergence distribution, especially at the BATS site, and has left behind residual weak linkage along the whole length of the HLII genomes.

The decay of linkage correlations on genic scales (SM 6. A.) and direct evidence of partial genic sweeps on a range of time and magnitude scales (SM 4. F.) suggests that much of the evolutionary dynamics is driven by quasi-sexual processes carrying homologous genetic segments throughout the population. Identifying the driving genetic elements is intrinsically hard because they are obscured by all the other genetic changes taking place, but they leave their hallmark on the statistical properties of the neutral variation, particularly the synonymous variation (SM 3. B.). We have seen that the simplest model of hitchhiking dominated by neutral variants transferred with nearby driving elements leads to a site frequency spectrum that is much closer to the data than either drift-like processes (caused by a multitude of unobservably small selective events) or a single expansion. However, this requires a distribution of magnitudes of hitchhiking events that includes ones large enough to raise allele frequencies to of order half the HLII population. There is no direct evidence for large, persistent genome-wide events more recent than the ancient bloom. However, there is evidence for frequent genic-scale events: many genes show evidence of relatively recent partial sweeps. Yet it is at this point unclear whether these are frequent enough, large enough, and spread out enough in time to account for the hypothesized dominance of hitchhiking by events that directly transfer synonymous variants that are near the driving genetic elements. To test this scenario requires further theoretical developments which includes – crucially – the spatiotemporal ecological variations which must drive the selective processes but prevent any genes – let alone genomes – from taking over the population.

The continual genetic mixing into and out of the descendants of blooms is evidence that blooms do not represent sexually-isolated “ecotypes” or other such groupings. Although their existence implies some subtle ecological differences originally, these may no longer be present in cells whose genomic backbones are only ghosts of the bloom or which have migrated to another ocean. By contrast, at higher phylogenetic levels there is little genetic mixing. Occasionally, core genomic material transferred from outside HLII is found, but evidence of successful inter-clade transfers is rare. The apparent gene flow barrier between the clades of *Prochlorococcus* means it is appropriate to view HLI and HLII as distinct species.

The prevalence in the Pacific ocean of genes from cells only found in the Atlantic shows that genes, even if not cells, regularly make it from the Atlantic to the Pacific, albeit perhaps a thousand-fold more slowly than surface-ocean mixing. The rate of geographic spread of genes is likely set by selective pressures rather than geophysical mixing constraints. Genic variants that are on average beneficial under a broad range of conditions, or in both oceans, can spread faster than typical ocean mixing times. Conversely, genes that are only beneficial in some locations mix – if at all – much more slowly than ocean mixing. This, together with recombination unlinking genes, results in inter-ocean diversity patterns that vary from gene to gene. Very weak ecological barriers could help localize more-closely-related strains; some tight clusters, such as C1, are only found in a single ocean in this dataset – although might also be found elsewhere with more sampling. For example, about half of closely-related pairs in the quasirandom dataset have one cell in each of the Atlantic and Pacific. Flexible genes turnover much faster than core genes, but still much slower than ocean mixing times, implying that differences in their distribution across oceans are due to differing selection or to barriers to spreading: they could be generally deleterious in some conditions or do worse in genomic backgrounds that prevail in some locations.

### Open questions and prospects

While the rough scenario we have laid out appears to be a plausible description of the evolutionary history of HLII, it is certainly only a very partial one. Within the framework of genetic hitchhiking – which certainly plays a large role – one can work out many more statistical properties, such as linkage correlations of various forms. Although this typically requires some extra assumptions and choices of parameters, there is enough information in, for example, joint distributions, to try to nail some of these down. A challenge is to understand which predictions are relatively robust to more detailed assumptions, and which can best distinguish between scenarios. For example, is it possible that the synonymous site frequency spectra, and other properties that we have argued support a scenario of continual genic-scale hitchhiking, can be explained by a very different scenario? Could enough genome-wide hitchhiking events whose descendants have died away from changing selection leave similar statistical properties? Alternately, could the gradual genetic blending of the ancient bloom with the rest of the HLII population result in quantitatively similar features?

A major puzzle that is not at all addressed by our analyses is the extremely broad range of time scales: how can extensive diversity without apparent niche structure persist for of order a million years – 10^8−9^ generations? The shortest time scale genomic data have provided evidence for is hundreds of years. Even relatively “fast” blooms lasted for 10^4−5^ years, and it takes 10^5−6^ years for genomes to be overwritten by recombination. In the meantime, selection pressures can act on the whole genome as well as individual genes. By contrast, in the laboratory *Prochlorococcus* evolution is quite fast: without even trying to exert selective pressure, some genetic changes take over in a few years implying a few percent fitness difference per generation (for example, in SS120, SS35, SS51, and SS2, four clonal derivatives of the strain LG (*48*)). Phage variants have very different efficacies in killing *Prochlorococcus* variants (*49*, *50*) which likely result in large variations in death rates in the ocean. Direct measurements of dynamics in the ocean are hard to interpret because of the turbulent advection, but Kashtan et al (*13*) observed clear seasonal variations in relative abundance of the C1-C5 clades at BATS. With such fast processes occurring, why don’t a small number of HLII variants, with sufficient ecological differences to stably coexist, take over much faster than the apparently very slow blooms?

What are the dominant forces driving selection? Might physical or chemical variations be sufficient? Comparing the Atlantic vs Pacific Oceans is one crude avenue to find discrete ecological niches, but few core gene alleles, and only a small fraction of the rapidly overturning flexible genes, are associated with one ocean over the other. Some correlations of gene content with local temperature have been found (*7*, *38*), but the effects are likely too weak to sustain putative ecotypes with the rapid ocean mixing. A potential mechanism for generating and maintaining diversity comes from the ocean turbulence itself. The descendants of an individual cell are spread around randomly over about a kilometer in a day. The number of other *Prochlorococcus* by which these descendants are potentially affected in one day is thus ∼ 10^19^, although they will only interact indirectly being affected by some average property of these many cells. However, cells can potentially interact directly with a comparable number of phages. Thus, each cell feels a different temporally varying environment, and spatiotemporal chaos means that no two feel exactly the same variations in their environments. If a variant were worse than another in all circumstances, it would certainly die out. To have a chance at surviving, variants must be better in *some* circumstances. If a variant is worse than another on *average*, one might expect that it would die out, but that is not correct. If the variant is somewhat worse on average but subtly different enough that the spatio-temporally chaotic environment it feels is sufficiently out of sync with the environment that a competitor experiences, then both the variant and its competitor can persist. It has been shown that this effect can, in simple models, enable subtly different strains to persist – in the absence of evolution – for times that are exponentially long in the spatial extent of their habitat (*51*, *52*). This scenario is most easily realized if the dominant differences between strains are how they are affected by a diversity of slightly different phages for which “kill-the-winner” dynamics can give rise to spatiotemporal ecological chaos. It is plausible – but not demonstrated – that similar effects could occur if the differences between genetic variants are primarily responses to, and impacts on, varying chemicals in the environment. Whether this scenario is quantitatively plausible for *Prochlorococcus* in the ocean, and whether it is robust under long term evolution, are questions for future research.

To advance understanding of *Prochlorococcus* diversity and evolution, it is essential to make use of far more data then we have done – including existing metagenomic data and more SAGS – and to figure out what further data would be most informative for distinguishing between scenarios. For example: at greater sampling depth, is “everything everywhere” at the tight cluster level? At the genic variant level? How deeply does one have to sample to find cells – or parts of genomes of cells – that are far closer related than the closest thus far? For this, short read metagenomic data is not useful, as it can only distinguish genomic material substantially more than 10^−3^ diverged. Very long read metagenomic data (≫10kb), if low enough error rate and deep enough, might provide much more information. If genomic regions that are sufficiently long and variable can be sequenced efficiently enough to identify very closely-related cells to then fully sequence individually, that might be the most promising route.

Time course data cannot directly provide insight into the rates of evolutionary processes in *Prochlorococcus*, and even obtaining information on rates of ecological processes is challenging due to the ocean mixing. However, correlations between abundances of *Prochlorococcus* genetic clusters and co-occurring bacteria and – especially – phages, sampled from many spatial locations and times, could reveal interpretable dynamical processes (*53*). For this, as well as more broadly for any extensive sequencing projects, theoretical frameworks to guide sampling, sequencing strategies, and – most crucially – predictive analyses, are mandated.

## Acknowledgments

We thank Paul Berube, Penny Chisholm, and Devaki Bhaya for useful discussions. This work was supported in part: by the Simons Foundation via a Postdoctoral Fellowship 730295 to GB and Sabbatical Fellowship to DSF; by NSF Grants PHY-1607606 and PHY-2210386 to DSF; by NSF GRFP, Stanford Graduate Fellowship, and CEHG Fellowship to AP. Data analysis was partly performed on the Sherlock cluster operated by the Stanford Research Computing Center.

## Supplementary Materials

### 1. Data Processing

In this section we describe the data and processing pipeline used in our study. The section has two parts. The first part describes the data we used, including sampling locations and times, and our estimates of sequencing error rates. This part also contains the definitions of the *deep clustered* and *quasirandom* datasets used in the main text. The second part describes the constructions of core and flexible gene groups used in the later analysis.

#### A. SAG selection, assembly statistics, and error rates

##### A.1. SAG selection

We use 96 SAGs from Kashtan et al.’s 2014 dataset – which we term the *deep clustered* dataset – all of which were sampled near Bermuda in the North Atlantic at 60 m depth over 3 sampling times which together spanned 5 months (November 2008, February 2009, April 2009) (*2*). Within this dataset, 90 SAGs are selected at random from a cluster of the High-Light II (HLII) clade called “CN2”, which Kashtan et al. define based on a 2% ITS divergence diameter. Given membership in this ITS cluster, SAGs are selected randomly. CN2 is divided into 5 subclusters, labeled “CN2-C1” through “CN2-C5,” defined using finer-scale clustering of ITS divergence. The largest of these subclusters, CN2-C1, contains 51 SAGs from the deep clustered dataset (2 additional CN2-C1 SAGs are excluded because they failed to be processed by IMG). We refer to this group of SAGs as *the C1 cluster* throughout. We find that two of the 51 SAGs within C1 are relative outliers at the whole genome scale (Figure S2). The two outlier SAGs are not included in Figure 1 or Figure 2 (c) in the main text. We will sometimes denote the remaining 49 SAGs as *C1 without outliers*.

We used, in addition, 167 HLII SAGs from Berube et al.’s 2018 dataset, which we term the *quasi-random* dataset (*1*). The SAGs in this dataset are chosen for sequencing based on one of three criteria:

1. Fast whole genome amplification, which correlates with good genome recovery
2. Presence or absence of the narB marker gene
3. ITS sequence, aimed at sampling diverse Prochlorococcus ecotypes

The cells chosen for fast amplification form a quasi-random sample – they are expected to not be biased by genome content or phylogeny, apart from perhaps higher AT content, which is not very variable within HLII (*3*) Figure S1 shows that cells selected based on the presence/absence of narB are distributed randomly across the entire diversity of HLII found in this sample. Figure S3 shows the ITS and whole genome divergence distributions of cells selected via the three methods above and, finding them to be remarkably similar. Because cells selected via the narB and ITS methods are similarly distributed to cells selected via whole genome amplification, we treat the entire HLII sample from the Berube et al. 2018 SAGs as quasi-randomly selected from the diversity of HLII. We use the genes.fasta files for these SAGs, processed by IMG, which list the ORFs identified in the assembled contigs of each SAG.

**Figure S1:**
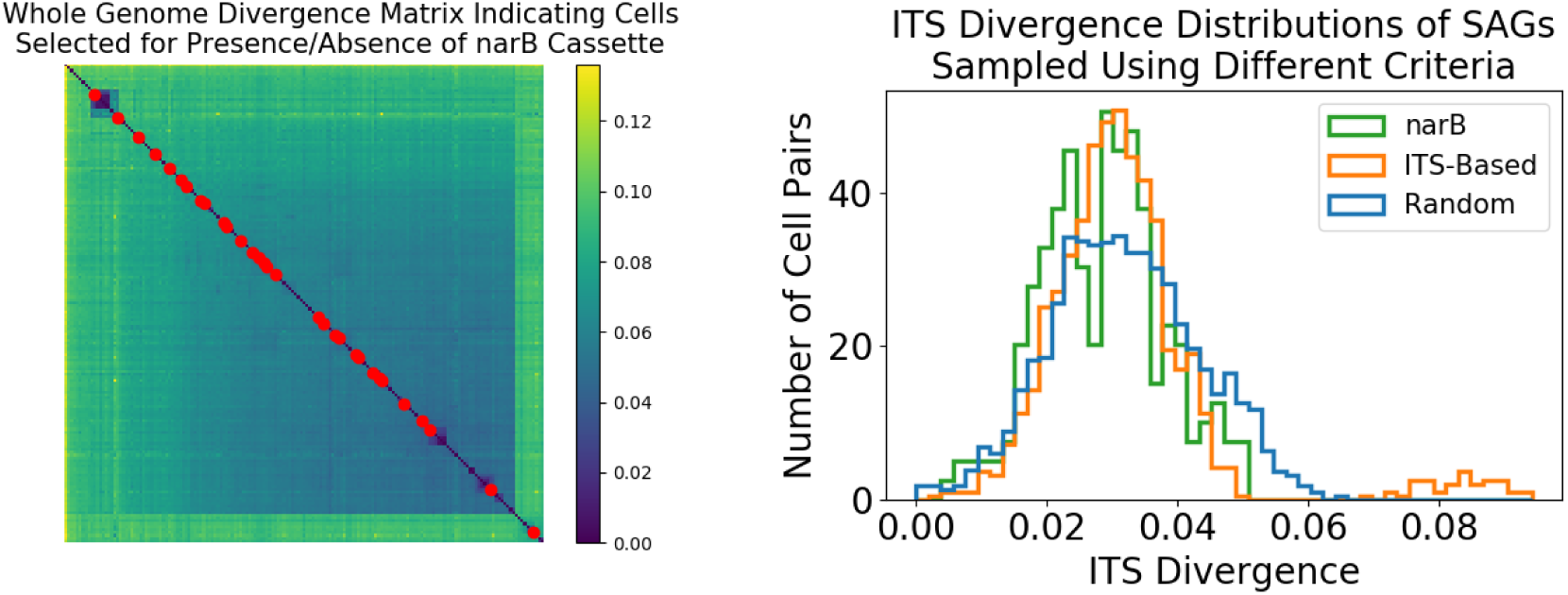
The 167 HII SAGs from the Berube et al. 2018 dataset can be treated as a quasi-random sample of HLII. **(Left)** Core genome divergence matrix of all 167 SAGs, with narB-selected SAGs shown by red dots along the diagonal. Divergences were calculated as explained in B.. Note that the red dots span the diversity of HLII and are not clustered. **(Right)** The distribution of ITS divergences of all 167 SAGs grouped by the three selection criteria used in (*1*): presence of narB gene (green), ITS sequencing divergence (orange), and fast whole-genome amplification (blue).

**Figure S2:**
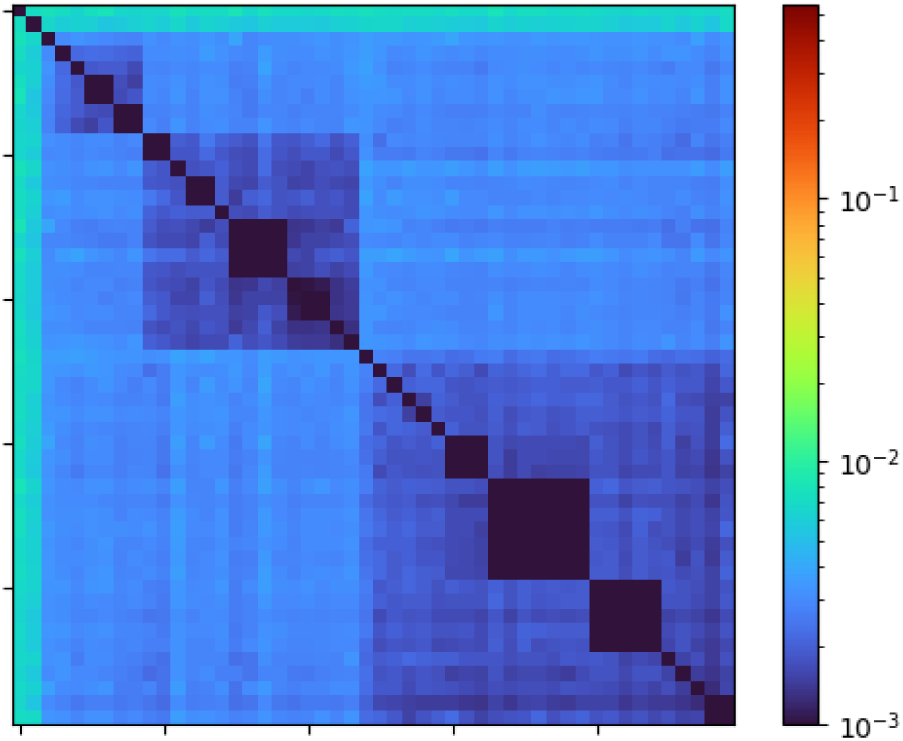
Median gene nucleotide divergence distance matrix of C1 SAGs from the deep clustered dataset, here including two outlier cells which cluster with C1 at the ITS level but not at the whole genome level. These SAGs were excluded from Figure 1 and Figure 2 (c) in the main text.

**Figure S3:**
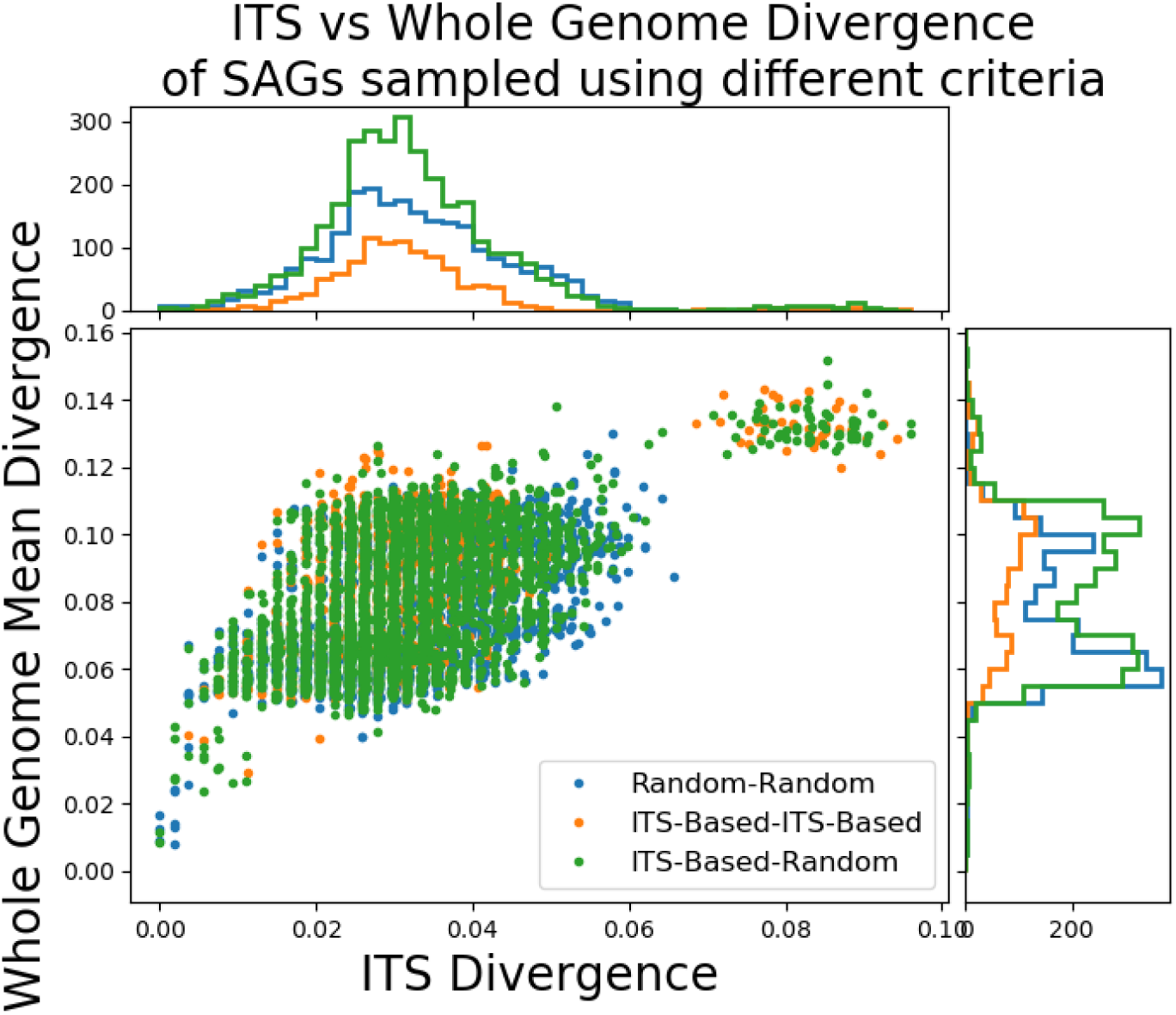
Comparison of ITS and whole-genome divergences of SAGs selected based on fast whole-genome amplification and ITS divergences from (*1*). Each point represents a pair of SAGs, with the three combinations of selection criteria shown in different colors. Marginal distributions of the ITS and whole-genome divergences for each group are shown in the top and right panels, respectively.

##### A.2. Forms of data

The SAGs we used were sequenced using Illumina and are available in multiple forms. They have been processed by Integrated Microbial Genomes (IMG), with each SAG presented as a FASTA file containing ORFs found on assembled contigs. IMG is a system for analyzing publicly available draft and complete genomes, available through the U.S. Department of Energy - Joint Genome Institute (DOE-JGI). IMG facilitates comparative analysis of genomes, in part by providing functionally annotated open reading frames (ORFs) from sequencing data (such as contigs) (*4*). In addition, Kashtan et al. construct a “C1 Composite reference” using 12 contigs from various SAGs in this sample, and then align the reads of each SAG to this constructed template. These are whole-genome alignments. We use both of these forms of data in the analysis.

##### A.3. HLII reference genomes in the context of the quasi-random sample of SAGs

Figure S4 shows the whole genome distance matrix of the 167 HLII SAGs from the quasi-random dataset together with 17 HLII reference genomes obtained from isolate strains. The 17 HLII reference genomes are commonly used for metagenome recruitment (*5*) and span the diversity of the SAGs. Some of the reference genomes occasionally form clusters with each other, but only very rarely cluster with the SAGs.

**Figure S4:**
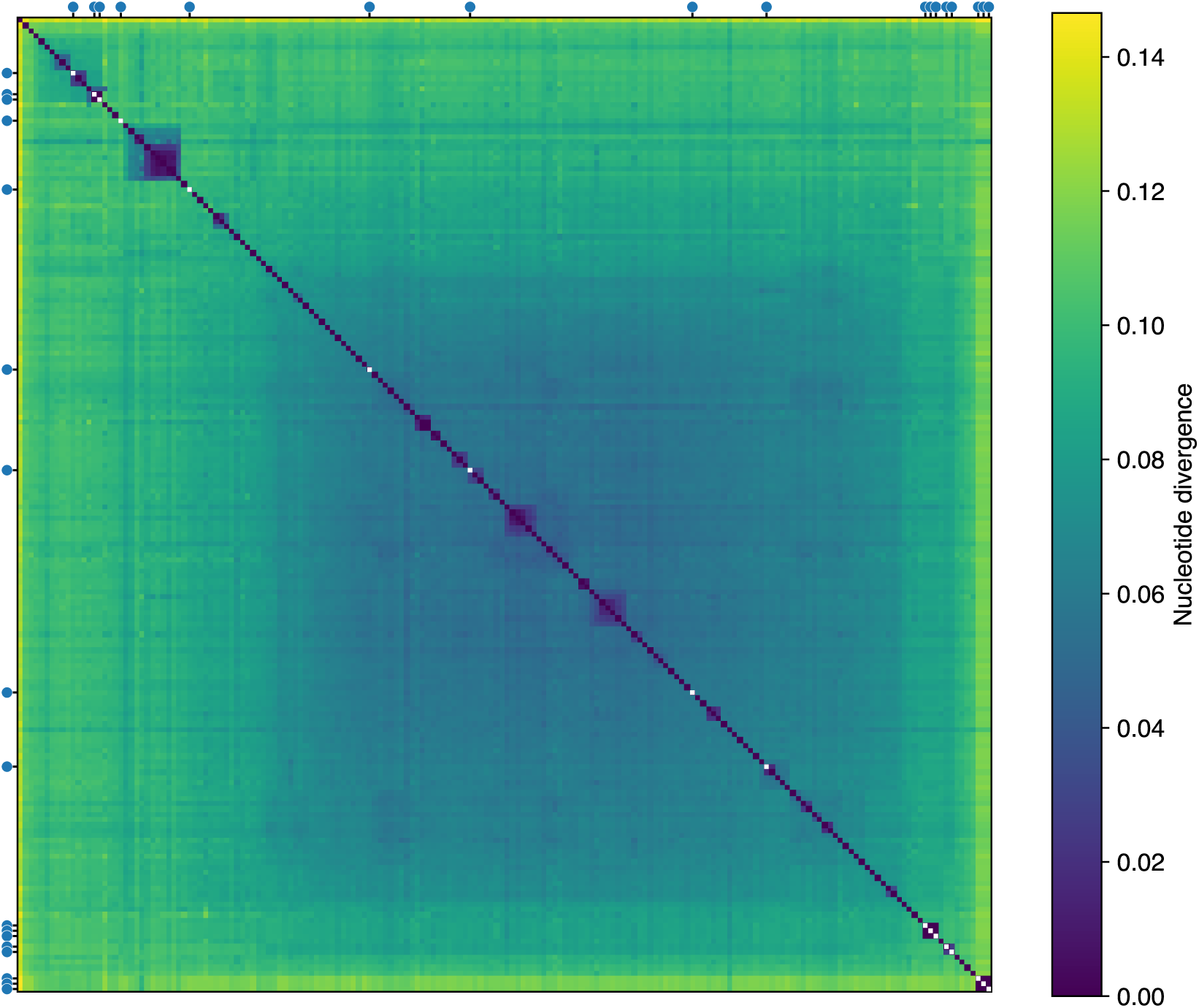
Whole-genome divergence matrix for 167 random sampled SAGs and 17 HLII isolate genomes. Tick marks and white dots on the main diagonal indicate the isolate genomes.

##### A.4. SAG sampling locations

The quasi-random HLII dataset is drawn from around the globe (Figure S5). It comprises 167 HLII SAGs. One hundred of these SAGs come from two sites that are part of long-term ocean monitoring studies: the Bermuda-Atlantic Time-series Study (BATS) in the North Atlantic, and the Hawaii Ocean Time-series (HOT) study in the North Pacific. Of these, 40 are from the BATS sampling site and 60 are from the HOT sampling site. In total, 54 SAGs are from the Atlantic Ocean, including the 40 BATS SAGs and SAGs from 5 additional sites. There are 63 SAGs from the North Pacific, including the 60 from the HOT sampling site. An additional 20 SAGs are from the South Pacific, across 5 different sites. All SAGs from the deep clustered dataset are from the BATS sampling site.

**Figure S5:**
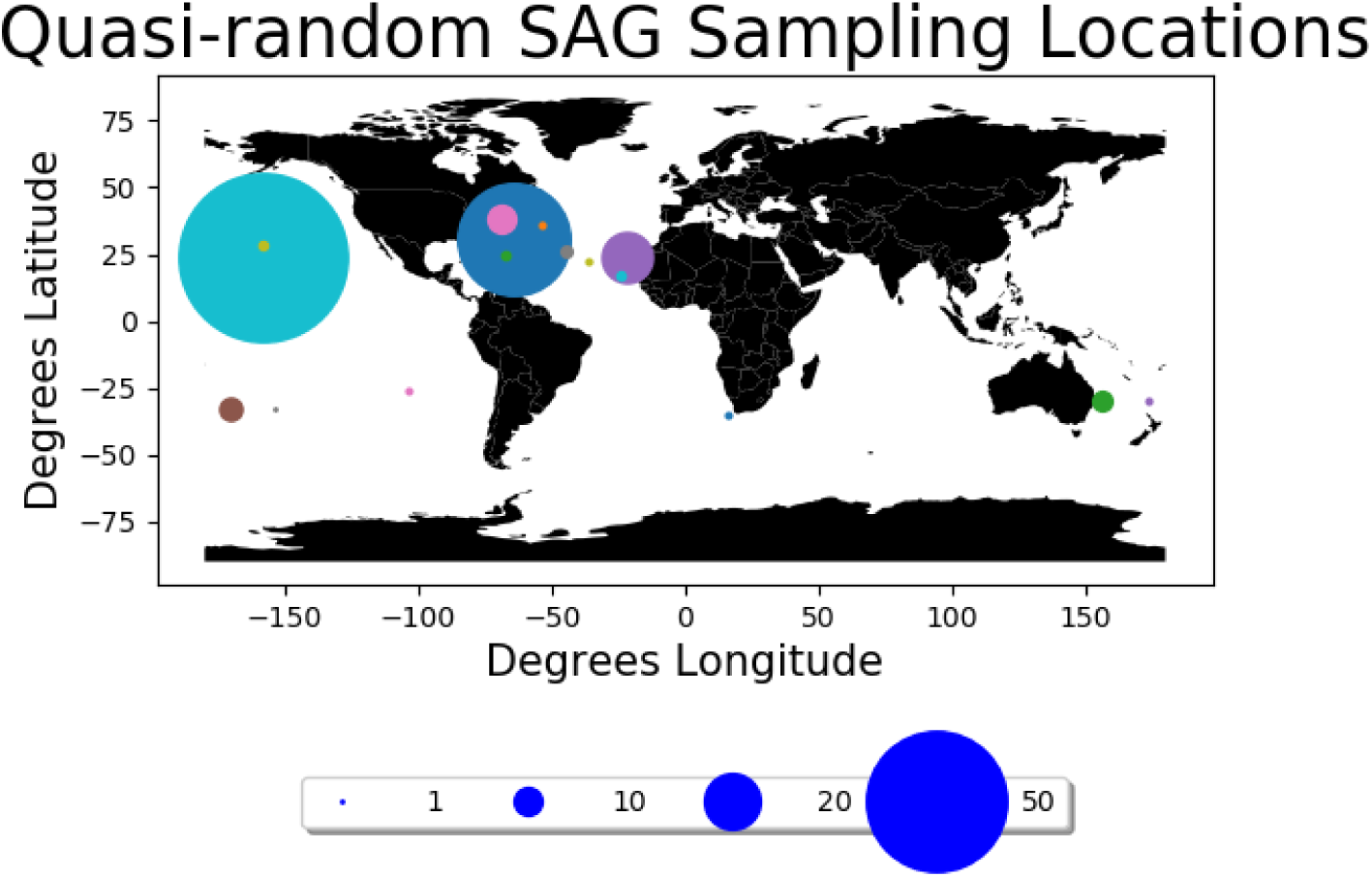
Sampling locations of 167 quasi-random HLII SAGs. The large turquoise point in the Pacific ocean is the HOT sampling site near Hawaii (60 SAGs) and the dark blue point in the Atlantic is the BATS sampling site near Bermuda (40 SAGs). The Westerlies Gulf Stream Province site, close to BATS, is shown in pink.

##### A.5. SAG seasonality

The deep clustered dataset contains 6 close *cliques* of 3-7 cells which are identical at more than half of their genes. All of these cliques contain cells from all three seasons sampled. The closest pairs are consistent with being randomly drawn across samples: 11 of the 40 closest pairs were from the same season compared to a null expectation of <u>^1^</u> of pairs in the same season.

##### A.6. SAG ORFs and Contigs Statistics

SAGs from the quasi-random dataset have on average 44 contigs (mean length 28 kb, median length 11 kb); SAGs from the deep clustered dataset have on average 184 contigs (mean length 6.5 kb, median length 485 bp). Both datasets have similar total lengths of ORFs per SAG. ORFs in the quasi-random dataset have a mean length of 764 bp, median length 633 bp, with a mean of 1600 ORFs per SAG. ORFs in the deep clustered dataset have a median length of 705 bp, median length of 505bp, with a mean of 1558 ORFs per SAG. Much shorter contigs were included in the deep clustered dataset than in the quasi-random dataset, which could explain the difference. Figure S6 shows the distributions of the fraction of C1 and HLII cells covered per gene, for HLII core genes. Cells typically had 50-90% coverage, with 70% coverage on average.

**Figure S6:**
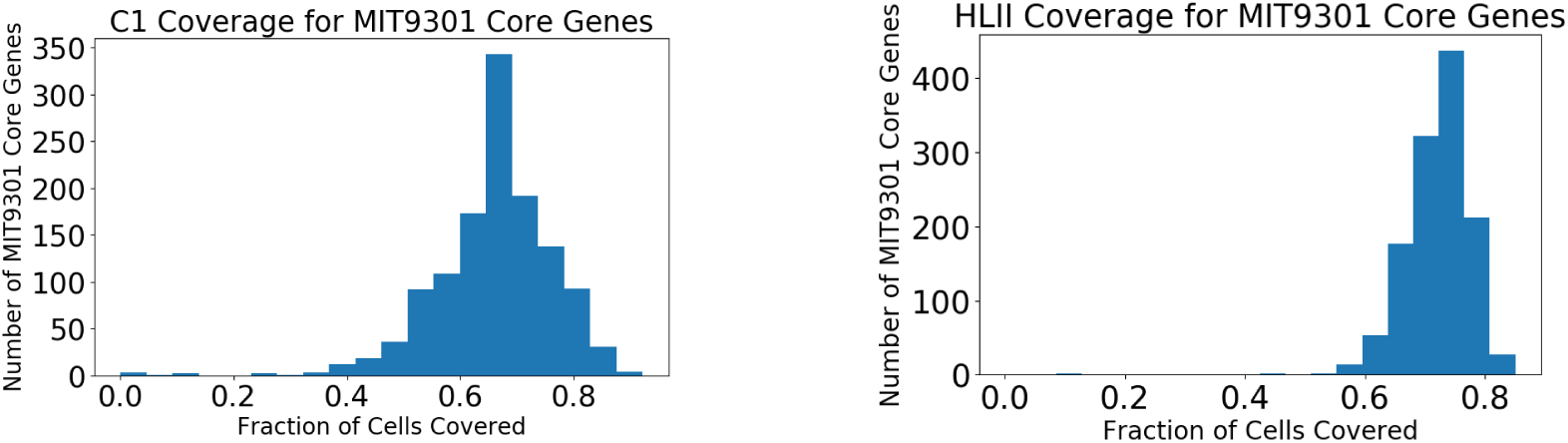
The fraction of SAGs covered at HLII core genes, using our core orthologous gene group construction methods described above, are similar to those reported by the data sources (*1*, *2*). Histograms represent the fraction of cells covered across all 1253 identified HLII core genes. **(Left)** Data for C1 SAGs (deep clustered dataset). The mean fraction of cells covered is 0.66, and the median is 0.67. **(Right)** Data for quasi-random dataset. The mean fraction of cells covered is 0.72, and the median is 0.73.

##### A.7. Estimating error rates

Errors in the reported sequences of SAGs can result from amplification errors, sequencing errors, or other factors. Errors would produce singleton SNPs at some sites where the sample is in reality monomorphic. Consider an error rate *ɛ*, accounting for the sum effect of errors from amplification and sequencing. If *n*_0_ sites are monomorphic in the population and the sample contains *N* covered cells on average per site, then the number of singletons *n*_1*,ɛ*_ produced by errors would be approximately

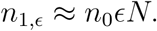

The numbers of monomorphic sites and singletons at nonsynonymous sites in the quasi-random HLII dataset are shown in Table S1. Even if all singletons in the dataset are due to errors, this gives *ɛ* ≲ 10^−3^ (using 120 cells on average covered per site). We use nonsynonymous sites to compute this statistic for the quasi-random sample because synonymous sites have undergone so much mutation in the history of the sample that singletons outnumber monomorphic sites. At the G/C end of the synonymous SFS, there are more monomorphic sites than singletons, but using nonsynonymous sites, which are more likely to be under (negative) selection than synonymous sites, sets a stronger error bound. The corresponding numbers for the C1 cells are shown in Table S2, which provides even tighter bounds on the error rate. The average number of C1 cells covered per site is about 36, giving *ɛ* ≲ 10^−3½^.

**Table S1:**
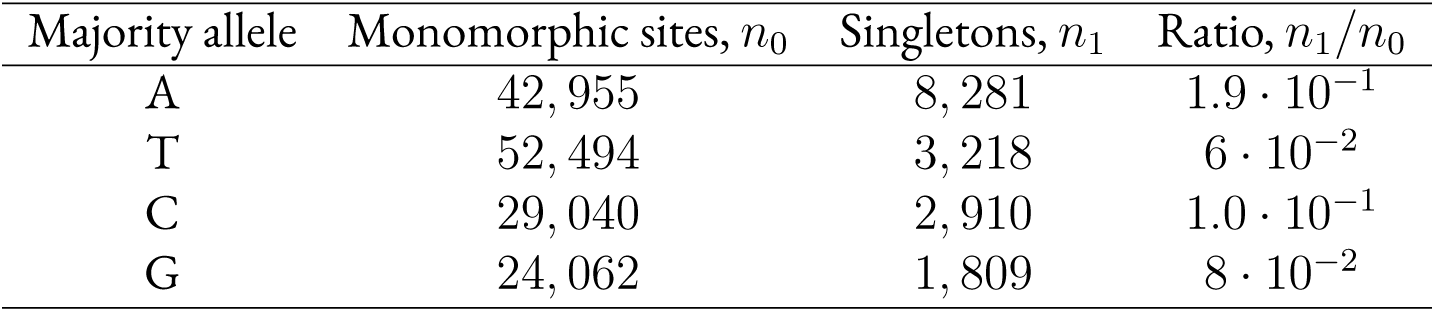
Number of monomorphic sites and sites with singleton mutations at nonsynonymous sites in the quasi-random HLII dataset.

**Table S2:**
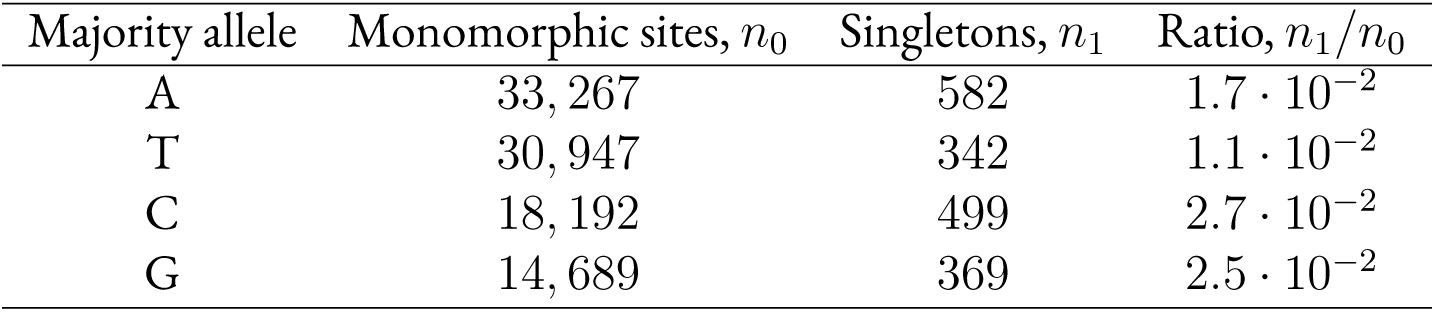
Number of monomorphic sites and sites with singleton mutations at nonsynonymous sites in C1 cells.

The tightest limit on error rates is given by the lengths of identical stretches of DNA of cells from the close cliques. The most closely-related pairs of cells have stretches of identical DNA up to ∼ 10^4½^ bp, the existence of which requires that the error rate be less than 10^−4½^. Note that this estimate is still higher than the 10^−5^ error rate of estimated experimentally by de Bourcy et al. in *E. coli* (*6*). Error rates at this level will not affect any of the results or analysis done here.

#### B. Construction of core and flexible orthologous gene groups

##### B.1. Construction of orthologous gene groups for HLII core genes

To construct clusters of HLII core orthologous genes for cells in the quasi-random dataset and deep clustered dataset, we scaffolded to the reference isolate strain MIT9301 because this strain is closely-related to the C1 group identified by Kashtan et al. (*2*), at the ITS level. We chose to scaffold to a reference genome in order to easily gauge the separation of core orthologous gene groups along the genome. We used reference genomes to verify that gene order is maintained across large genomic distances. We found the gene order in HLII genomes was nearly identical and was well conserved even across clades (Figure S7).

**Figure S7:**
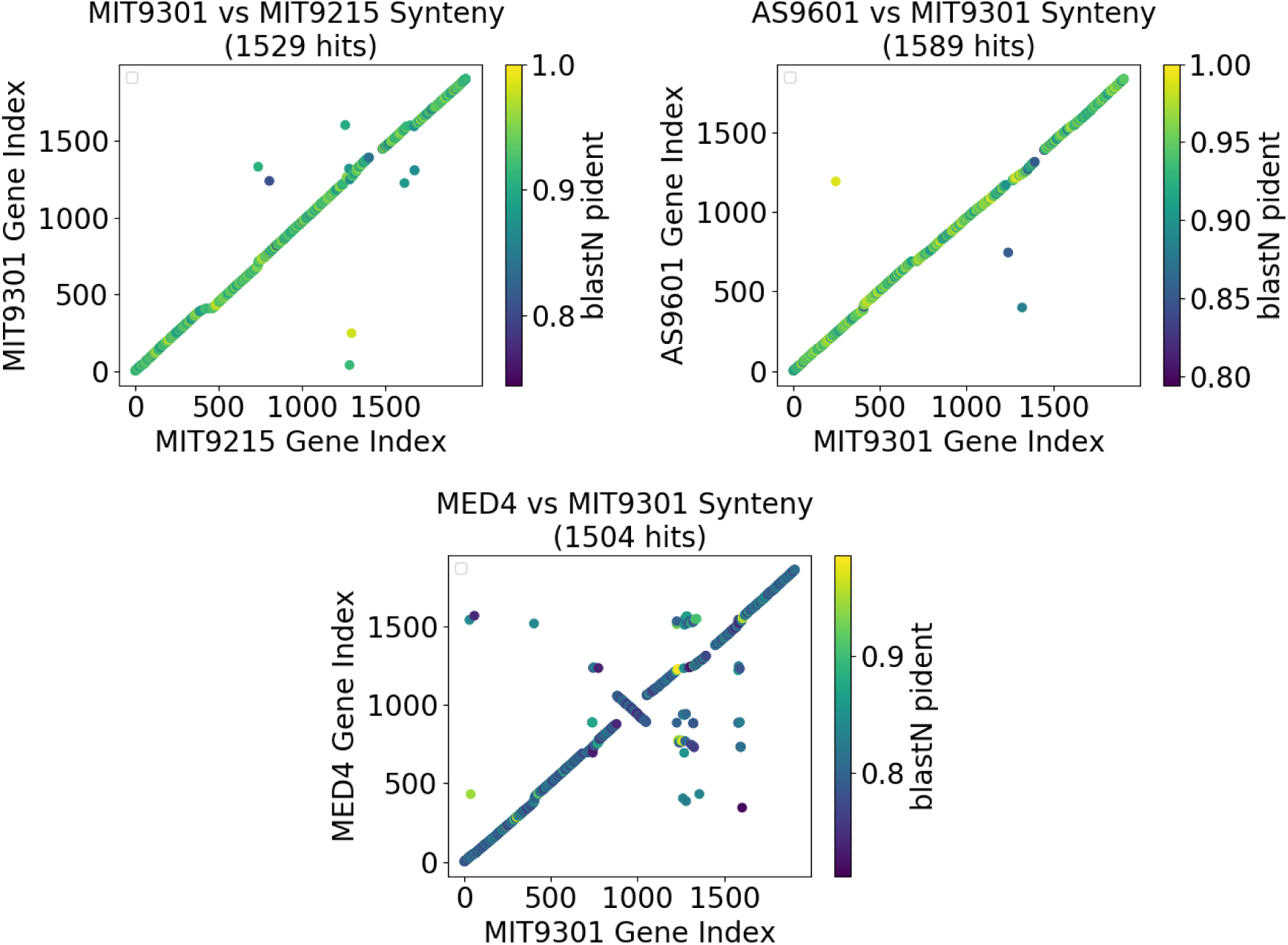
*Prochlorococcus* genomes are highly syntenic. The different panels compare the gene order in one of the HLII reference genomes (MIT9301) with three other references. Top panels show comparisons to MIT9215 (**top left**) and AS9601 (**top right**), both of which are part of HLII. **Bottom** panel shows comparison to the LL reference MED4. In all panels, individual homologous genes are shown as points colored according to the nucleotide identity as reported by BLAST. Only reciprocal best hits are shown.

We started with the IMG FASTA files containing ORFs for each SAG in the Kashtan dataset and each HLII SAG in the quasi-random dataset. We then used BLAST (*7*) to align the ORFs of each SAG to the MIT9301 reference genome. BLAST reported hits as diverged as 40% at the nucleotide level and beyond, while within HLII core alleles are typically within 10% nucleotide divergence or less; therefore, BLASTing to a single HLII reference genome is sufficient to align even highly-diverged alleles at HLII core genes. We identified the reciprocal best hit for each gene in the MIT9301 reference genome in each SAG (if such a hit existed and was covered). We constructed orthologous gene groups for each MIT9301 gene by compiling the best hit ORF from each SAG with a hit. Each orthologous gene group then underwent multiple sequence alignment using mafft (default parameters) (*8*). Finally, we imposed a length cutoff, requiring that each ORF of each orthologous gene group be within 10% length, in bp, from the median nucleotide length for its orthologous gene group.

We checked the coverage for each core gene (see Section B.2. for the construction of the HLII core genome). By definition, each core gene should be present in each HLII cell, and any absences of core genes in SAGs should be due to coverage lapses alone. We checked for duplicate core genes by BLAST-ing `(-qcov_hsp_perc 50)` all core genes of the MIT9301 genome against themselves. We found that eight HLII core genes produced a hit with another (often consecutive) gene on the genome, but that the secondary hit was not a core gene. Only one example of a HLII core gene producing a hit with another core gene was found. Thus, we do not expect duplicate core genes to interfere with our constructure core orthologous gene groups. We found an average of 70% coverage per HLII core gene among HLII SAGs in the quasi-random dataset. Berube et al. estimate a 73.7% average whole genome recovery for HLII cells (*1*). Part of the difference in our observed coverage (70% per gene (Figure S6) vs Berube et al.’s reported coverage is that we require complete genes in order to count a gene as ‘covered’ in a particular SAG; genes cut off by the ends of contigs may not be counted.

##### B.2. Identification of HLII Core Genome

To construct a HLII core genome, we chose five fully-assembled HLII reference genomes roughly spanning the diversity of the HLII clade, which included the MIT9301 reference genome used for creating orthologous gene groups (Section B.1.). The 5 references were AS9601, MIT9312, MIT9215, MIT0604, and MIT9301. Each reference was BLASTed against MIT9301. Only MIT9301 genes with reciprocal best hits in all four other reference genomes were used to build the MIT9301 core. Hit criteria were 80% coverage and 80% nucleotide identity. This resulted in a core genome comprising 1253 MIT9301 ORFs, of the 1907 ORFs identified on the genome. The results are relatively robust to reasonable choices in coverage and nucleotide identity. Note that requiring instead only that at least 3 of the 4 other HLII reference genomes have a reciprocal best hit with a given MIT9301 gene resulted in 1505 genes in the core. We used the stricter cutoff.

The distribution of orthologous gene group sizes has a peak at high coverage even for flexible genes (Figure S8). This is due to core-like genes that have not been included in our construction of the HLII core because they are not found in at least one of the five HLII reference genomes used to construct the HLII core (Section B.2.). When analyzing flexible genes, we restrict the analysis to orthologous gene groups found in 34 of the quasi-random HLII SAGs or fewer (≲ 30% coverage) to exclude these core-like genes. The cutoff roughly corresponds to the lower bound of the trough in the flexible gene histogram from Figure S8.

**Figure S8:**
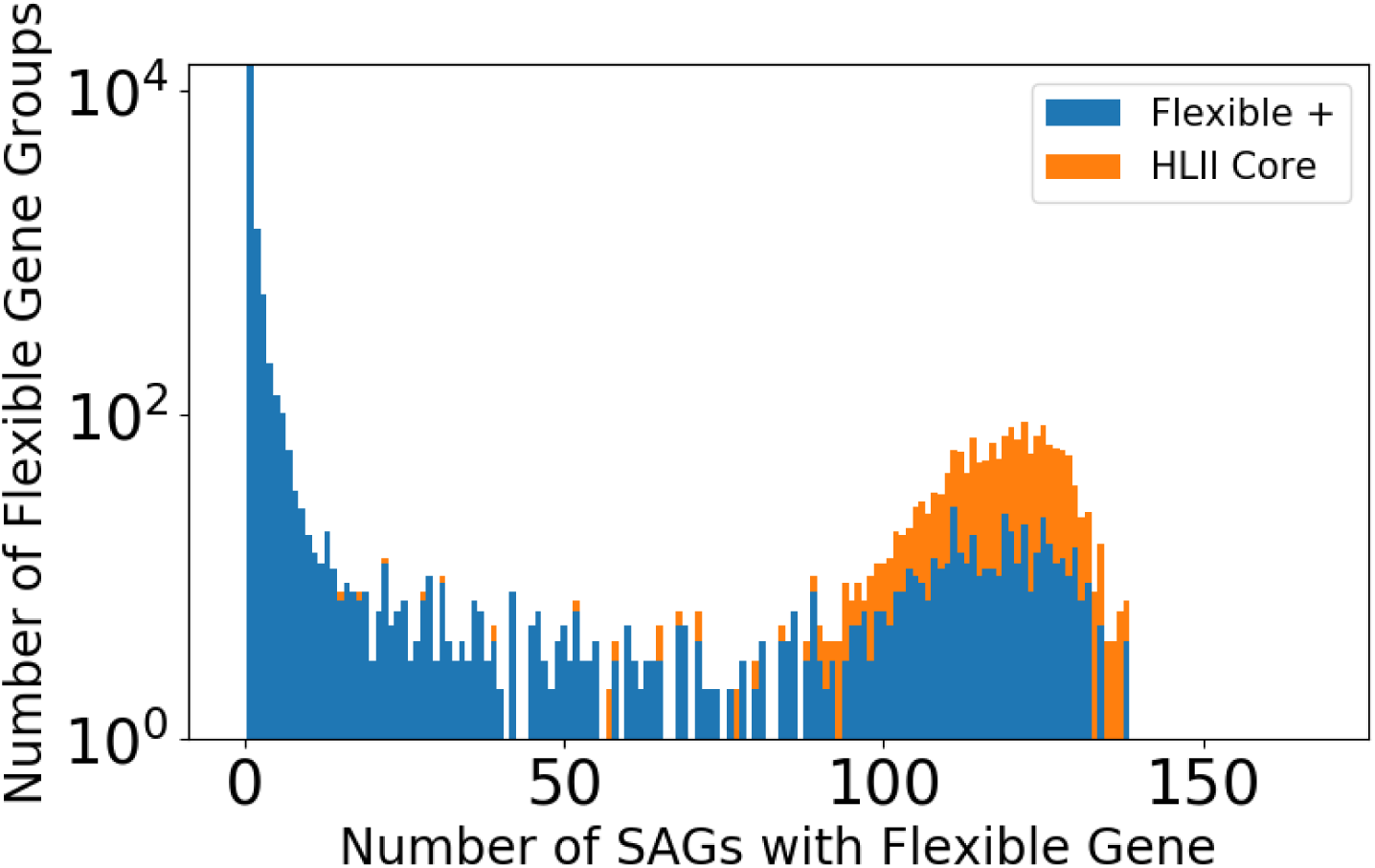
Distribution of sizes of core and flexible genes within the quasi-random dataset. Histograms show number of cells per gene for flexible (blue) and core HLII (orange) genes, as defined as explained in Sections B.3. and B.2.. The peak at high coverage in the flexible gene distribution is due to genes not found in one of the five HLII reference genomes used to construct the HLII core genome. These genes were excluded from analyses of flexible genes, as explained in Section B.1.).

##### B.3. Construction of flexible orthologous gene groups

To create clusters of orthologous gene groups for flexible genes, we cannot scaffold to a reference genome. We separated the ORFs of each SAG into core (those in an orthologous gene group of a HLII core gene) and non-core. We compiled all non-core ORFs across all SAGs and BLASTed the nucleotide sequences all against all. We then constructed a graph with edges connecting ORF pairs whose BLAST hit exceeded 60% coverage and 80% identity in both directions and which were not from the same SAG, and verified at the end that our classification was robust to variations of these parameters. We then found the connected components of this graph. We checked that each connected component contained at most one ORF from each SAG and found this this true in most cases. When a connected component contained more than one ORF from a SAG, we discarded ORFs until there was only one per SAG. This does not change the results for evaluating the presence or absence of flexible genes in various SAGs, which is the principle use of the flexible gene groups in this analysis. Duplicate ORFs per SAG in the same flexible orthogroup may result from genes being broken into two ORFs, from similar function genes having similar sequence and being grouped together, or from paralog flexible genes, for example.

We then equated the connected components to orthologous flexible gene groups. Note that the choice in query coverage to define a match has the largest effect on singleton and doubleton flexible ORFs, which are not used in our analysis of ocean-specific genes. Figure S8 shows the distribution of sizes of flexible gene orthogroups (the number of SAGs with a gene in the orthogroup that is covered) compared to those of HLII core.

Note that a fraction of flexible gene groups we identified using the method described above have similar sizes to the HLII core gene groups. To distinguish between low-coverage core genes and genuine flexible genes, we restrict most of our analysis of flexible genes to sizes of 4-34 SAGs in the quasi-random dataset, which we call *intermediate-frequency flexible genes*. These represent gene groups which are with high confidence part of the flexible genome, but which are still found in a substantial fraction of the population. We find a total of 421 intermediate-frequency flexible gene groups which we use for subsequent analysis in Sections 7. and 8..

##### B.4. Construction of core and flexible genes for deep clustered dataset

We also separately defined a core and flexible genome for the 51 C1 cells that form the deep clustered dataset in order to estimate flexible gene turnover among closely-related cells. We used BLAST (*7*) to align each each gene from the deep clustered SAGs against the C1 composite reference constructed by Kashtan et al. Mutual best hits with *>* 40% identity and *>* 40% of the query length covered were identified for each SAG. The distribution of the fraction of C1 cells with hits at each ORF in the composite reference genome was compared to a binomial distribution with *n* = 51 (the number of C1 cells) and *p* equal to the mean C1 SAG coverage. A scenario in which each ORF were present in every SAG would be represented by this binomial distribution. The ORFs that were consistent with the binomial were labeled as core genes and ORFs with lower coverage than expected by the binomial distribution we labeled as flexible genes. Additionally, we found a number ORFs that were present in more cells than would be expected due to coverage which we excluded from the rest of the analysis.

For each C1 SAG, we collect the ORFs identified by IMG that do not fall into the core or excess orthologous groups identified. These are taken to be the flexible genes for each SAG. Note that many of these flexible genes fall in one of a few *islands*. We BLAST the flexible genes of all C1 SAG pairs to determine shared flexible gene content. For each C1 SAG pair, we compare the number of shared flexible genes to the number which would be expected if the pair shared all of their flexible genes, given incomplete coverage. The number of flexible genes shared per pair as a function of the clonal backbone divergence is shown in Figure S10. The results show that hypervariable flexible genes are completely different by the time a cell pair is diverged of order 10^−2½^, a factor of ten faster than core genes turnover.

**Figure S9:**
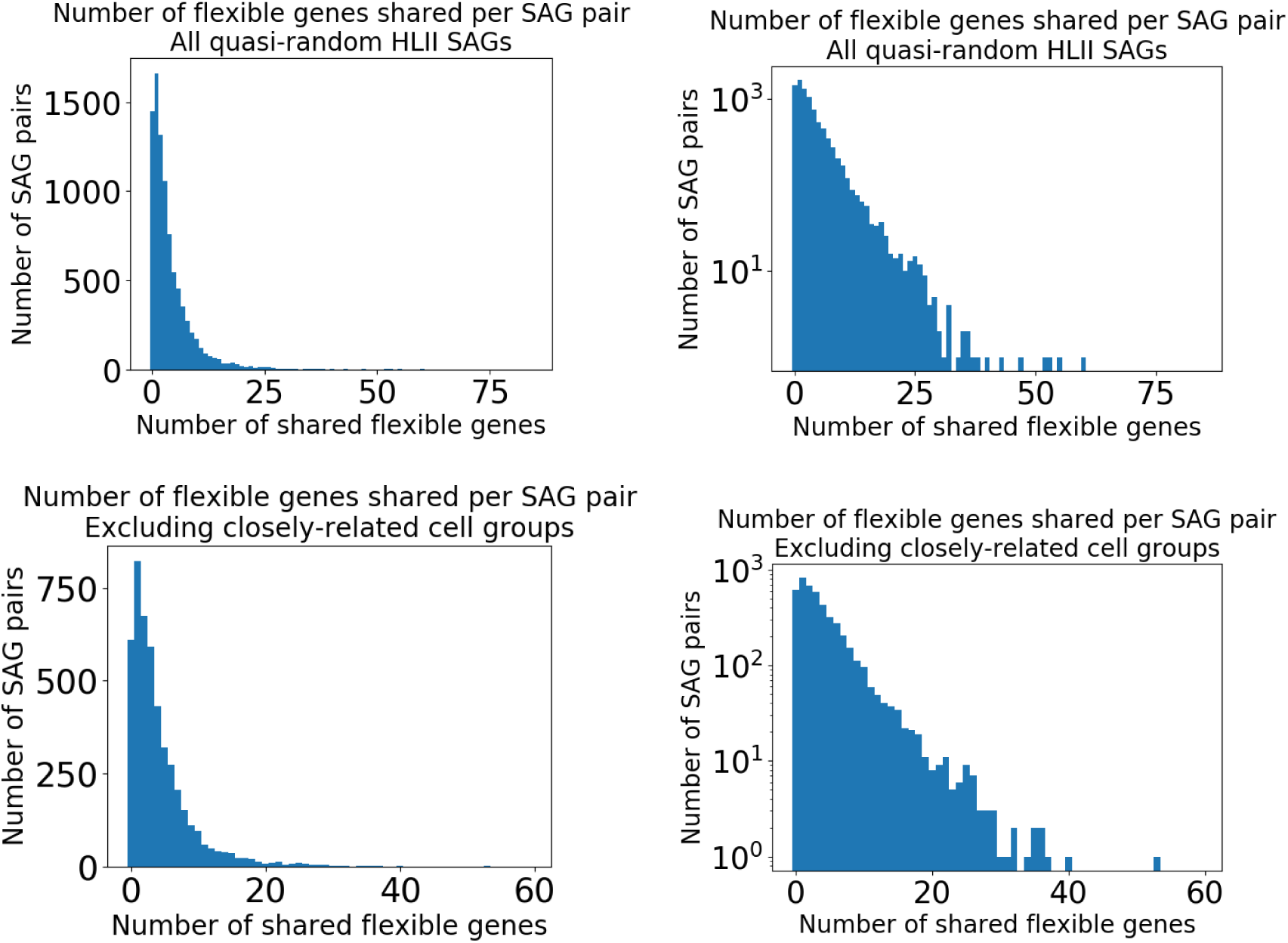
The number of flexible genes shared between SAG pairs has roughly exponential distribution with mean 4. Histograms for quasi-random dataset are shown in **top** panels and those for quasi-equidistant subset of HLII (Section 2.) are shown in **bottom** panels. Within each row, the same data is shown with the y-axis on a linear (**left**) and a logarithmic (**right**) scale. Most SAG pairs share 10 or fewer flexible genes, with the plurality of SAG pairs sharing just a single flexible gene.

**Figure S10:**
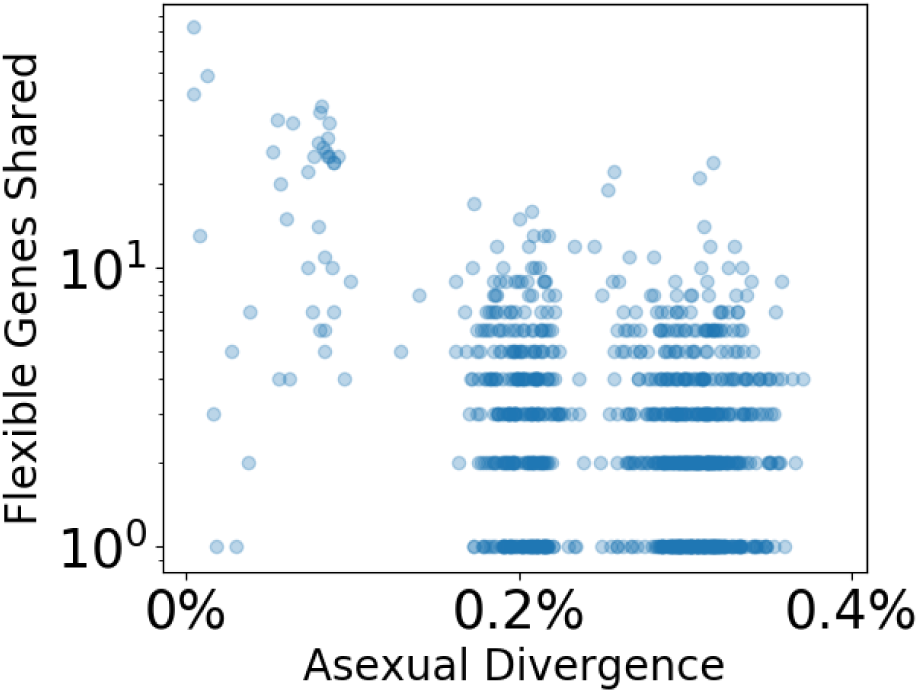
The number of shared flexible genes decreases with the asexual backbone divergence within C1. Each point represents a pair of C1 SAGs. The asexual divergence was estimated using the Poisson model described in Section A.1. and is shown on the x-axis. The number of shared flexible genes include all genes not part of the C1 core genome, as defined in Section B.4., and is shown on the y-axis.

### 2. Population Structure

In this section we characterize the population structure within the *Prochlorococcus* High-Light clades (HLI and HLII) using whole genome divergences. Based on this analysis, we define the *quasi-equidistant* subset of the quasi-random dataset, which is used in the main text and subsequent analysis. We also identify *tight clusters* of closely related SAGs within the quasi-random dataset which are used in later analysis in Sections 5. and 9.. Finally, we compare these results to ones based on ITS divergences.

#### A. HLI-HLII comparison

We use the median divergence across genes to characterize the population structure in HLI and HLII. Note that we only included SAGs from the quasi-random dataset for this analysis to get a relatively unbiased sample from both clades. We observe qualitative differences in population structure between HLI and HLII. Within HLI, we find 3 large clusters, within which divergences are significantly smaller (on a logarithmic scale) compared to typical HLI SAGs (Figure S11 left). Within these large clusters, several smaller cluster can be seen on increasingly smaller divergence scales. This pattern suggests some of these clusters may be due to ecological differences. By contrast, we find very few clusters within HLII and the ones we do have only a handful of SAGs (Figure S11 right). These results suggest that ecological differences may play less of a role in generating and maintaining diversity within HLII compared to HLI.

**Figure S11:**
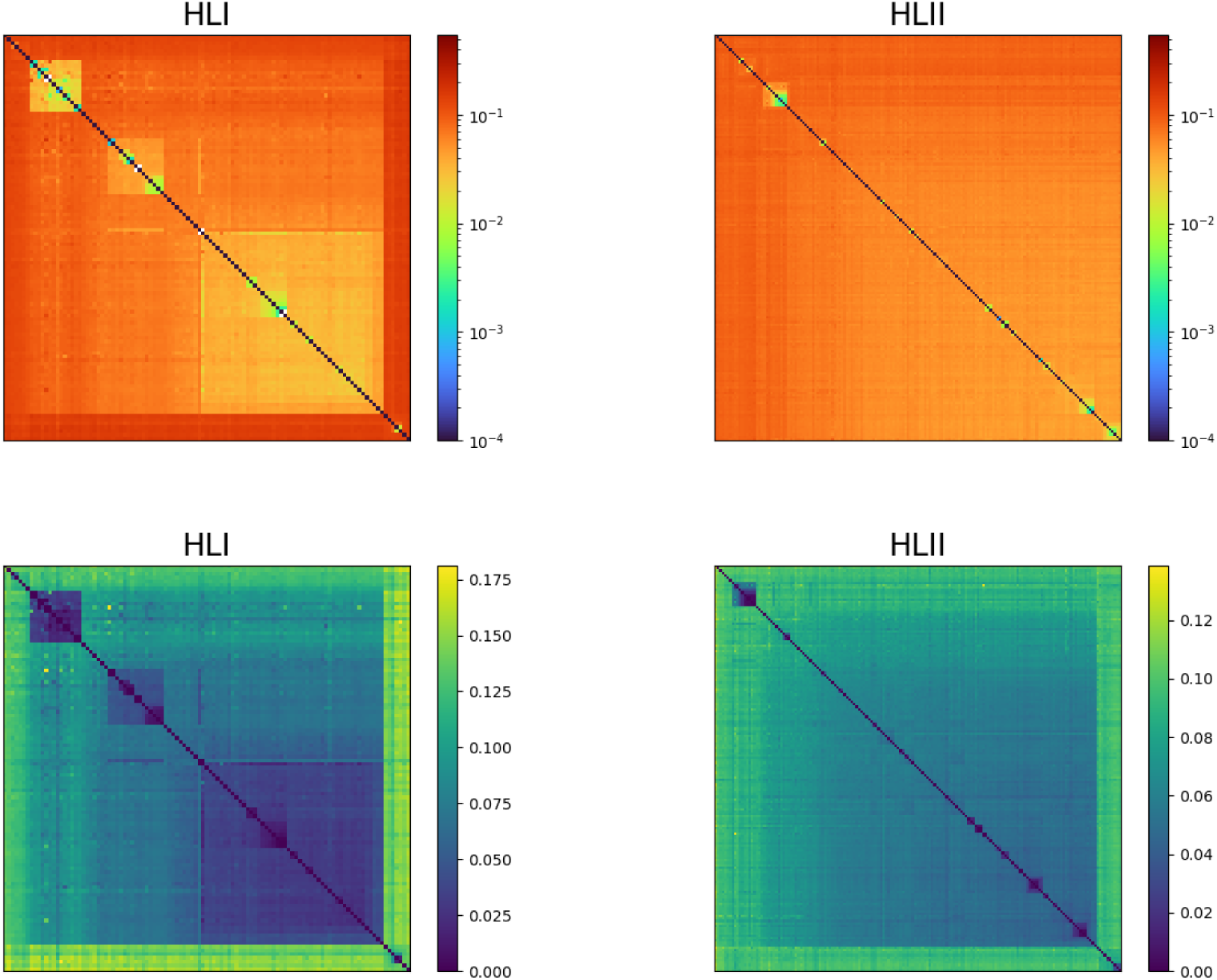
A quasi-random sample of HLI cells shows more structure at the whole genome level than a quasi-random sample of HLII cells. Divergence matrices are calculated using the median nucleotide divergence across core genes for each cell pair. **(Top)** Divergence matrices for HLI (**left**) and HLII (**right**) SAGs from the quasi-random dataset, as defined in Section 1.. The color scale is logarithmic and is the same as used in Figure 1 in the main text. **(Bottom)** Same as top but using a linear scale for the color.

#### B. Population structure within HLII

We searched for evidence of population structure within HLII by calculating the pairwise nucleotide divergence between all pairs of SAGs, averaged over shared genes. The distribution of divergences across pairs of SAGs from the quasi-random dataset has several peaks between 5% and 10% divergence (Figure S12). The most prominent peak spans 5-7% divergence and corresponds to a large cluster (blueish region in Figure S11, lower right panel) that can be clearly separated from other HLII SAGs using complete-linkage clustering. This cluster can be seen more clearly within the subsample taken from the BATS station (Figure S13). Within the larger cluster, several smaller and less diverged subclusters can be seen, including the C1 SAGs analyzed in (*2*). Note that the time scale corresponding to 5-7% divergence within this cluster is millions of years and is much higher than ocean mixing times (*9*). We would thus expect to find members of this cluster spread throughout the globe. Consistent with this expectation, we find each sampling site contains at least 5 SAGs belonging to this cluster, with roughly equal fractions within Atlantic and Pacific samples.

**Figure S12:**
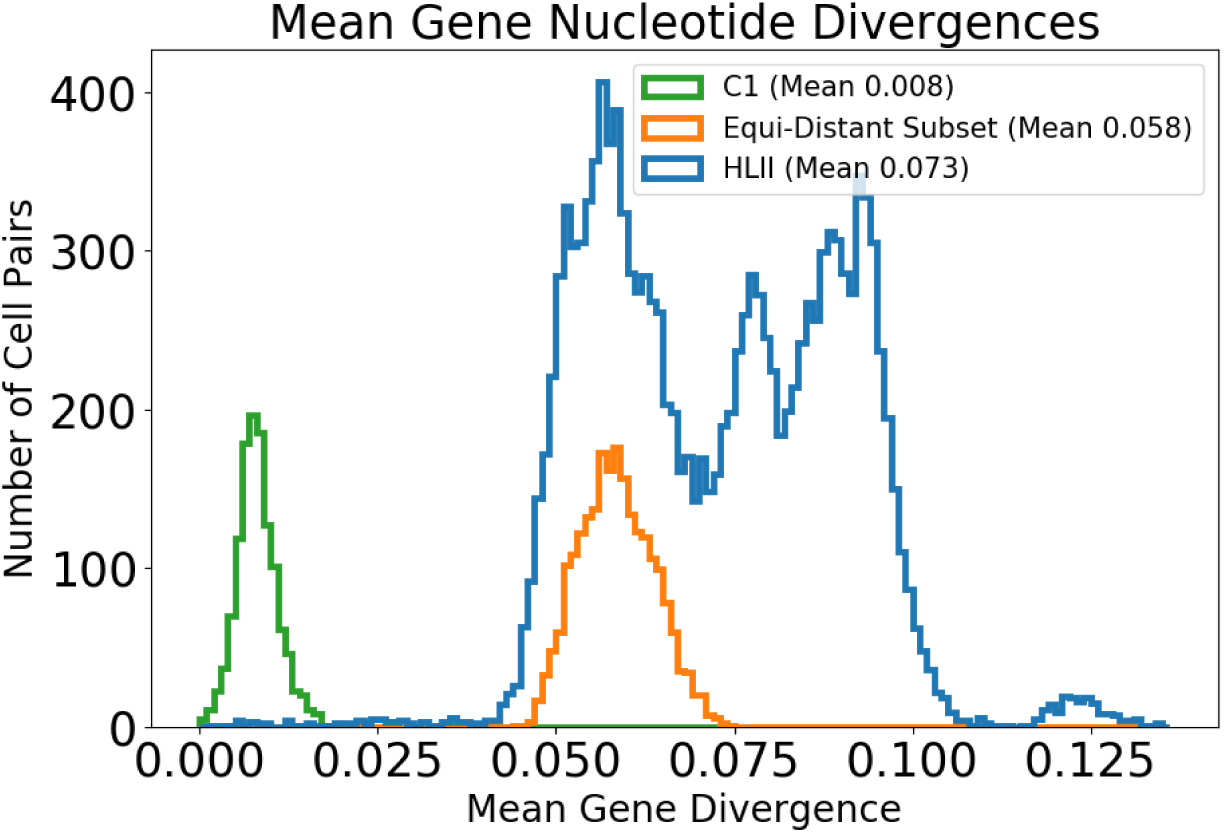
The distributions of whole genome divergences for three different cell groups: the quasi-random HLII dataset (blue), C1 from the deep clustered dataset (green), and the quasi-equidistant subset of cells from the quasi-random dataset that were chosen to minimize structure at the whole genome level (orange).

**Figure S13:**
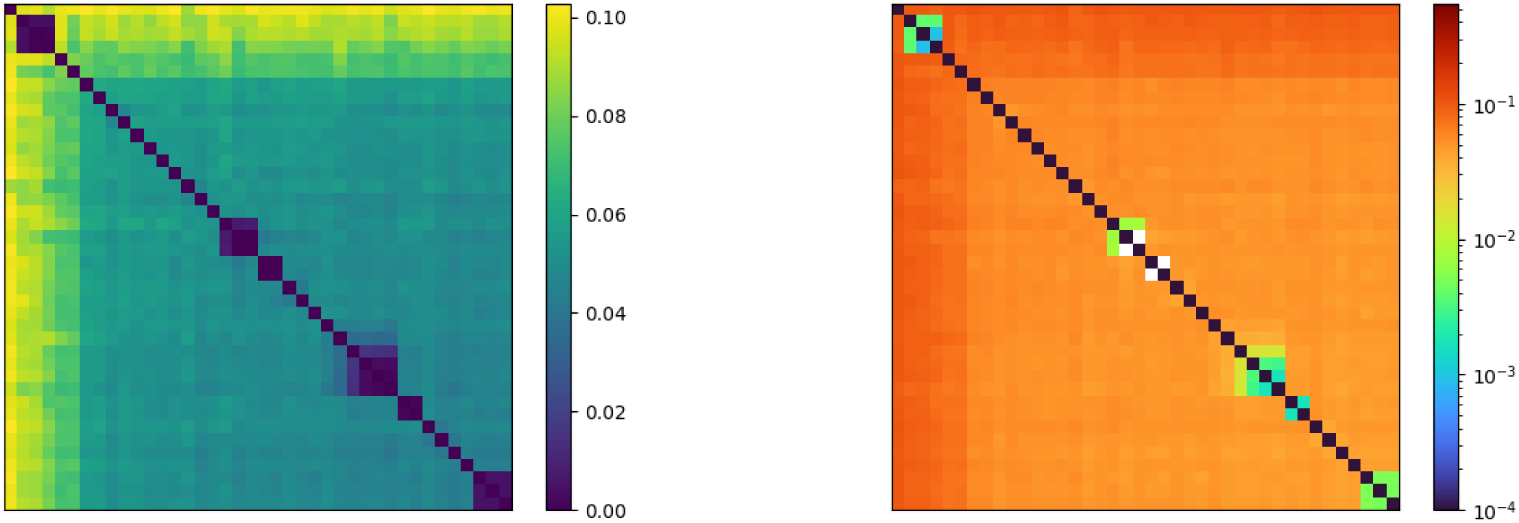
Whole genome divergence matrices for SAGs from the BATS sampling site in the quasi-random dataset showing several tight clusters, the largest of which comprises C1 cells, as well as a larger, more diverged cluster that comprises most of the cells. The later cluster was used to construct the quasi-equidistant subset of HLII (see text). Divergence matrices are shown on two color scales: a linear color axis over the full range of divergences in the matrix **(left)**, and the logarithmic color scale used in the whole genome distance matrices of Figure 1 in the main text **(right)**.

To better understand the evolutionary processes that generated and maintained diversity within the large cluster, we removed the tight subclusters of closely-related cells as follows. Using the scale of 4% divergence, below which there is a sharp decline in the number of cell pairs (Figure S12), we find cliques of cells that are diverged 4% or less from within the cluster. We remove all but one representative cell per *<* 4% group. The remaining group contains 68 SAGs, which we call the *quasi-equidistant subset* of HLII. The quasi-equidistant subset has a tightly peaked distribution of whole genome divergences (Figure S12) and is used for testing hypotheses regarding persistence of residual SNP or gene linkage at long length scales along the genome (Section 6.).

We also systematically identify *tight clusters* of closely-related SAGs within the entire quasi-random dataset using the same 4% divergence cutoff. Larger groups were identified as the connected components of a graph with edges connecting pairs with a mean core genome divergence less than the cutoff. We found 21 tight clusters of closely related cells within the quasi-random dataset, containing a total of 61 SAGs. The tight clusters were divided into 14 pairs, 4 groups of three cells, and 3 groups of seven cells, all mutually exclusive. We refer to these as *quasi-random tight clusters*. We label the 3 larger clusters of seven cells as QR-C6-8, to distinguish them from the CN2-C1-5 found in the deep clustered dataset (Section 1.). These clusters are used in the analysis from Sections 5. and 9..

To understand the origin of these tight clusters, we analyzed the divergence scales within and between them. The diameters of the tight clusters are overall comparable to C1 from the deep clustered dataset, but their number and size is dependent on the cutoff used to define them (Figure S14, top). Thus, there is no single characteristic scale for the tight clusters. Instead, we find a range of divergence scales, suggesting a scenario in which blooms occur continuously in time. In addition, pairwise distance matrices of these closely-related groups show that larger groups contain subclusters of even more closely-related cells, showing smaller blooms have occured within the larger ones (Figure S14, bottom). Above 4% divergence, typical HLII cell-pairs begin and the clustering structure is lost.

**Figure S14:**
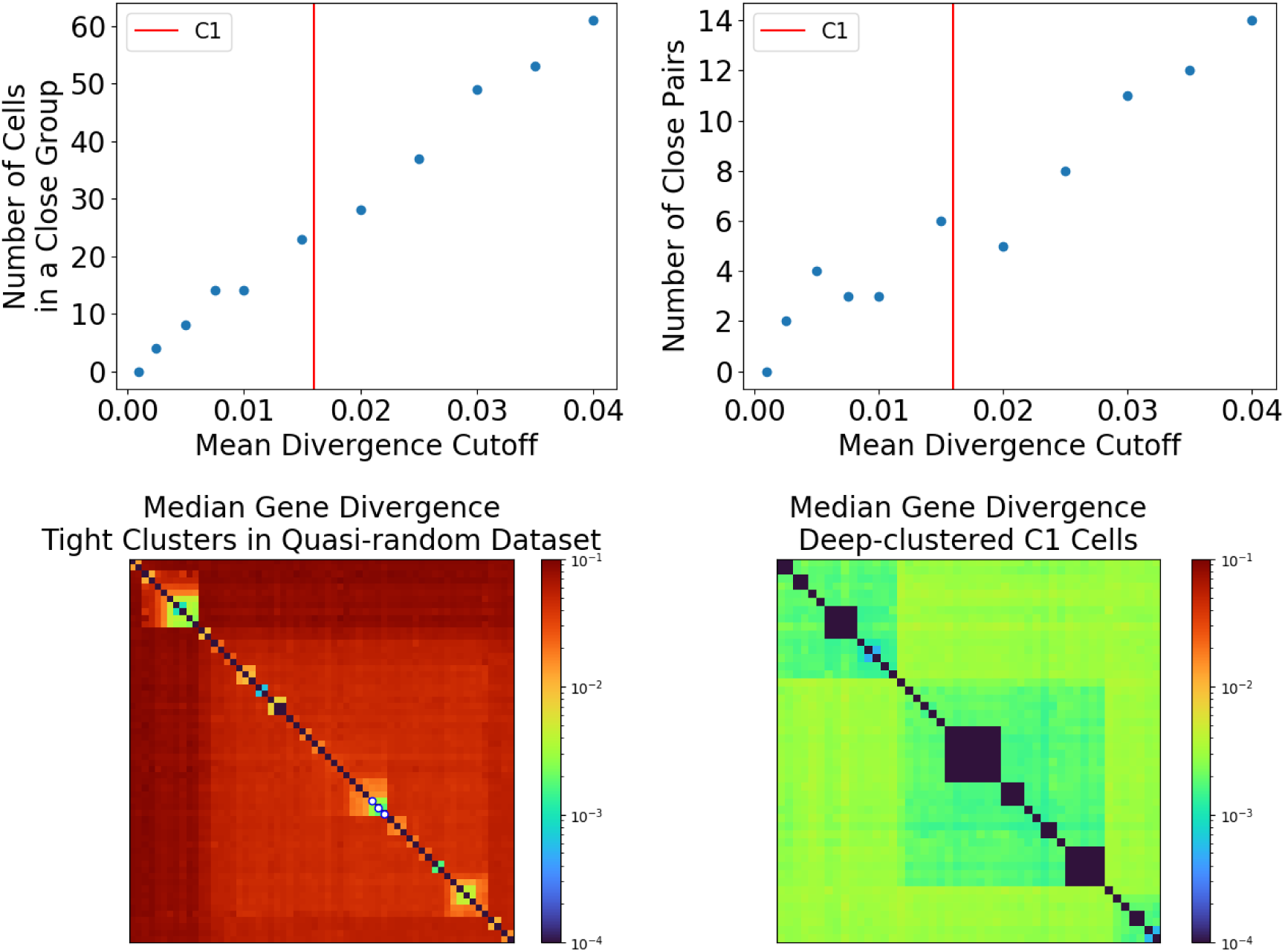
Groups of closely-related cells in the quasirandom dataset have a wide range of divergence scales. **(Top left)** The number of cells in the quasi-random dataset (167 SAGs total) that belong to a close group of at least 2 cells as a function of cutoff in the mean core genome divergence used to define groups (see text). **(Top right)** Same as top left but showing the number of close pairs (groups of exactly two cells) on the y-axis. **(Bottom left)** Distance matrix between quasi-random SAGs that are part of a close group with diameter of ≤ 4%. In total, 3 groups of seven cells each, four trios, and 14 pairs are shown. Distances were calculated as the median over nucleotide divergences of shared core genes for each pair. The three SAGs from the quasi-random sample that cluster with C1 at the whole genome level are shown with white points, outlined in blue, along the diagonal. **(Bottom right)** C1 SAGs from the deeply clustered dataset are shown for comparison using the same logarithmic color scale as in bottom left. Note that the color scale is a slightly narrower than the one used in Figure 1 in the main text.

#### C. Comparison between ITS and whole-genome divergences

How much population structure can be inferred from marker gene diversity? To answer this question, we compared the whole-genome divergences calculated previously to those calculated from the ITS only. Consistent with previous studies (*10*), we found that ITS sequences were reliable in distinguishing between genomes belonging to different clades. Surprisingly, within HLII we found that this was no longer true. As an example, we considered the CN2 group from the BATS dataset reported in reference (*2*). The ITS sequences from this group form a distinct cluster with a typical divergence of 2% (*2*). However, at the whole-genome level we found that CN2 spans most of the diversity of HLII and does not form a monophyletic cluster (Figure S15). This conclusion was robust for whole-genome divergences calculated using both synonymous and amino acid differences, and across different cluster linkage methods. These results are consistent with a scenario in which widespread recombination shapes the diversity within HLII.

**Figure S15:**
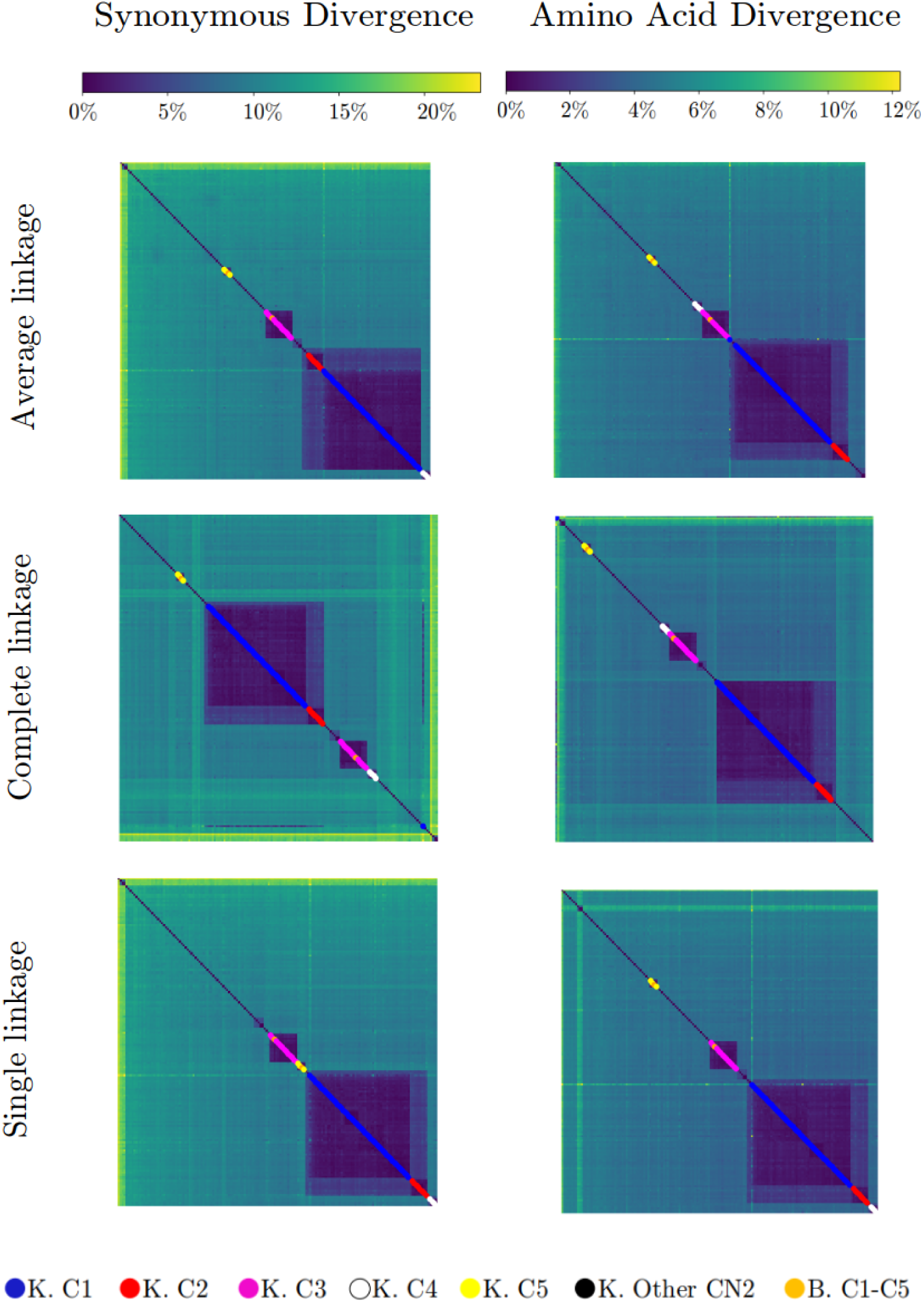
Mean core genome distance matrices for all cells from the deep clustered and quasi-random datasets. Divergences were calculated using both synonymous sites (**left**) and amino acids (**right**) separately. Distance matrices calculated using both measures were clustered using three different cluster-linkage algorithms: average linkage (**top**), complete linkage (**middle**), and single linkage (**bottom**). Dots along diagonal show cells from the deep clustered dataset colored according to the ITS ribotype identified in (*2*). Note that in all cases the ribotypes from the deep clustered dataset span most of the diversity of HLII and do not form a well-defined cluster, contrary to the ITS-based clustering from (*2*).

### 3. Mutation Rate, Neutrality of Synonymous Sites, and Calculating Divergences

In this section we estimate the mutation rates in HLII and discuss the impact of selection on synonymous sites on long evolutionary time scales. We conclude that synonymous sites are close to neutral over the time scales of HLII evolution.

#### A. Mutation rate measurement by Chen et al

Chen et al (*11*) measure the mutation rate of HLII strain AS9601 in the laboratory by serial dilution, reporting 3.50 ± 0.76 · 10^−10^ mutations per base pair per generation. Nonsynonymous mutations were included in this calculation. In the procedure presented by Chen et al, cells are allowed 3-4 weeks of growth between transfers. Using 0.947 cell divisions per day, as estimated by the authors, suggests about 10^6½^ cells grown per line between transfers starting from a single cell. Selection coefficients that are a few percent per generation are therefore important and selection cannot be ignored. Indeed, the *pN/pS* value obtained in Chen et al.’s experiment is significantly higher than *pN/pS* ≈ 0.3 which we find in typical core genes in the quasi-equidistant dataset, suggesting positive selection may have had a significant effect. Nevertheless, the accumulation of synonymous mutations in these experiments still provide a useful baseline for inferring time scales in natural populations.

We examined the 11 synonymous mutations reported by Chen et al. to gauge if the synonymous mutation rate differed from the reported rate. Using about 365,000 synonymous sites on the AS9601 genome, 141 cell lines in the experiment, 1258 generations per line, and 11 synonymous mutations, we gauge about 1.7 · 10^−10^ mutations per bp per generation. Using the 4:1 mutation rate ratio we inferred from the data (Section B.), this gives

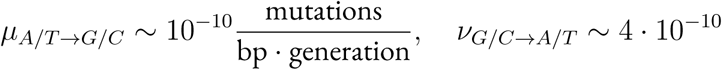

This agrees fairly well with the rate calculated by Chen et al. The uncertainty is approximately ±30% from counting statistics.

From here, we can estimate an overall mutation rate for twofold degenerate sites as

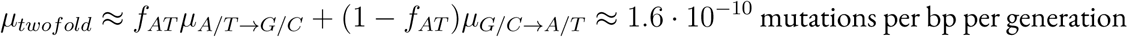

with *f_AT_* ≈ 0.8 the fraction of A/T alleles at twofold degenerate sites. To convert to an effective mutation rate for total nucleotide divergence in HLII, we multiply by the ratio of the total nucleotide heterozygosity to the twofold degenerate site heterozygosity, which is approximately ¾. The final result is

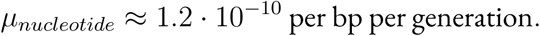

With roughly 10^2½^ generations per year, this gives *µ_total_* ≈ 10^−7½^ per bp per year. Taking into account the factor of 2 in nucleotide divergence since a common ancestor, a time *T* ago, *d* = 2*µT* gives an accumulation rate of pairwise divergences of 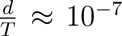 per bp per year with perhaps a factor of 10 uncertainty.

Note that among synonymous mutations reported by Chen et al., there is a roughly equal number of mutations that increase AT content as mutations that decrease AT content. Since there are about four times as many AT synonymous sites on the AS9601 genome as GC synonymous sites, there results are consistent with 4 : 1 mutational bias towards AT we inferred from the data in Section B..

#### B. Mutation bias towards A/T

##### B.1. Measuring 4-fold difference in mutation rates based on GC content

Within the HLII dataset there have been many generations since most recent common ancestor and synonymous sites have been subject to multiple mutations in the history of the sample. We examine the distribution across HLII cells of the fraction of A/G twofold degenerate core sites that have allele A per cell (excluding 6-fold degenerate codons) and find it tightly peaked (mean 0.801 standard deviation *<* 0.01 across 167 quasi-randomly-sampled HLII SAGs), as with the fraction of T/C twofold degenerate sites per cell with allele T (mean 0.798, standard deviation 0.01). This indicates that the twofold degenerate sites are in mutational equilibrium at typical HLII divergences. Note that 79.5% of twofold degenerate sites are polymorphic in this sample so the tightly-peaked distributions are not the result of an ancestral genome that has not undergone much mutation. In this scenario, we expect

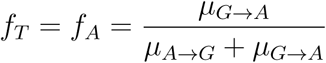

where the *µ*’s are observed mutation rates and *f_A_*, *f_T_* are the fraction of A at A/G twofold degenerate sites, and the fraction of T at T/C twofold degenerate sites, respectively. This gives

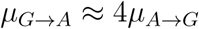

A mutation bias towards A/T has been found in other species, with ratios 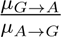 ranging from 1.4 to 4.4 (*12*). One notable exception are thermophilic cyanobacteria from the Yellowstone Caldera, which have a strong mutational bias towards G/C, resulting in 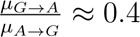 (*13*).

##### B.2. Validation of the neutrality of synonymous sites on HLII-relevant timescales

The neutrality of synonymous sites was validated by comparing the SFS for different two and four-fold degenerate amino acids. We found that the SFS are highly similar across all amino acids in each of the three categories: A/G twofold degenerate, T/C twofold degenerate (with one exception), and fourfold degenerate. Of course when comparing between the three categories, fourfold degenerate sites have much higher heterozygosity, as expected. There are very few monomorphic sites (Figure S16), although a slight excess of G or C monomorphic sites over theoretical expectations is observed, suggesting a small fraction (about ∼ 0.3% of sites) are under selection strong enough to affect the statistics. There may also be an excess of monomorphic A or T sites, but up to a few percent substantially affected by selection would be hidden among the synonymous monomorphic sites that are expected with the limited sample size if all sites were neutral.

**Figure S16:**
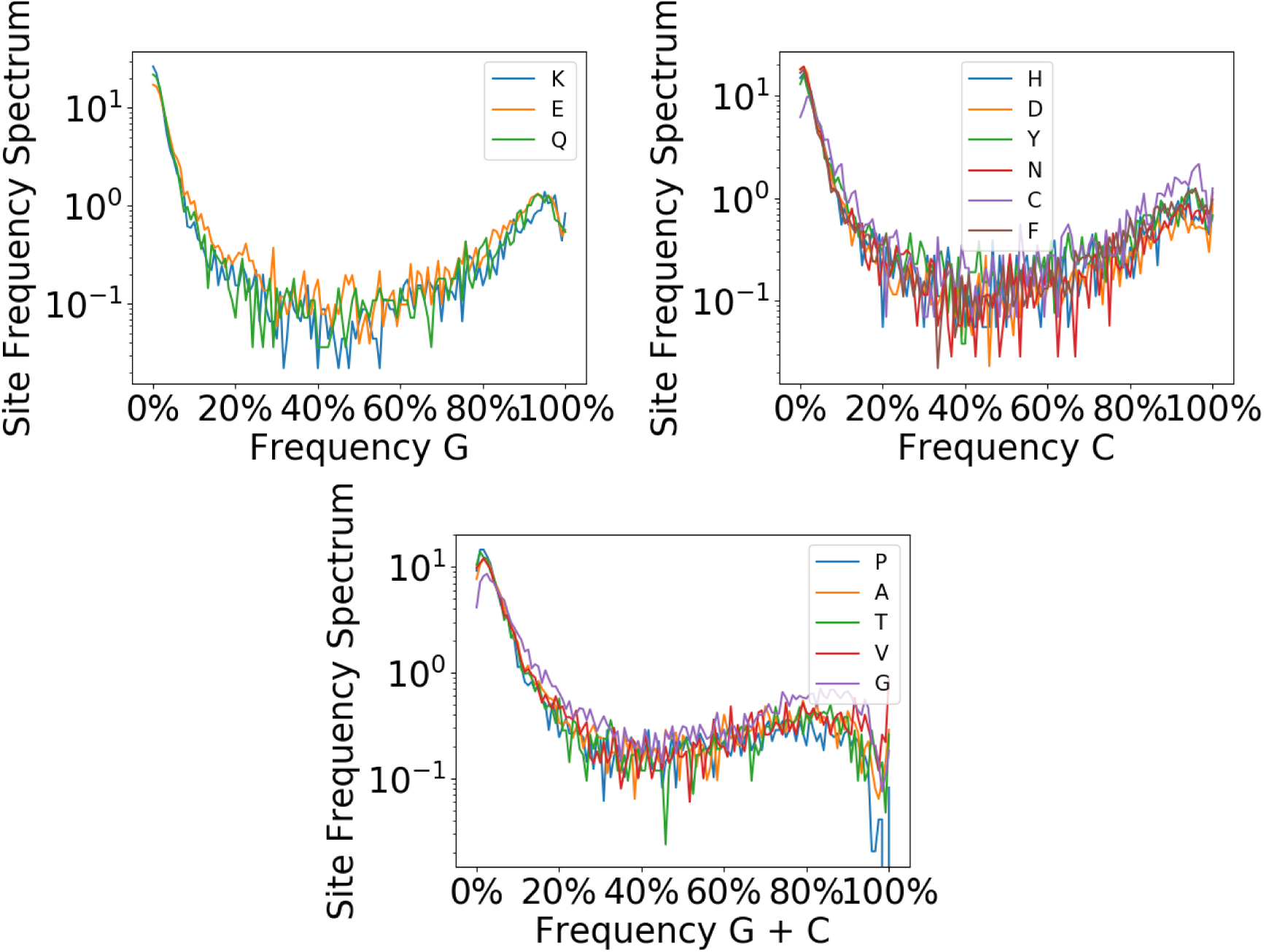
The site frequency spectrum of twofold and fourfold degenerate sites is highly similar across different amino acids (sixfold-degenerate amino acids are excluded). The three panels show the site frequency spectra of A/G twofold degenerate (**top left**), T/C twofold degenerate (**top right**), and fourfold degenerate sites (**bottom**) labeled by the amino acid coded by their respective codons. Generally, the curves are highly similar, indicating neutrality of synonymous sites on HLII-relevant timescales. Differences in shapes of the SFS, and/or differences in the proportion of monomorphic sites, would alternatively indicate selection on synonymous sites (for example, due to which tRNAs are present on the genome). Note the non-monotonic shape of cysteine site frequency spectrum (labeled “C” in the top right panel) at low C frequencies, which is distinct from other amino acids with T/C twofold degenerate site third sites (Section B.2.).

Cysteine is is a noteworthy exception; this T/C twofold degenerate amino acid differs in its frequency of the T vs C allele with respect to other T/C twofold degenerate codons, both in the SAGs studied here and in published HLII reference genomes. While typical T/C twofold degenerate sites have about 20% frequency of C, Cysteine has 32% frequency C. The heterozygosity of cysteine is additionally about 33% higher than the average of the other 4 T/C twofold degenerate codons (excluding sixfold degenerate amino acids). A possible explanation is that, while cysteine is produced from codons TGT and TGC, the other two possibilities at the third site produce Tryptophan (TGG) and a stop codon (TGA). This asymmetry could lead to a bias in the frequencies of the T vs C allele of cysteine, since nonsynonymous third-site mutations that produce cysteine might be more likely to occur from a tryptophan than from a stop codon. More cysteine produced as TGC would make the two somewhat out of mutational equilibrium increasing both the average C vs T frequency, and the heterozygosity. Cysteine is the only two-fold degenerate amino acid with this property.

There is a slight anomaly in comparing the heterozygosities of two fold and four fold sites, together with the estimates in Section B.1. and Section D. of ratios of relevant transversions to transitions: *T* → *G* vs *T* → *C* and the others that affect the overall *G/C* → *A/T* rates, and the reverse rates, for four fold sites. While the expected relationship of two-fold to four-fold heterozygosity does depend on the distribution of coalescent times, with the observed distribution (or even more so with a simple exponential distribution), the two-fold heterozygosity is somewhat smaller than expected. However, the statistics of all two-fold generate sites will be affected, as for cysteine, by mutations to and from the other amino acid which shares the first and second letters. Given that nonsynonymous mutations seem to occur at rates not that much lower than synonymous ones, this could affect the heterozygosity while leaving the average frequency similar. Understanding this would require substantial modeling which we have not carried out.

#### C. On typical HLII timescales, A/T content is not selected on

We simulate a null model in which there is no selection on AT content of synonymous sites (Figure S17). In this model, the dynamics of mutations away from G/C synonymous sites are the same as the dynamics of the sum of four mutations away from A/T. In other words, the only difference between the dynamics of mutations away from G/C and mutations away from A/T is that the mutation rate of the former is four times as large as the latter. In the simulation, we randomly draw four sites with majority A/T (up to about 30% minority allele frequency, to better ensure that sites are mutations away from A/T rather than mutations away from G/C that have risen to very high frequency) from the downsampled synonymous site frequency spectrum. We sum the frequencies of the minority alleles (including zero for monomorphic sites) and plot the convolution in comparison to the distribution of the frequencies of majority G/C synonymous sites. The curves agree remarkably well, for A/G twofold degenerates sites, T/C twofold degenerate sites, and fourfold degenerate sites as a function of *f_C_*+ *f_G_*. This shows that the asymmetry in the polarized site frequency spectrum can be explained by mutational bias alone without any selection on A/T content of synonymous sites. Note also that any neutral hitchhiking model, such as that which matches the SFS in Figure 5 (a) in the main text, implicitly includes zero selection on AT content: the agreement with the hitchhiking model is additional support for validity of the neutral but with mutational bias, hypothesis (see also Section 4.).

**Figure S17:**
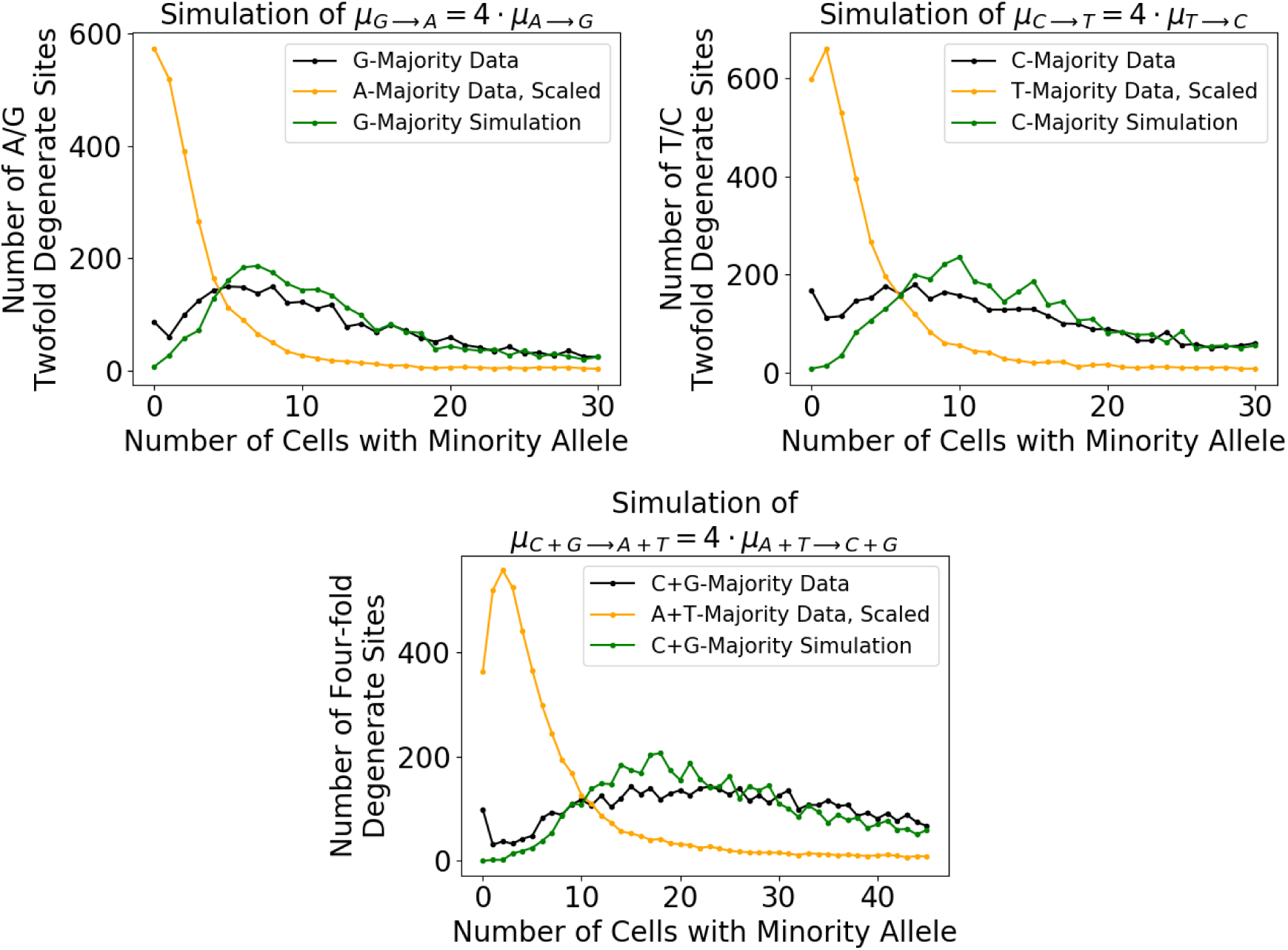
The frequency distribution of synonymous sites with majority alleles G or C is consistent with the distribution of the sum of four frequencies drawn from the frequency distribution of sites with majority alleles A or T. (**Top**) The frequency spectrum of A/G (top left) and T/C (top right) twofold degenerate sites within the quasi-random dataset is plotted, showing the minority allele frequency for sites binned by majority allele. (**Bottom**) Same as top but using fourfold degenerate sites. Orange curves show sites with majority allele A or T and blue curves show sites with majority allele G or C. Sites with high minority allele frequency are not included because the ancestral allele is highly uncertain in these cases. The green curve is a simulation of the frequency distribution of sites with majority allele G or C based on the sum of four random draws from the distribution of majority A or T sites, respectively. There is good agreement between the data and the simulation, suggesting that *θ_GC→AT_* ≡ 2*T_c_µ_GC→AT_ >* 1 and *θ_GC→AT_* ≈ 4 · *θ_AT →GC_* and that the asymmetry of the AT/GC-polarized synonymous site frequency spectrum can largely be explained without selection.

Shenhav et al. report codon usage bias at synonymous sites for the amino acids threonine and isoleucine within *Prochlorococcus* (*14*). While a full discussion is beyond the scope of our current manuscript, we note that one of their main results is that the third letter of threonine is biased towards T and C compared to A and G, respectively. The polarized SFS in Figure 5 (a) in the main text does not distinguish bias between A and T, or G and C. and our findings of neutrality of synonymous sites at the A/T vs G/C level does not directly contradict this finding. We leave more detailed comparisons as an interesting topic for future work.

#### D. Estimation of additional mutation rate ratios

We estimate the ratios of additional mutation rates using fourfold degenerate sites with low minority allele frequency (here, less than 10%). We use dimorphic sites among closely-related cells only (here, the deep cluster C1) to reduce to impact of homoplasy on the results. We select fourfold degenerate sites whose majority allele has at least 90% frequency and group these sites into mostly-A, mostly-T, mostly-C, and mostly-G sites. For mostly-A sites, we take the ratio of A-T dimorphic, A-C dimorphic sites to A-G dimorphic sites to estimate the ratio of 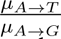 and 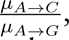 respectively. We perform an analogous analysis for mostly-T, C, and G sites, always normalizing by the rate of transition mutations. To attempt to use a common denominator, we quote mutation rate ratios for mutations away from A and T with respect to 4 times the transition rates *µ_A_*_→*G*_ and *µ_T_* _→*C*_), respectively, because of the relations 4*µ_T_* _→*C*_ ≈ *µ_C_*_→*T*_ and 4*µ_A_*_→*G*_ ≈ *µ_G_*_→*A*_.

The ratios computed for the C1 deep cluster, downsampled to 36 cells, is shown in Table S3. As a check, we used these ratios to calculate the steady state nucleotide frequencies and found *f_A_* ≈ 0.42, *f_T_* ≈ 0.41, *f_C_* ≈ 0.09, and *f_G_* ≈ 0.08. These values were within 1% of those obtained from the data (*f_A_* ≈ 0.4, *f_T_* ≈ 0.42, *f_C_* ≈ 0.1, and *f_G_* ≈ 0.08). Thus, the ratios in Table S3 provide a good fit to the data. But note that actual mutation rates between complementary bases – e.g., from *C* → *T* and *G* → *A* – should be the same as they depend only on the strand. We suspect the apparent differences might be due to context dependence of some mutations on neighboring sites (analogous to the case of cysteine discussed in Section B.2.) or possible selection on a small fraction of synonymous sites.

**Table S3:** Number of monomorphic sites and sites with singleton mutations at nonsynonymous sites in the quasi-random HLII dataset.

| Ratio | C1 estimate |
| --- | --- |
| $\mu_{T \rightarrow G} / 4\mu_{T \rightarrow C}$ | 0.09 |
| $\mu_{T \rightarrow A} / 4\mu_{T \rightarrow C}$ | 0.28 |
| $\mu_{A \rightarrow T} / 4\mu_{A \rightarrow G}$ | 0.38 |
| $\mu_{A \rightarrow C} / 4\mu_{A \rightarrow G}$ | 0.15 |
| $\mu_{C \rightarrow A} / \mu_{C \rightarrow T}$ | 0.53 |
| $\mu_{C \rightarrow G} / \mu_{C \rightarrow T}$ | 0.16 |
| $\mu_{G \rightarrow C} / \mu_{G \rightarrow A}$ | 0.15 |
| $\mu_{G \rightarrow T} / \mu_{G \rightarrow A}$ | 0.56 |

### 4. Synonymous Statistics: Site Frequency Spectra and Coalescent Time Distributions

In this section we introduce several models for the evolutionary dynamics and how these are probed by the statistics of synonymous sites, focusing on the site frequency spectrum and the distribution of coalescent times.

For simplicity, we consider both two-fold degenerate sites and four-fold degenerate sites but for the latter lump together A&T and G&C with the frequency, *f*, denoting the fraction, *f_GC_*of G and C lumped together. Note that for four-fold sites one has to use different effective mutation rates which group together *T* → *C* and *T* → *G* (as well as the complementary mutation rates which are indistinguishable) and likewise group together the opposite direction *C* → *T* and *C* → *A*: we will refer the combinations of these rates as *µ*_4_ and *ν*_4_ and the actual transition rates for the two-fold sites, as *µ*_2_ and *ν*_2_ respectively, usually dropping the subscripts. With perfect neutrality, which we assume (but see note in Section 3. B.2.), the average *GC* frequency *f̄* = *µ/*(*µ* + *ν*). For both types of sites, this yields *ν* ^∼^= 4*µ*.

#### A. Models of neutral dynamics and evolutionary history

##### A.1. Hitchhiking and effective drift

The dynamics of neutral markers such as synonymous SNPs are driven by demographic processes (including birth-death and migrational stochasticity or geographical variations in whole-population growth rates), as well as by differential selection on other genetic differences (*15*). Any process that increases or decreases SNP frequencies in an unbiased way is neutral. The dynamics of neutral markers may however be strongly influenced by non-neutral processes driven by other genetic changes on the same genome. A broad class of such processes is captured by a model in which one cell suddenly becomes fitter than it was because of a genetic change somewhere else on the genome: the synonymous allele of interest then hitchhikes on this selection until it either becomes unlinked from the selected element, or the expansion peters out. This results in the descendants of the neutral allele in the original cell jumping to a frequency *h* ≤ 1 with *h* the *hitchhiking factor*, and all the rest of the cells at that site – including those that share the neutral allele but not the selected element– decreasing in frequency by a factor of 1 − *h*. Thus if a *G* or *C* is the “lucky” neutral allele, *f_GC_* (which we denote as *f*) increases to *f* ^′^ = *h* + *f* (1 − *h*); if instead *A* or *T* lucky, *f* decreases to *f* ^′^ = *f* (1 − *h*). The chances that a *GC* is the one that hitches is *f* and the *AT*, 1 − *f*. Hitchhiking with different *h*’s happens at different rates: in general, we define a rate for hitches in the range (*h, h* + *dh*) to be *w*(*h*)*dh* (note that this integrates to a rate, not to unity). The limit that all *h*’s are extremely small behaves like demographic drift: if it is just a lucky birth or death, each *h* would be simply 1*/N*, the total population size. If the hitches are all much smaller than the sample size measured, the dynamics will appear like drift with an effective population size 1*/N_eff_*. (Unfortunately, this term is used far more broadly – roughly as the mean coalescent time in generations – and the source of much confusion: we will not use it here). Generally, the appropriately weighted overall hitchhiking rate, *H* ≡ ∫_0_^1^ *h*^2^*w*(*h*)*dh*, is the average pair-coalescent rate in phylogenies with the average coalescent time hence *T_c_* = 1*/H*.

The simplest process is when the hitches are fast enough that few mutations accumulate during them: the hitchhiking can then be treated simply as a jump process with mutations occurring gradually – and effectively deterministically in a huge population – in between hitchhiking events. (For considering effects of recombination not associated with the substantial hitchhiking events, these are again most simply considered as only occurring between hitches). In the simple approximation of exponential growth of the selected genomic element at rate *s*, the time for this driving element to sweep to a substantial fraction of the total population is *τ_sw_* ≈ log(*N*)*/s*. During this time the linkage between the synonymous site of interest and the driving element will decay at a rate *r* – most likely due to the element jumping around to other genomes – which reduces the hitchhiking caused by a full sweep to *h* ≈ exp(−*rτ_sw_*). If only a partial sweep occurs, this would be reduced further. In any case, the distribution of *r* will be reflected into a distribution of *h* which is very broad, naturally on a scale of log(*h*).

If recombination were so fast that only sites very close to the driving selected element are substantially hitchhiked, then the distribution of distances, *ℓ*, between the sites would give rise to a distribution of *r* ≈ *R_X_ℓ*, with *R_X_*the effective recombination break point rate per genomic distance. This, combined with the uniform distribution for the site to lie at a genomic distance between [*ℓ, ℓ*+*δℓ*], immediately gives *w*(*h*) ∝ 1*/h*. However for *ℓ* ∼ *L* (with *L* the typical length of the segment that is transferred around with the selected element), all of the further-away sites on the genome become unlinked at the same rate ∼ *R_X_L*. Thus for all relatively fast sweeps with *τ_sw_* = *k/R_X_L*, the whole genome will hitchhike by *h* = *e*^−*k*^. As perhaps hundreds as many sites are affected by such genome-scale events, they could play a major role.

For bacterial populations with extremely rapid decay of linkage, such as thermophilic *Synechococcus*(*16*, *17*), it appears that such genome-wide hitchhiking events are very rare. But for *Prochlococcus*, linkage decays more slowly with separation and we know that persistent genome-wide hitchhiking has occurred, as evidenced by in the predominantly asexual blooms. If these genome-wide events are predominantly quite small *h* – only the large ancient bloom would appear to have had large *h* – then they primarily contribute extra to the small *h* part of the distribution *w*(*h*). A crude model is

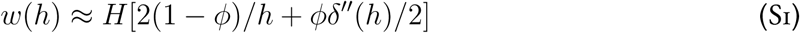

with the second derivative of a delta function capturing the excess small *h* parts. In writing this, we have scaled the average pairwise coalescent rate, *H* to be fixed so that *ϕ* is the fraction contributed to this from the genome-wide small *h* events, as well as possibly very partial genic sweeps. Potentially, genome-wide hitchhiking events could be large enough to give an excess of *h* near unity and dominate over genic-scale events for large *h* because of the much larger number of sites they affect. However, it is natural to assume that an asexual bloom to a very large fraction of the population – even if possible without ecological limitations – would rapidly be suppressed by phages and thus not give persistent changes in frequencies of synonymous variants (or be still evident as tight genomic clusters).

We can make a crude estimate of the rates of persistent genome scale hitchhiking from the distribution of the numbers and sizes of tight clusters as a function of their divergence (Figure S14). If small *h* ≤ O(1*/S*) – with *S* the sample size –dominate, then there would be a lot of coalescences – at initial rate roughly proportional to *HS*(*S* − 1)*/*2 – in the near past. Such are not seen. If somewhat larger *h* dominate, the rate would still be much larger than *H* while the actual observed rate is *< H/*2. This is still true after taking into account the likely absence of large *h* events in the recent past (even though on average over many population histories, they could have caused a large fraction of the recent coalescences). Thus it appears that the full genome coalescences are too infrequent to play much role in the overall hitchhiking statistics felt by SNPs. Therefore for simplicity – and to avoid a prolideration of parameters – we use the quasi-sexual genic-scale hitchhiking model with *ϕ* = 0 so that *w* = 2*H/h* which has only a single parameter.

In general, the most important parameter is the mean-square-weighted rate of hitchhiking, *H*, whose inverse is the average coalescent time. This – more specifically its ratio to a combination of the mutation rates – determines the heterozygosity. The simplest assumption for the population is that it is in a statistical steady state of hitchhiking with the ancestral population in mutational equilibrium. In this scenario, the probability that the ancestral state was *C* or *G* is *f̄* = *µ/*(*µ* + *ν*) and the heterozygosity is:

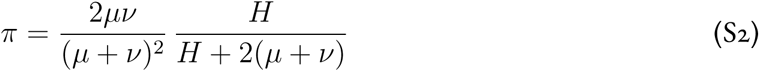

with the first factor what the heterozygosity would be if the alleles at all cells were independent: *π*_∞_ = 2*f̄*(1 − *f̄*), and the second the consequences of correlations due to the phylogeny. (Note that this expression is identical to that for drift with *H* replaced with 1*/N_eff_* : as noted earlier, effective drift is a special case of hitchhiking.) The mutational imbalance, as measure by *f̄*, and the hitchhiking rate, *H* in units of the mutation rates – inferred from *π* –, are the minimal set of parameters for any neutral model.

##### A.2. Uniform expansion

The other process we consider is a slow uniform expansion from a very small set of ancestral cells to a population of *N* ∼ 10^27^(although only log(*N*), which has little uncertainty, matters). Simplest is an expansion at a steady exponential rate, *γ*, extending to the present with no major fluctuation events – such as hitchhiking – except initially. Again, there is a single parameter.

#### B. Site frequency spectrum

The general hitchhiking process can be analyzed either from the dynamical integro-differential equation for the site frequency spectrum, *ρ*(*f*), or by computing the dynamics of moments of *f* : just as for drift, the moment equations close. For finding sampling probabilities out of a sample of size *S*, all moments up to order *S* are needed and these can be combined (exactly) into a matrix equation which also includes effects of mutations. If the population is in *evolutionary steady state*, then the SFS for a sample can be found directly as the stationary solution to the dynamics of the moments of *ρ*(*f*). The computations and data are for a sample of *S* = 120.

##### B.1. Effective drift SFS

In steady state, the SFS for effective drift (and for the general hitchhiking process with only small hitches with *h* less than 1*/S* in a sample of *S*) has the well-known form:

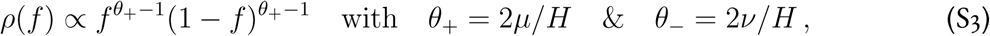

the asymmetry showing up in the sharpness of the peaks. For *θ*_±_ both small, most of the weight is very near 0 and 1 and concomitantly the heterozygosity is low. Fitting *π* and *f̄*yields the black curve in Figure 5 (a) in the main text, which is very far from the data shown. Note that even if *θ*_±_ had been *>* 1 (it is close to this for four-fold sites) so that there was a minimum rather than maximum at *f* = 1, there would only have been one peak: at *f* = 0 unless also *θ*_+_ were *>* 1.

##### B.2. Hitchhiking SFS of HLII and quasi-equidistant subset

For hitchhiking with *w* = 2*H/h* and the dimensionless ratio (*µ* + *ν*)*/H* chosen to give the correct *π*, the steady state SFS for a sample of 120 is shown in Figure 5 (a) in the main text (red curve). Note the remarkably good – and perhaps fortuitous? – agreement with the data for the lumped four-fold degenerate sites SFS of the full quasi-random HLII sample. This simple model certainly captures the main features: a peak substantially below *f* = 1, and the hints of a peak at low frequency. To get these with the given heterozygosity and mutation rate asymmetry requires a substantial amount of hitchhiking with *h* not small.

Adding a small amount of weak hitchhiking events (*ϕ >* 0) somewhat smears the low frequency peak but does not – at this resolution of frequency caused by the low sampling depth – noticeably change the shape, and for *ϕ* = 0.05 the fit at small *f* is somewhat better. However by *ϕ* = 0.1, the curves are substantially worse fit to the data in the mid-range of *f*.

Including an excess contribution of very large events with *h* close to unity affects the SFS minimally in the middle but makes the fit worse for *f* close to unity already for *ϕ_strong_* = 0.2. Of course, adding extra either small or large *h* events comes at the expense of adding one or more additional parameters.

To test the hypothesis that the dominant hitchhiking processes are genic rather than genome scale, we plot the SFS and the hitchhiking prediction for the quasi-equidistant subset in which there are no tight clusters – and thus no direct evidence of genome-scale hitchhiking – and almost no genome-scale linkage. The data and hitchhiking prediction with the simple *w*(*h*) = 2*H/h* with a somewhat smaller *H* reflecting the shorter mean coalescent time of this group (obtained again by fitting the heterozygosity which is about 15% smaller) are shown in Figure S18. The agreement is again remarkably good, supporting the conjecture.

**Figure S18:**
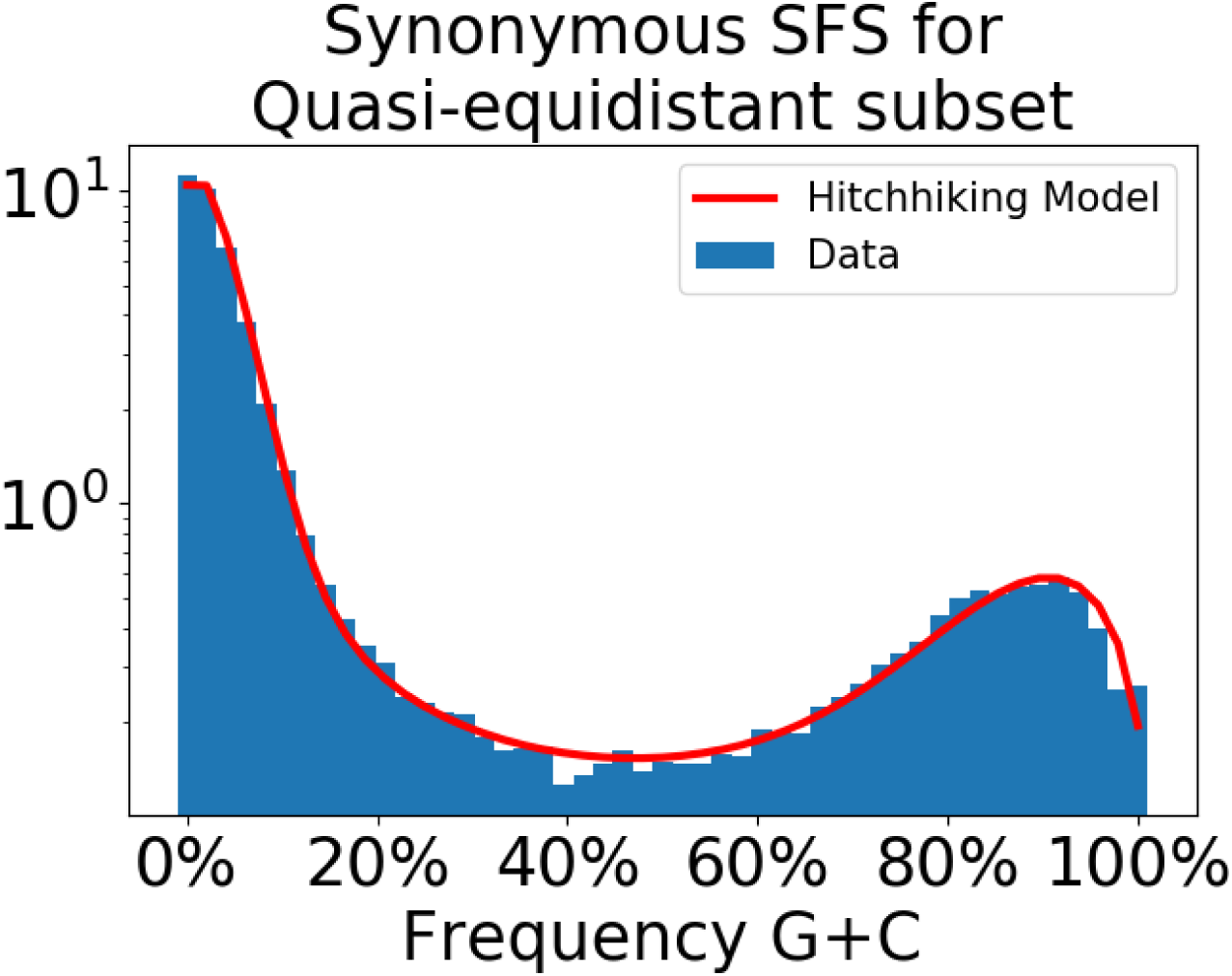
Synonymous site frequency spectrum (using fourfold degenerate sites at core genes) for the quasi-equidistant subset, subsampled to 48 cells, compared to a genetic hitchhiking model (with no free parameters (Section 4. A.1.)). The SFS agrees remarkably well with the hitchhiking model. This demonstrates that, while the quasi-equidistant subset of cells may be interpreted as an ancient bloom, it was not the result of a uniform expansion. Instead, extensive hitchhiking on gene and genome-wide sweeps and partial sweeps has occurred within this bloom. Consistently, tight clusters of cells exist within the ancient bloom at the whole genome level.

##### B.3. Expansion SFS

In a scenario in which the population results from an exponential expansion from a single ancestor, the population is not in an evolutionary steady state. Initially mutations occur rarely in a Luria-Delbrü ck process. If an allele starts from *A/T* and grows exponentially at rate *γ*, this gives at early times on average an SFS *ρ* ≈ *µ/*(*γf* ^2^). However, during this early stage most sites will not have been mutated at all, so most of the weight is in a delta function at *f* = 0. For initially *G/C* sites, the spectrum in this regime is *ρ* ≈ *ν/*[*γ*(1 − *f*)^2^]. If initially in mutational equilibrium, then averaging over sites the first part will be multiplied by (1 − *f̄*) and the second by *f̄* so that both have equal weight: except for the peaks at 0 and 1, the SFS will appear symmetric. (Note this symmetry for intermediate frequencies also occurs for small *ν/H* with effective drift.)

With the continuing exponential expansion, once the population becomes large enough that all mutations occur every generation – roughly *N* (*t*) ∼ *N_M_* = 1*/*(*µ* + *ν*) – the mutations become essentially deterministic. At each site, *f* (the frequency of *G/C*) evolves deterministically and the spectrum is squeezed towards *f̄* from both sides, with more rapid squeezing for larger *f* since *f̄* is small. The resulting form can be well-approximated analytically by delta functions at the squeezed frequencies from *f* = 0 and 1, and the squeezed Luria-Delbrü ck spectrum in between. The peaks are at *f*_−_ = *f̄*(1 − *E*) and *f*_+_ = (*E* + *f*_−_) with *E* = exp[−(*µ* + *ν*)*T_M_*] the decay of correlations since the time in the past, *T_M_* ≈ log(*Nɛ*) at which the population was large enough that the mutations became deterministic: 1*/N* (−*T_M_*) ≈ *ɛ* ≡ (*µ* + *ν*)*/γ*. The SFS at the end of the exponential growth is then non-zero between *f*_−_ and *f*_+_ with

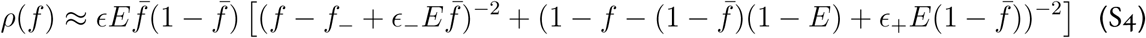

with *ɛ*_−_ = *ɛ*+*ɛ*^2^*f̄*[1−log(*f̄*)+log(1−*f̄*)] and *ɛ*_+_ = *ɛ*+*ɛ*^2^(1−*f̄*)[1+log(*f̄*)−log(1−*f̄*)] the corrections giving the correct heterozygosity. This is the one fit parameter) *π* = *π*_∞_(1 − exp[−2(*µ* + *ν*)*T_c_*]), the same as for the other processes except here there is no average over the distribution of coalescence times as they are almost all very near to when the expansion started: *T_c_* = log(*Nγτ_gen_*)*/γ* + O(1*/γ*) with *τ_gen_* the generation time.

The above expression is more than accurate enough for a sample of S=120. Choosing the one parameter, *γ* ≈ log(*N*)(*µ* + *ν*)*/p* with 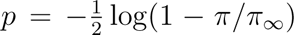 to fit the heterozygosity (as done for the other fits) yields the sampled result in Figure 5 (a) in the main text (orange curve). In the limit of a much larger sample, there would be essentially no weight outside of the two peaks: for the actual parameters, this is seen above the upper peak, but the lower one is smeared out by the shallow sampling. Although the asymmetric two peak structure is qualitatively like the data, more quantitatively it is not good. Note that if the model had been changed to include effect of expansion stopping before the present, the peaks would have been squeezed in even more, approximating the data even worse.

#### C. Downsampling for the site frequency spectrum (and linkage correlations)

Singletons and doubletons are informative features of the site frequency spectrum and require a discretized distribution to discern. Because each site is covered by a variable number of cells due to amplification, for the site frequency spectrum we discard all sites with less than the mean coverage and down-sample all sites, sampling without replacement, that are covered by more than the mean number of cells – 120 for the quasi-random sample, 48 for the quasi-equidistant set, and 37 for the deep sample of C1. Downsampling adds clarity to the nonzero peak at low C/G frequencies. But the features most affected by downsampling are the fractions of singletons and of monomorphic sites as the ratio of these provides information on the weighted average of low-frequency polymorphisms. However, this ratio is roughly consistent between the original and downsampled data (Table S4).

**Table S4:** Singletons and monomorphic sites in original and downsampled data for the quasi-random HLII dataset and the C1 SAGs from the deep clustered dataset

| Cell Group | Site Type in Codon | Downsampled? | Frac. Sites Monomorphic | Frac. Sites Singleton |
| --- | --- | --- | --- | --- |
| Quasi-Random HLII | Second site | No | 74.3% | 12.8% |
| Quasi-Random HLII | Second site | Yes | 77.1% | 8.4% |
| Quasi-Random HLII | Fourfold degenerate | No | 4.0% | 6.7% |
| Quasi-Random HLII | Fourfold degenerate | Yes | 2.6% | 3.5% |
| C1 | Second site | No | 97.6% | 2.9% |
| C1 | Second site | Yes | 97.6% | 1.8% |
| C1 | Fourfold degenerate | No | 86.5% | 8.7% |
| C1 | Fourfold degenerate | Yes | 87.6% | 8.1% |

For quantities involving pairs of SNPs (linkage measures such as ⟨*r*^2^⟩, SNP pairs incompatibilities, etc.), the coverage of SNP pairs is downsampled to 50% of cells in the total cell group, which is roughly the expected SNP pair coverage based on genome recovery statistics (we used 80 cells for the quasi-random HLII sample, 34 for the quasi-equidistant subset, and 25 cells for the deep sample from C1).

#### D. Could the synonymous site frequency spectrum be due to selection?

Site frequency spectra that do not look like drift are often interpreted in terms of a distribution of selection coefficients – not *apriori* unreasonable here even for synonymous sites given the very long divergence times. Although the concordance between SFS across amino acids strongly suggests otherwise (Sec. B.2., Figure S16), we here ask: What distributions of selection would be required in order to produce the synonymous SFS?

The stationary distribution for a site frequency spectrum with drift and uniform selection (”mutation-selection balance”) is

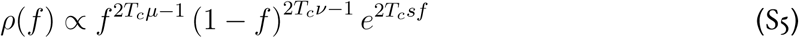

where *s* is the selection coefficient. *T_c_* is the mean coalescent time, *µ* the mutation rate per time from AT to GC; and *ν* from GC to AT. In the data, we measure *ν* ≈ 4*µ* but if selection is responsible for the shape of the SFS, then there could be different ratios of mutation rates. However, the observed anisotropy of the lab-measured mutation rates (Section 3. A.) is consistent with mutation rate dominating over selection (Section B. B.1.) to determine the anisotropy.

In the limit where *T_c_µ* ≫ 1 and *T_c_ν* ≫ 1, the dynamics for both selection and mutations are essentially deterministic and this results in a tightly peaked distribution centered around frequency *f* ^∗^, with *f* ^∗^ given by (*15*)

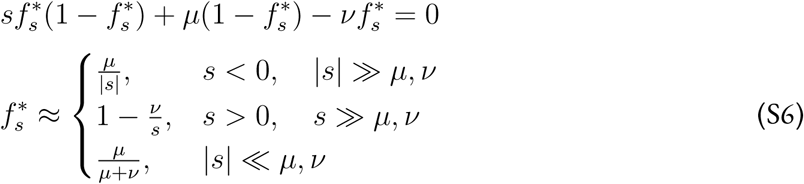

The whole SFS would be obtained by averaging over the *s*’s for each site. In this limit *T_c_µ,* & *T_c_ν* ≫ 1, the SFS would be a sum of peaks each with position determined by mutation-selection balance for individual sites along the genome, each site with its own positive or negative fitness *s* (such that an A or T allele at the site has relative fitness 0, and a G or C allele at the site conveys relative fitness *s*). The observed peaks of the synonymous SFS occur at *f* ≈ 1%, which would correspond to the regime where *s <* 0 and |*s*| ≫ *µ, ν*; and at *f* ≈ 95%, which would correspond to the regime where *s >* 0 and *s* ≫ *µ, ν*. Using the mutation rate measured using data from Chen et al (Section 3. A.), these peaks would correspond to |*s*| ≈ 10^−8^ per generation. Accounting for the widths of the peaks would give a range from |*s*| ≈ 10^−9^ to |*s*| ≈ 10^−7½^. The vast majority of sites in the fourfold degenerate SFS would have to fall within this range. This would require a remarkable coincidence of a very fine range of exceedingly low fitness values for nearly all fourfold degenerate sites along the core genome – and even more so to get the SFS for each amino acid to be so similar. This range of fitness values is too narrow to provide a convincing explanation for the synonymous SFS. In addition, these intermediate-frequency peaks would require that *µ*, *ν* and |*s*| were of comparable values, another remarkable coincidence. One could also try with intermediate *T_c_* so that the peaks from (Equation S5) would not be sharp. But that would only introduce another parameter – *T_c_* – which would have to by coincidence be similar to *both* the mutational and selection time scales as we know that for |*s*| ≪ *µ* the shape of the drift SFS is wrong.

Note that while any SFS can trivially be fit in the deterministic limit by a perfectly tuned distribution of *s*, we suspect that most such distributions will be incompatible with *any* neutral model. Thus the agreement with the simple hitchhiking model, and more generally with the approximation of the *GC* end of the spectrum by convolution of 4 of the *AT* end, already makes a very good case that neutral processes rather than selection dominate the synonymous site statistics.

#### E. Nonsynonymous site frequency spectrum

For nonsynonymous sites, of course the SFS reflects selection. Figure S19 shows the AT/GC-polarized site frequency spectrum for the second letter of each codon in the HLII core genome, for the quasi-random HLII dataset. The second letter is always nonsynonymous. The nonsynonymous SFS has peaks at *f_GC_* =0% and 100%, in contrast to the peaks at intermediate AT content in the fourfold degenerate SFS, and is much much symmetric around 50% AT content than the synonymous SFS. This is due to selection; many amino acid changes are likely sufficiently deleterious, even across the widely varying variants of each gene, that they are effectively purged from the population to a frequency far below the ∼ 1% sensitivity provided by the quasi-random HLII dataset. Sites at intermediate frequency are most likely selected on differently in different genomic backgrounds, but a small fraction being very close to neutral is not ruled out: studying subgroups of the population could provide information on this. It is important to note that interpreting the parts of the spectrum other than monomorphic and singleton sites as arising mostly from mutation selection balance runs into the same problem as trying to force-fit the synonymous SFS this way: an unreasonably narrow range of small selection coefficients would be required. A more plausible scenario is selection that varies with genomic background, ecological factors, and time.

**Figure S19:**
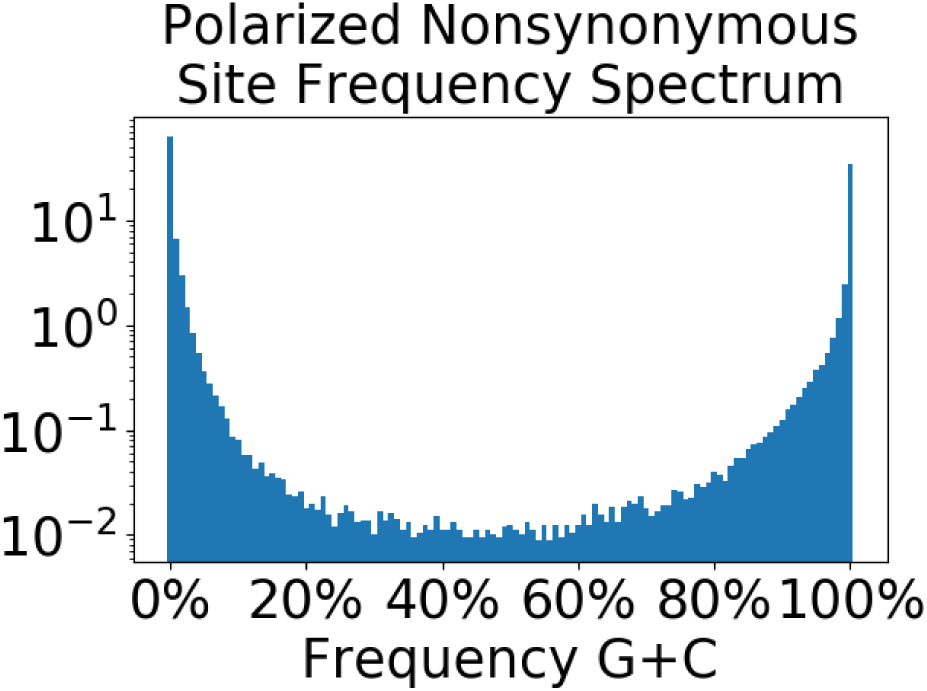
The AT/GC polarized nonsynonymous SFS for the quasi-random dataset, constructed from the second letter of each codon in the core genome, downsampled to 120 cells. This is much more symmetric in the midrange of frequencies than the synonymous SFS, and has large peaks at the extremes: apparently monomorphic sites with observed *f_GC_* =0% and 100%. In the downsampled data (shown), 22.9% of second letter sites are polymorphic (while in the original data, 25.7% of second letter sites are polymorphic).

#### F. Coalescent time distributions

The site frequency spectra reflects the whole phylogeny but is very “noisy” because it focuses on single sites. A complementary quantity is to focus on pairs of cells – thus bits of the phylogeny at a time – but look at the divergences between a longer segment of the genomes than a single site. If this is short enough that recombination has not broken it up, the divergence distribution reflects the distribution of asexual coalescent times – times to common ancestor, *T* – in a way that does not depend on the recombination process.

For rapid hitchhiking, pairs of cells only coalesce to a common ancestor a time *T* in the past if there was a hitchhiking event of size *h* from which *both* of the pair descended: hence a factor of *h*^2^. In steady state, the rate of coalescence per time in the past is constant at rate 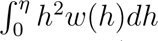 yielding an exponential distribution of coalescent times: *He*^−*HT*^ *dT*, with mean 1*/H*. This is exactly the same form as drift.

For more complex coalescent processes – already the time to common ancestor of 3 cells – the behavior for drift and hitchhiking with a broad range of *h* are different. The SFS of a sample of *S* cells is determined by the coalescent properties – i.e. the phylogenetic trees – of the whole set of *S* cells.

For a uniform expansion all the coalescences occurred at the same time *T_c_* ≈ log(*N*)*/γ* ≈ 62*/γ* in the past so the distribution is approximately a delta-function at that time. (There are slight corrections to this due to when mutations occurred in the very early Luria-Delbrü ck regime, but these can be ignored).

##### F.1. Simulating synonymous divergence distributions

We compare several theoretical coalescent time distributions to the data by simulating the synonymous gene pairwise divergence distributions yielded by each. The transformation between the coalescent time distribution and the pairwise divergence distribution involves two steps: transforming the coalescent time to the probability that a given site differs between the cell pair, and accounting for sampling noise due to finite gene length.

To transform the coalescent time to the probability that the two cells differ at any given site, we use a bi-allelic model. We compare to the data using twofold degenerate sites and fourfold degenerate sites polarized to A/T vs G/C. The two types of sites are weighted according to their heterozygosities in order to achieve a proxy for divergence time, (the typical amount of time for a cell pair to accrue a mutation at a twofold degenerate site is longer than the typical amount of time for a cell pair to accrue a mutation at a polarized fourfold degenerate site) resulting in one twofold-degenerate site having about 1.48 times the weight of one fourfold degenerate site. The mutation rate ratio measured in Section B. is input into the biallelec model.

The probability that a single site has not been mutated during the time, *T*, from its common ancestor with another cell, decays exponentially, hence its memory of the common ancestor similarly. Averaging over the ancestral state assuming mutational equilibrium gives the probability that a site differs between a cell pair

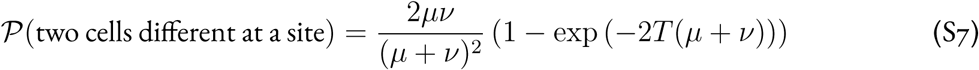

where have measured mutation rates per time rather than per generation. This immediately enables sampling over an assumed distribution of coalescent times for the divergences of genes, adding up many sites.

To combine both twofold and fourfold degenerate sites, we weight twofold degenerate sites by the ratio of fourfold to twofold site heterozygosity (as described above) and the fraction of synonymous sites at non-sixfold-degenerate codons that are twofold, *f*_2_ ≈ 0.62. As noted, we use that the mutation rates for polarized fourfold degenerate sites that are each about 1.5 times as high as for twofold degenerate sites, *µ*_4_ ≈ 1.5 · *µ*_2_ and *ν*_4_ ≈ 1.5 · *ν*_2_ (where subscripts 2 and 4 denote twofold and polarized fourfold degenerate sites, respectively). The distribution for P(different|*T*) (the probability that two sites differ in allele as a function of divergence time *T*) is obtained using the distribution for the coalescent time *T* for each theoretical model. The simulated divergences at individual genes are obtained by binomial sampling given the infinite-length gene divergences, and plotted in Figure 5 (b) in the main text.

In addition to the exponential and delta-function distributions of coalescent times, we also simulate a model motivated by the observed decay of linkage correlations within genes, by close to 50% by some measures. This implies that some recombination events have broken up many of the genes since their common ancestor. A crude model is to treat each gene as a sum of two half genes with independent phylogenies. The distance distribution would then be roughly like that from a coalescent time distribution of the form (4*T/T* ^2^) exp(−2*T/T_c_*) which has mean coalescent time *T_c_*. The simulations of divergence distributions for this, Figure 5 (b) in the main text (green curve), are closer to the data than the other forms.

##### F.2. What gives rise to the apparent coalescence distribution?

The two half-gene model, while it looks quite like the data, may be problematic. First, it is not clear that the linkage correlations decay enough to justify convolving two fully-independent halves. Even if they do for typical coalescent times, for short coalescent times, which is where the standard exponential form is worst, genes would be less likely to have had a recombination event and thus smaller divergences would be less suppressed. Analysis of linkage correlations within the hitchhiking models could help resolve whether the various quantities measured are consistent with the simplest models.

There are other possible resolutions of the puzzle of anomalously few low divergence pairs of cells. Conventional coalescent theory is usually applied to sexual populations where different chromosomes, and far away parts of the same chromosome, can have very different histories: thus the assumption that what is observed averages over a huge number of the statistically similar histories is reasonable. For an asexual population at the opposite extreme, there is only one phylogeny and thus only one pairwise coalescence distribution. In asexual populations, because there have been a large number of coalescent events in the recent past, the small *T* part of the coalescence distribution should be well sampled. However the long time part will not be: there is a single longest coalescent time and a substantial fraction of pairs will coalesce then. Thus the large divergence part of the distribution will not be well approximated by the average over phylogenies.

For genome-scale hitchhiking with abundant large *h* values, the situation could be worse. If hitchhiking is mostly asexual, even when there is a lot of recombination between hitchhiking events there is still only one set of hitchhiking events. The rates of coalescence are proportional to *h*^2^ and thus dominated by large *h*. However, the largest *h* that is likely to have happened more recently than a time *T* in the past is typically roughly *η*(*T*) given by 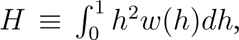 any *h* larger than *η* is not likely to have happened. This means that typically the effective rate of coalescences up to time *T* is crudely 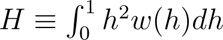 which can be much smaller than the full integral of *H*. If the hitchhiking in HLII were primarily genome wide, then HLII acts like just one “sample” and so typically will not have had many coalescences in the recent past. (Of course, in some other universe, there would have been a large hitchhiking event very recently, accounting for many pairs, thereby filling in some of the hole in the coalescence distribution). Note that the absence of large tight clusters does not completely rule out relatively recent large genomescale hitchhiking events: it is possible that even recent asexual blooms may have died down and left no clear genome-wide trace, even though they brought some synonymous sites up to high frequency. Yet, it is unclear whether such transient events could give rise to the observed SFS. Furthermore, if large hitchhiking events had not occurred since the ancient bloom, the SFS of the quasi-equidistant subset would be very different.

For genic-scale hitchhiking, each section of the genome of lengths of typical recombination events is roughly independent so that the averaging over the genome would effectively average over the hitchhiking distribution and the above concerns would not apply.

Another way in which recent coalescent events might be suppressed is slow blooms, in which many mutations accumulate during the rise of the bloom. These would suppress the number of low divergence pairs of genes. However, just as for a single expansion, they would also tend to squeeze the SFS in from the edges, so it is not clear if this scenario is consistent with the data.

Another possible scenario is that the important changes in selection are not due a sudden genetic change, but instead a sudden ecological change – perhaps a new phage to which some fraction of the population is less susceptible – which raises the abundance of a whole range of genomes, perhaps all those that have some set of variants of one phage-related gene. This would affect the SFS – although in somewhat a different way than hitchhiking from a single cell –of sites that are linked to the favored gene. But because of the large number of ancestors involved, it would not give rise to coalescent events.

This brings us back to the question of whether the dominant driving events are primarily asexual. We have argued that the shortage of asexual blooms in the recent past that would have been expected if the genome-scale events dominated, and the good fit to the genic-scale hitchhiking model, strongly suggest that the latter dominate and the concerns above may be moot.

Genic-scale hitchhiking by a segment being transferred around with a selected genetic element will cause decay of linkage between sites at different distances away. Analyzing this process would potential enable disentangling linkage correlations caused by this process, versus being caused by closer-to-neutral recombination between substantial hitchhiking events.

What combination of these or other scenarios might better explain the data – and suggest what kind of data would be most informative moving forwards – we leave as interesting theoretical and data analysis challenges.

### 5. Clonal Backbones and Recombination Events

In this section we describe how we directly identify recombination events to and from closely-related groups of cells. Most of this analysis focuses on the C1 cluster from the deep clustered dataset (Section 1.). We also perform a similar analysis on additional clusters within the quasi-random dataset.

#### A. Clonal backbones and recombination events in tight clusters

Because of the high diversity within HLII, it is difficult to identify recombination events directly. However, within tight clusters which can still share a substantial clonal backbone, recombination events with cells outside of the cluster will appear as regions of unusually high divergence. This allows one to directly identify recombination events and estimate evolutionary parameters, such as the typical length of transferred segments and the ratio of mutation and recombination rate. Different approaches for distinguishing between the clonal backbone and recombined genome segments have been proposed (*18*, *19*). Here, we use a statistical approach to estimate the range of divergences within the clonal backbone of SAGs from tight clusters and identify recombined segments whose divergences are inconsistent with this range.

##### A.1. Estimation of clonal backbone divergence and fraction of genome recombined for closely-related cell pairs

We used the whole-genome alignments provided by Kashtan et al. to estimate the clonal backbone divergence within the deep clustered dataset. Within the clonal backbone, we assumed mutations accumulate following a Poisson processes. For a given cell pair, the number of SNPs within the clonal backbone is also given by a Poisson process with a rate *d* given by the clonal divergence. Because different pairs of cells have different divergence times, *d* will vary between pairs. However, different genome segments within the clonal backbone of a given pair should have the same rate. We verified this by plotting the distribution of the lengths of identical genome segments for each pair of cells from the C1 cluster in the deep clustered dataset. Under the Poisson model, the distribution of the lengths of identical segments *ℓ* is exponential *P* (*ℓ*) ∝ *e*^−*dℓ*^. We found the data was consistent with these expectations (Figure S20).

**Figure S20:**
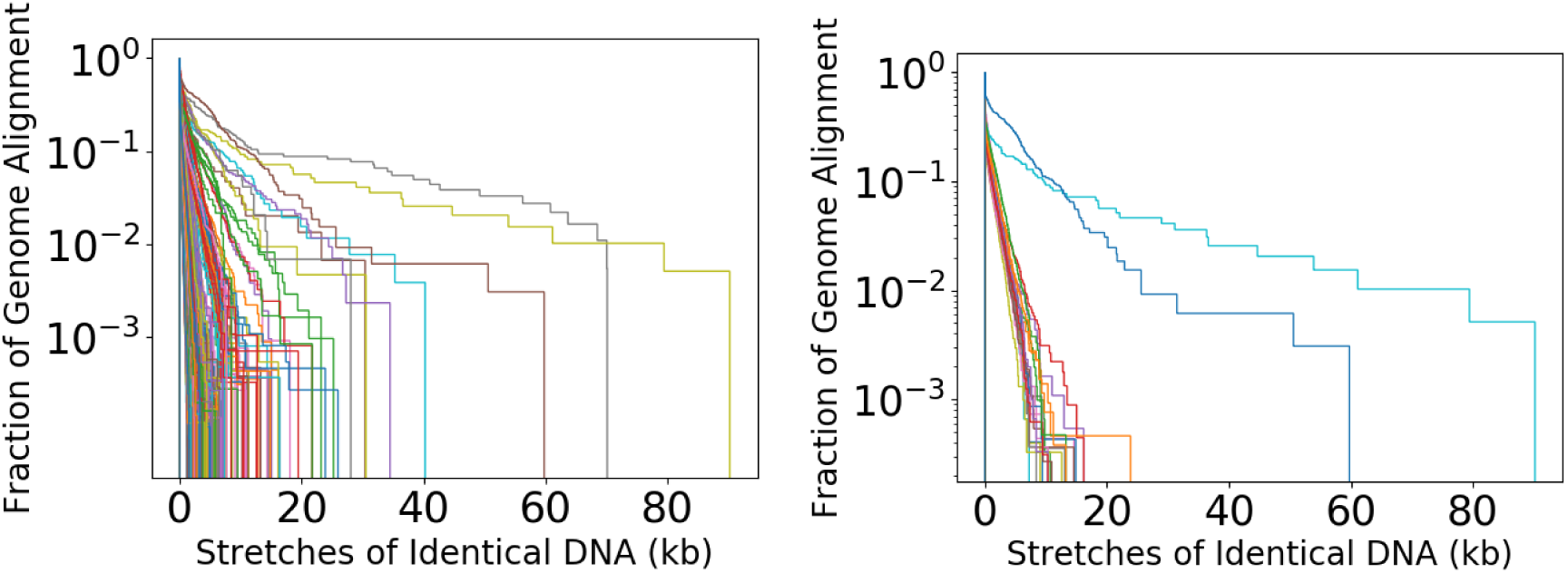
Distribution of the lengths of identical stretches of DNA for cell pairs, using Kashtan’s provided alignments, for C1 (**left**) and a 7-cell close clique within C1 (**right**). Note that the tails of the distributions are close to exponential as expected for a Poisson mutation model with a time to common ancestor for each cell pair that does not vary along the genome.

To estimate the divergence *d* we partitioned the genome into segments of 1 kb length and counted the number of SNPs per kb, for each cell pair. The distribution of the number of SNPs per kb within low-divergence segments was well-approximated by a Poisson distribution (Figure S21). We use the fraction of identical kb, *f*_0_, for each pair to estimate the clonal backbone divergence using *λ* ≡ *dℓ* = − ln(*f*_0_), where *ℓ* = 1 kb. The divergences obtained using the method described above were similar to those from exponential fits to the distribution of lengths of identical genome segments from Figure S20. These results are thus consistent with the low-divergence regions forming a single-clonal backbone.

The clonal backbone divergence was then used to estimate *f_rec_*, the fraction of the genome that was recombined, for each cell pair. For closely-related cell pairs, recombination will very likely introduce segments with much higher divergence (Figure S22). We used the fitted Poisson distribution for 1 kb segments as a null model and labeled segments as recombined using a p-value of *p* = 0.01. We verified that our results are robust to the change of *p* by varying it from 10^−3^ to 10^−1^ and found the average *f_rec_* varied by less than 2 percent. The number of SNPs contained in recombined segments as defined above increased linearly with the inferred clonal backbone divergence between pairs, consistent with a roughly constant rate of recombination (Figure S23). The inferred *d* and *f_rec_*for each cell pair in C1 are shown in Figure 2 (c) in the main text.

**Figure S21:**
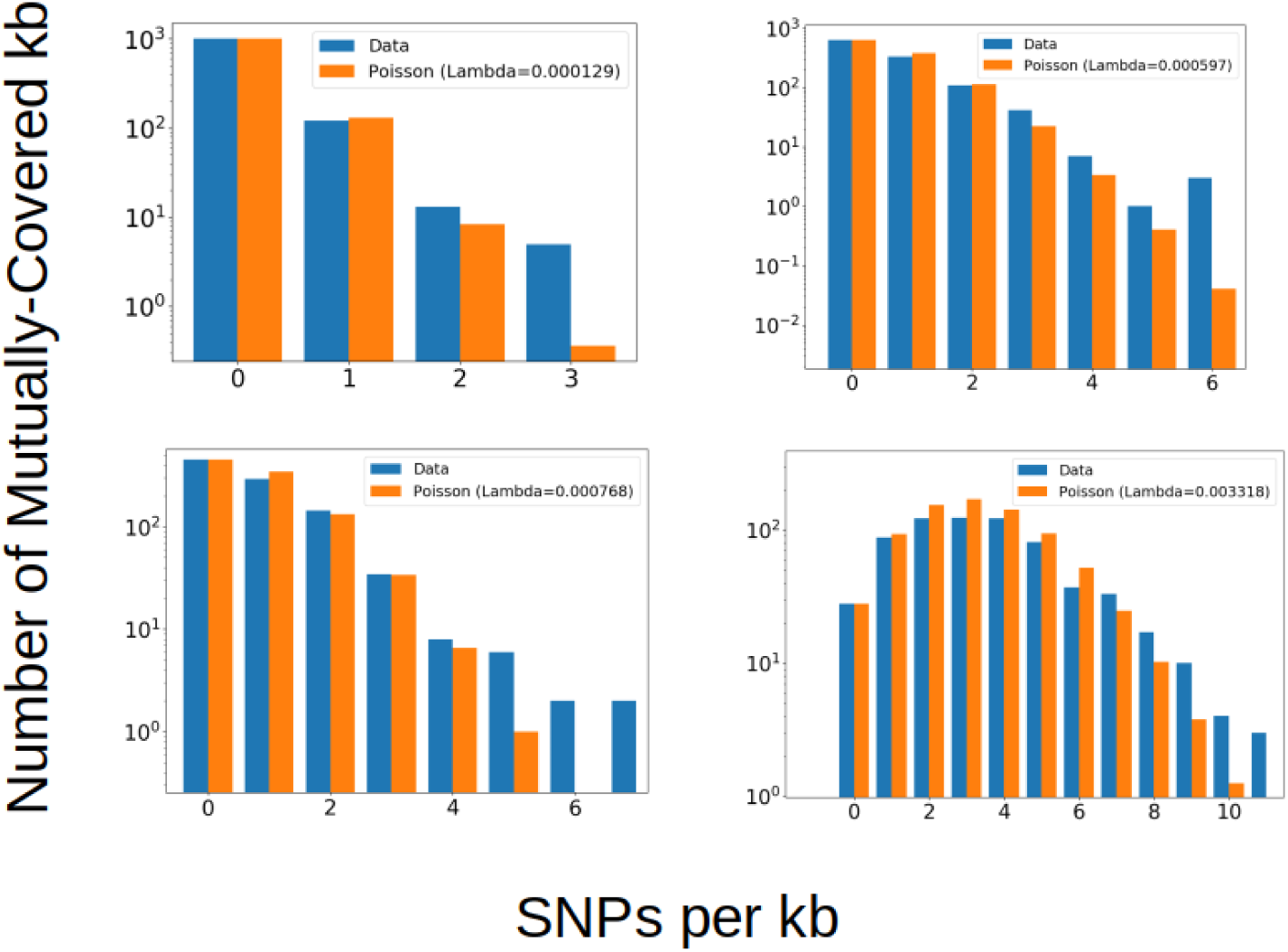
The low-divergence regions of closely-related cell pairs from the deep clustered dataset agree well with a Poisson mutation model, as expected if pairs share a clonal backbone. Each panel shows the distribution of SNPs per kb using the Kashtan-provided reference-guided alignment for a given pair of cells in the cluster C1 (blue). The Poisson distributions showing the expectations from the clonal backbone divergence are shown in orange for comparison. The pairs chosen include one of the most closely-related cell pairs sequenced to date (upper left), which has no identified recombined segments in the core genome (**top left**) along with three other pairs (**top right** and **bottom**) chosen for illustration. The Poisson distributions are fitted using the fraction of mutually-covered kb (requiring at least 80% of the kb be mutually covered) that are identical, for each cell pair.

**Figure S22:**
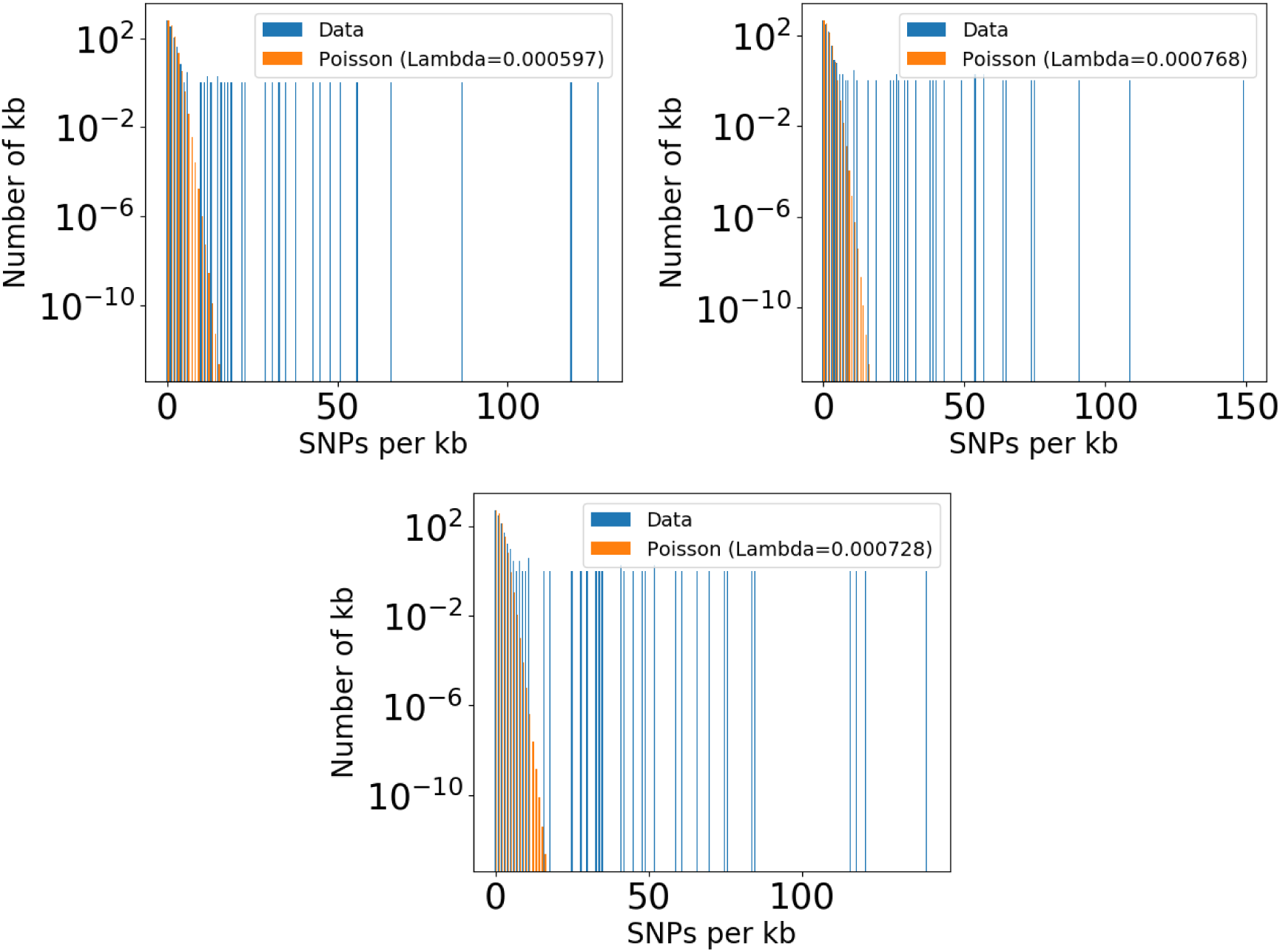
The full distributions of SNPs per kb for three of the four cell pairs shown in Figure S21, excluding the most closely related pair. High-divergence kb that have implausibly low probability of occurring in the asexual Poisson model are clear for each pair, and contribute to the fraction of genome recombined estimated for that pair. The long tail of each distribution that is inconsistent with the asexual model is used to estimate the fraction of recombined genome for each pair (see text).

**Figure S23:**
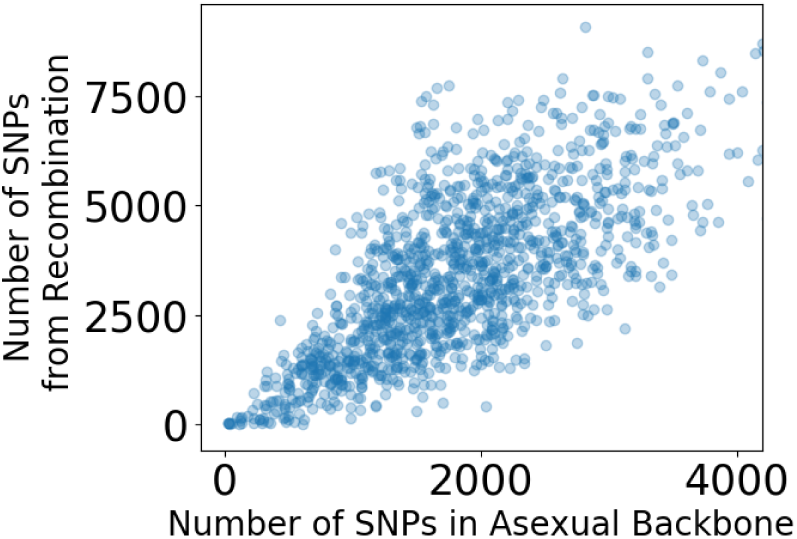
The estimated number of SNPs introduced by high divergence recombination events compared to the number of SNPs in the clonal backbone for cell pairs within C1. Figure includes most C1 pairs. A small number of outlier points with up to 10,000 SNPs from recombination and 6,000 SNPs on their asexual backbone are excluded. Outlier points are consistent with the trend in the bulk of the data shown here.

##### A.2. Uncertainty in calculating clonal backbone divergence

Our estimates of *f_rec_* above Section A. A.1. assume that the divergence in recombined segments is higher than in the clonal backbone. It is possible that the low-divergence regions of the genome have been subject to recombination between very closely-related cells that share the same clonal backbone. Since our method would not detect such events, they would lead to an underestimate in *f_rec_*. However, in order for a significant fraction of the low-divergence regions of genome to have been subject to gene exchange, a marked increase in the rate of recombination as a function of donor-recipient divergence would be required at very low divergences, especially because so few cell pair exist at very low divergences relative to cell pairs with typical HLII divergences. Simply to achieve a few percent of the genome recombined with closely-related cells would require over a factor of 100 increase in recombination rate at low divergence.

This is because, for a given cell, the fraction of other HLII cells that are within a few percent asexual divergence is typically less than 1%. Figure 2 (b) in the main text does not provide evidence for this scenario, instead showing a relatively uniform rate of recombination for divergences less than 5%.

##### A.3. Identification of high-divergence recombination events among closely-related cells

We used two methods to group recombined genome segments from the deep clustered dataset into distinct *recombination events*. Within closely-related cliques we use the whole-genome alignments from Kashtan et al. and identify recombined regions as follows. First, we cluster each 1 kb segment using an agglomerative clustering algorithm using average cluster linkage, as implemented in SciPy (*20*). Next, we identified distinct allelic clusters within each segment using a 1% divergence cutoff. This groups all of the sequences into clusters such that the distance between clusters (defined by the average cluster linkage) is *>* 1%. We selected those segments that contained at least two clusters by this definition. We then examine the alignments of these segments and manually identify recombination breakpoints and exclude segments that appear to be caused by alignment errors. An example of recombined segments identified in this way are shown in Figure 2 (a) in the main text and Figure S24. All regions of the genome, including flexible genes, are included in this analysis. Within each close clique, most island regions (in which many of the flexible genes occur) have not undergone multiple recombination events and individual events can still be distinguished.

**Figure S24:**
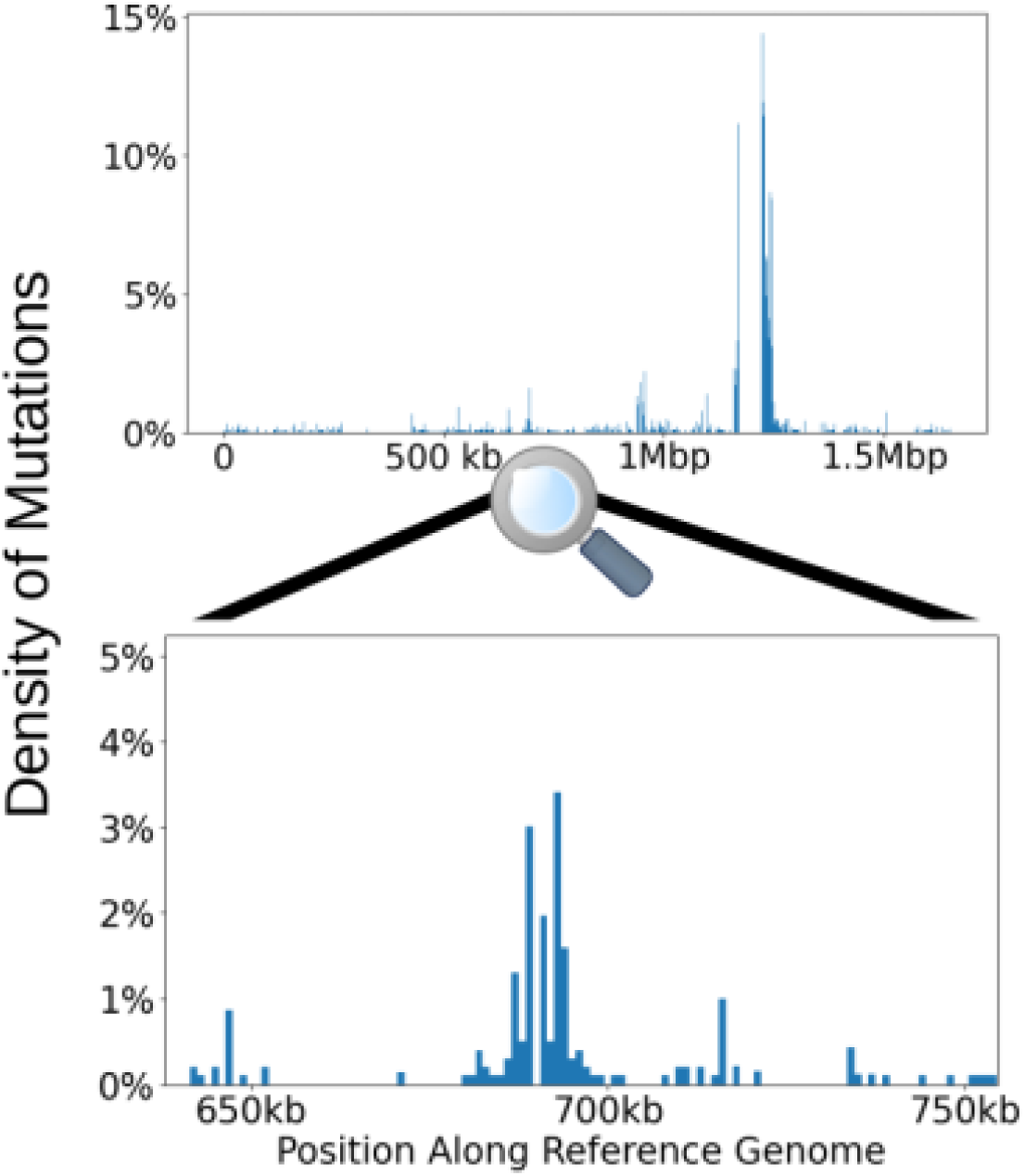
The divergence per kb along the genome of a closely-related cell pair in C1, using the whole-genome alignment from (*2*). Distinct high-divergence events are visible atop a low divergence clonal background. Some gaps in the divergence distribution indicate no coverage (for example, at the location of the hypervariable island 4).

We used a similar procedure to identify recombination events in the entire deep clustered dataset, but here the choice of divergence cutoff to define allelic clusters is more difficult. We were only able to distinguish recombination events in the core genome, because the highly-variable island regions of the genome often had multiple overlapping events and many highly-diverged alleles. Figure S25 shows that the number of core kb with more than 1 cluster for different divergence cutoffs roughly flattens out for divergence cutoffs larger than 1%. We interpret this as evidence that 1% is the typical clonal backbone divergence scale for C1. About half to two thirds of kb in the core genome have only 1 cluster with a 1% cutoff; our interpretation is that these single-cluster kb have not been subject to a successful recombination event in the history of any of the C1 cells in the deep clustered dataset. This result was fairly robust to choice of cluster linkage (average, single, complete). Based on this result we used a conservative cutoff of 2% to define recombined allelic clusters. Next, we looked for groups of at least 2 consecutive core genes that have the same clustering structure at 2% divergence and grouped these into distinct recombination events. We confirmed the edge cases by hand.

**Figure S25:**
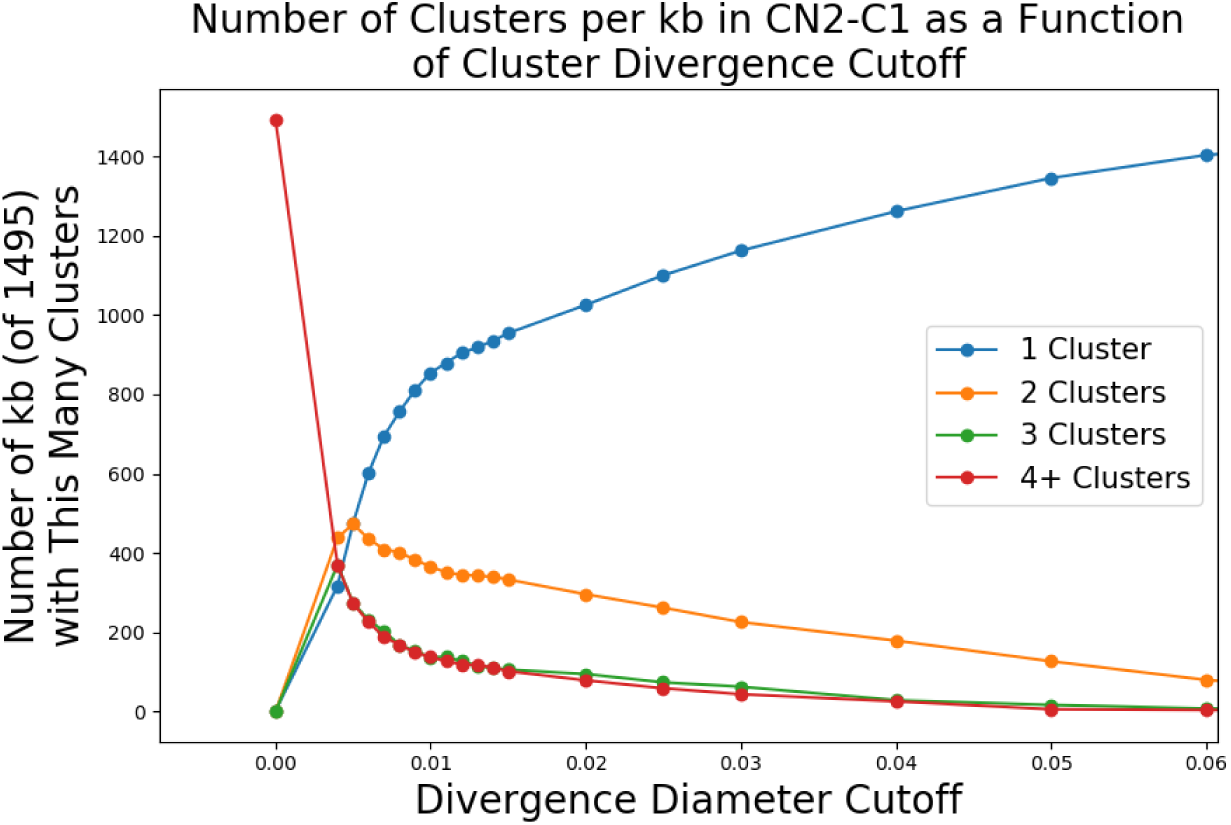
The number of clusters for each core gene within C1 as a function of the divergence cutoff used to define clusters. Pairwise distances were calculated using the whole-genome alignment from (*2*) and were clustered using the average cluster-linkage algorithm implemented in SciPy. Curves show the number of core genes (out of a total of 1495) that contained 1 (blue), 2 (orange), 3 (green), and 4 or more (red) clusters. Similar results were obtained using other cluster-linkage algorithms (data not shown). Note the sharp change in slope for the 1-cluster curve shown in blue at around 1% divergence cutoff.

Using the procedure described above, we found 21 events in closely-related cliques within the deep clustered dataset, and 45 additional events within the core genome of the whole deep clustered dataset. We used these events to estimate the nucleotide (Figure 2 (b) in the main text) and synonymous (Figure S26) divergences of recombined segments. We also examined the lengths of recombination events. Figure S27 shows the distribution of estimated lengths of high-divergence core genome transfer events found in the close cluster C1 and the 6 close cliques of CN2, suggesting a rough range of 1-20 kb and a typical length of about 10 kb. Note that because of the difficulty in identifying such recombination events directly, we did not use this method to estimate the fraction of genome recombined per cell.

**Figure S26:**
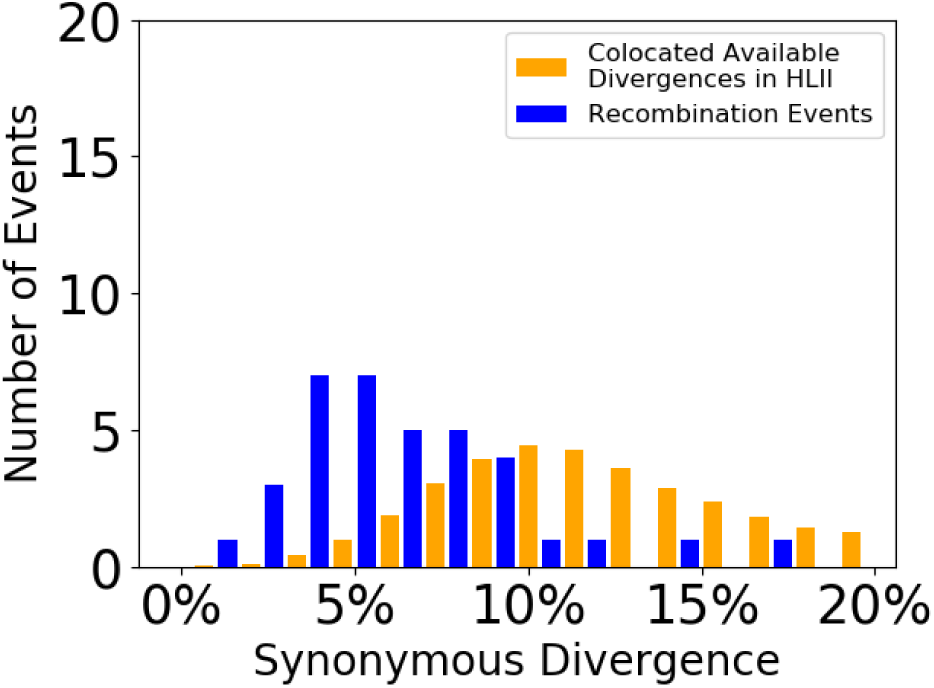
Synonymous divergences of distinct high-divergence recombination events identified within core genome regions of C1 cells. Compare to all-nucleotide divergence shown in Figure 2 (b) in the main text.

**Figure S27:**
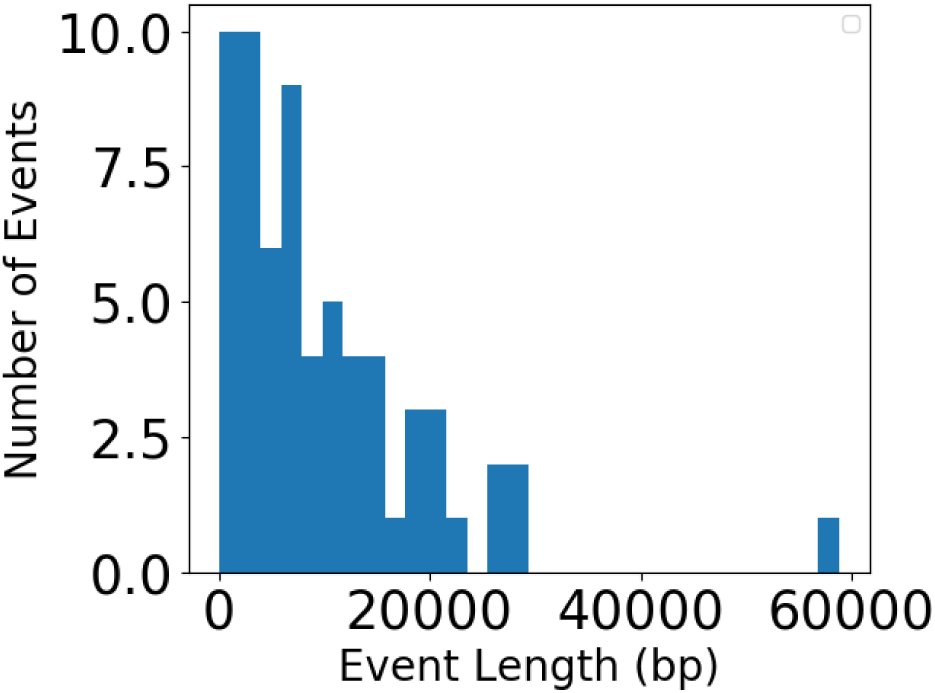
Estimated lengths of transfer events found in C1 and the 6 close cliques of CN2.

##### A.4. Identifying donors of recombination events identified in C1 SAGs

We searched for genome segments transferred from other quasirandom SAGs *into* C1. High divergence recombined regions identified in C1 SAGs from the deep clustered dataset (Section A.3.) are aligned using BLAST against a database of 489 *Prochlorococcus* cells from the Berube et al. dataset (*1*), which includes the quasirandom HLII sample. This is done to roughly gauge whether a donor cell can be identified. We find best hit putative donors with 1% divergence or less for 13 of 42 recombination events identified (see main text) in the core genome of C1. Here, we will use the term “putative donor” to refer to the best match SAG identified within 1% divergence of the recombination event. About 1*/*3 of putative donors are from the Pacific, despite the fact that all C1 SAGs are from the BATS sampling site in the Atlantic. All identified putative donors of core events are in HLII, although a putative event 5 genes in length in the flexible genome (not included in the 42 core events) has a ∼ 99.5% identical match in HLI.

#### B. Recombination events with tight cluster donors

##### B.1. Identifying deep clustered alleles

We quantify the extent to which cells in the closely-related groups C1, CN2-C2, CN2-C3, CN2-C4, and CN2-C5 identified by Kashtan et al. (*2*) act as donors in core homologous recombination events. We refer to these clusters as C1-C5 throughout unless otherwise noted. We first identify core genes at which each of C1-C5 forms a distinct cluster. To do this, we consider only genes at which the divergence diameter of each cluster C1-C5 is less than the minimum divergence between a cell in that cluster and any cell in one of the 4 other clusters (considering only the C1-C5 cells from the deep clustered dataset). We identify 453 core genes that meet these criteria. We then use the divergences of alleles from the quasirandom dataset with the alleles from the C1-C5 clusters defined above to identify alleles that were likely transferred from C1-C5 into other HLII cells represented in the quasi-random dataset. Specifically, each allele in the quasi-random dataset is assigned to one of the C1-C5 clusters if the maximum divergence between it and all other sequences from the cluster is less than or equal to the diameter of the cluster.

##### B.2. Transfers of deep clustered alleles into quasi-random cells

We find seven SAGs from the HLII quasi-random dataset that cluster with one of C1-C5 at a majority of the 453 genes examined. Note that the clusters in these cells are consistent across several hundred genes, which provides stronger support for a shared clonal backbone than clustering based on ITS (*2*). Of the seven SAGs, 3 were assigned to C1, 1 to C2, 2 to C3, and 1 to C5. All 7 of these cells are from the same sample taken from BATS at 10 m depth on 7/14/2009 (*1*). Note that the C1-C5 cells in the deep clustered dataset are all from the BATS sampling site with sampling times ranging from November 2008-April 2009 (*2*).

To identify transfers from the C1-C5 clusters into other SAGs, we exclude the 7 C1-C5 SAGs and analyze the remaining 160 quasi-random SAGs. We find about 13% of the 453 examined *genes* have at least one quasi-random SAG whose allele clusters with one of C1-C5. We refer to the alleles in the quasi-random SAGs that cluster with C1-C5 as *C1-C5 alleles*. The genes that have at least 1 cell clustering with C1-C5 have on average 3 such cells. In total, about half of the 160 quasi-random SAGs contain at least one C1-C5 allele across all 453 genes, and some SAGs contain C1-C5 alleles at up to 12 genes. The distribution of the number of C1-C5 alleles per quasi-random SAG is similar between Atlantic and Pacific SAGs with slight enrichment in Atlantic SAGs (Figure S28). When we divide the Atlantic SAGs into BATS and non-BATS SAGs, we find BATS SAGs have 50-100% more C1-C5 alleles compared to non-BATS Atlantic and Pacific SAGs.

**Figure S28:**
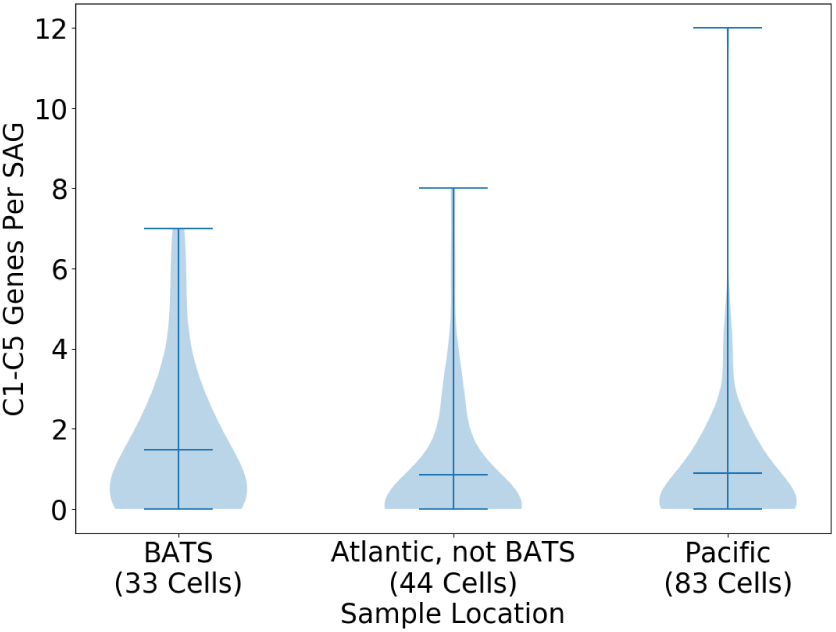
Alleles apparently originating from the deep clusters C1-C5, found at the BATS sampling site in the Atlantic ocean, are found in cells spanning the diversity of HLII, in both the Atlantic and Pacific oceans. Cells from the quasi-random dataset are searched for core gene alleles that cluster with cells from C1, C2, C3, C4, or C5. 453 core genes are used in the search, selected because the clusters C1-C5 form distinct, non-overlapping groups at these genes (see text). For each cell in the quasi-random HLII dataset, excluding the 7 cells that cluster with one of C1-C5 along the entire genome, the number of core genes which cluster with one of C1-C5 is counted and binned by sampling location. Note that the Atlantic and Pacific oceans have highly similar distributions, despite BATS (located in the Atlantic) being slightly enriched for C1-C5 alleles (Section B. B.2.).

##### B.3. Null model for transfers from C1

We compare our results to a simple null model in which transfers of core genes across HLII occur at a single constant rate. Here we focus our analysis on just C1, which is the largest of the CN2 clusters. A simple assumption is that C1 had a constant frequency *f_C_*_1_ within HLII since its most recent common ancestor. We use this assumption and then check the resulting estimates against the data. Within HLII, the time for the core genome to be overwritten by recombination, *T_overwrite_*, is given by *µT_overwrite_* ∼ 3% (Figure 2 (c) in the main text). If recombined segments are drawn randomly from the HLII population, then the fraction of transfers from C1 into any other cells is given by

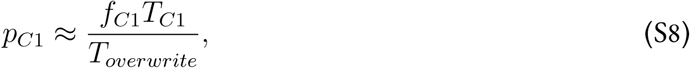

where *T_C_*_1_ is the time since common ancestor of C1, given by *µT_C_*_1_ ≈ 0.3% (Figure 2 (c) in the main text).

We use both ITS data and the quasi-random dataset to estimate the frequency of C1, *f_C_*_1_, in the ocean. While our results show that ITS is a poor predictor of whole-genome divergence at the level of HLII (Figure 2(d) in the main text), it can give a rough estimate of the abundance of tight clusters such as C1. For this analysis, we used a total of 3398 ITS sequences, including 1381 from the BATS site in the Atlantic, obtained from (*2*) and (*1*). We found that the C1 and C2 clusters were not distinguishable based on ITS so we grouped both together. Among the 1381 BATS sequences, about 10% cluster with C1/C2. Three of the 39 SAGs (≈ 8%) from a sample from the BATS site in the quasi-random dataset cluster with C1 at the whole genome level and one SAG in this sample clusters with C2 at the whole genome level. Among the remaining 2017 ITS sequences not from BATS, ITS sequences clustering with C1/C2 are found almost exclusively in one sample, which is labeled as the Westerlies-Gulf Stream Province (located about 570 miles away from the BATS sampling site (Figure S5)) (*1*). At this sample, 10 of 147 ITS sequences (≈ 7%) cluster with C1/C2. Using the results from the Atlantic samples as a rough estimate, we obtain *f_C_*_1_ ≈ 10% (similar to the ∼ 8% obtained using SAGs that cluster with C1 at the whole genome level, from the BATS site).

Substituting the values above into (S8) gives *p_C_*_1_ ∼ 0.01, meaning that we would expect about 1% of core genes in a typical non-C1 HLII cell to have a C1 allele. In the quasi-random dataset we found 0.1% -1% of core genes had a C1 allele –roughly, and perhaps surprisingly, consistent with the null model.

##### B.4. Transfers from other tight clusters into quasi-random cells

To further test the null model proposed above we performed a similar analysis on the tight clusters of closely-related cells in the quasi-random dataset found in Section 2.. We estimate the number of core alleles transferred from the three tight clusters consisting of seven SAGs each (labeled QR-C6-8 in Section 2.) using the same procedure as for the deep clustered alleles described above. To minimize false positives, we only include core genes in which the QR cluster diameters are less than 2% nucleotide divergence. Note that no well-covered core gene has divergence diameter below 5% across the full quasi-random data set, and only 9 well-covered core genes have median cell pair divergence less than 2.5% within the quasi-random sample. We therefore concluded that the gene clusters detected using this method are unlikely to be due to gene conservation and instead represent transfers within the population.

We identified alleles transferred from each of the three QR clusters and compared them to the expectation from our null model as in the C1 case. Unlike before, we do not have complete genomes for the QR clusters so the expected fraction of transferred alleles is

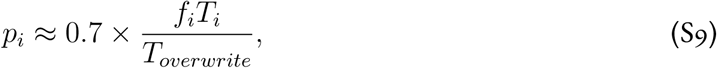

where the extra factor of 0.7 accounts for the typical genome coverage. The results for all four clusters are shown in Table S5. As in the case of C1, we find a remarkable agreement between the fraction of alleles transferred from each cluster and our null model. This results supports our conclusions that there are no significant barriers to gene exchange within HLII.

**Table S5:**
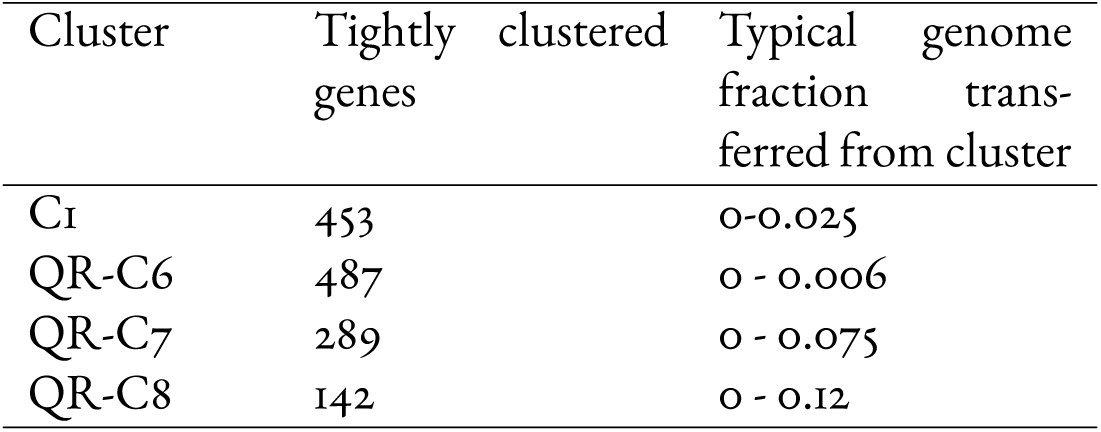
Fraction of gene alleles from each tight cell cluster found in other quasi-random SAGs. The second column shows the number of core HLII genes with a cluster diameter ≤ 2% that were used in the analysis, as explained in the text.

### 6. Statistical Measures of Recombination

In this section we introduce several methods to assess the impact of recombination on HLII evolution. In the first part we review different measures for quantifying recombination rates based on mutation statistics and define the divergence correlations *R* and *R̃* used in the main text. In the second part we analyze divergence correlations within the HLII core genome and discuss two simple scenarios that could explain the main features of the data.

#### A. Correlations along genome: measures of linkage disequilibrium

Correlations along the genome – linkage disequilibrium – can be quantified in various ways. The Pearson correlations, *R* and *R̃*, we have used, examine linkage at the gene scale, while measures such as ⟨*r*^2^⟩, *σ*^2^, and other site-level distance correlations (see Section A. A.1.) examine linkage at the SNP level. The most commonly used linkage correlations, ⟨*r*^2^⟩ and *σ*^2^, are defined as follows:

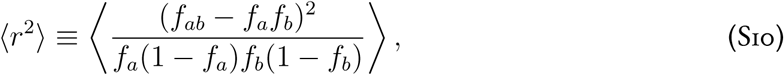

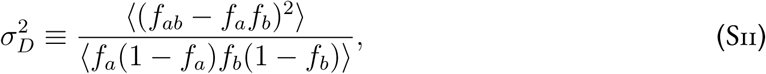

with *f_a_* and *f_b_* the frequencies of one of the alleles at each of two dimorphic sites, with *A* and *B*, respectively, the other alleles; *f_ab_*is the frequency of the combination *ab*. The averages in Equations S10 and S11 are taken over SNP pairs separated by a given distance, *ℓ*, along the genome. ⟨*r*^2^⟩ assigns higher weight to the low frequency sites than σ^2^_*D*_. Both are plotted as a function of separation between SNP pairs in Figures S29 and S30, respectively. The expectation from a random shuffle (”RS”) is also shown. The ran-dom shuffle for ⟨*r*^2^⟩ is 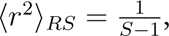 with *S* the number of cells in the sample, here about 80 for HLII due to downsampling to account for site pair coverage. Within the HLII quasi-random sample, the excess linkage above background (for ⟨*r*^2^⟩) decreases by about a factor of two over the length of a typical gene. A similar, though slightly less substantial, decay over a gene length is seen in *σ*^2^_D_.

**Figure S29:**
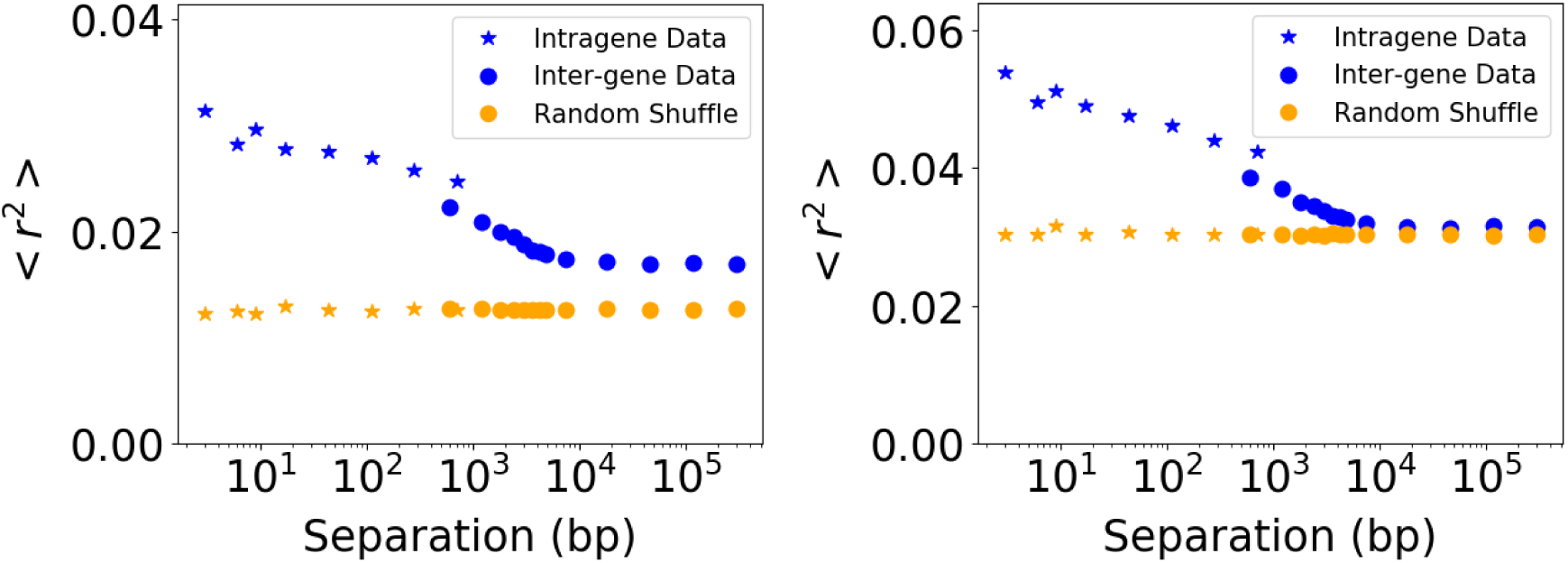
SNP-level lineage measure ⟨*r*^2^⟩ (Equation S10) using twofold degenerate sites at HLII core genes, for the entire quasi-random sample of HLII cells (**left**) and the quasi-equidistant subset of HLII cells chosen to minimize structure at the whole-genome level (**right**).

**Figure S30:**
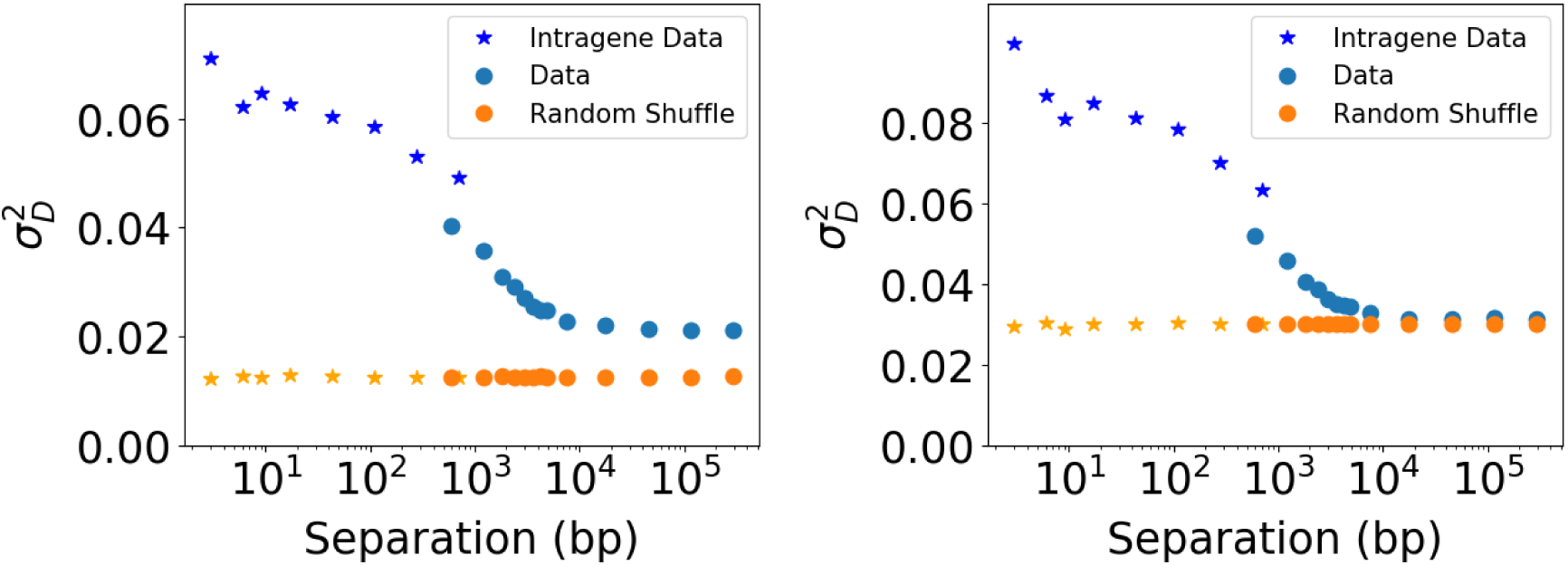
An alternative measure of SNP-level linkage disequilibrium, *σ*^2^ (Equation S11), shows similar lengthscales of decay as ⟨*r*^2^⟩. Panels show results for the entire quasi-random HLII sample (**left**) and the quasi-equidistant subset (**right**) separately. Core dimorphic twofold degenerate sites were used to calculate *σ*^2^_D_ for both datasets.

##### A.1. Divergence-based measures of linkage

The measures of linkage we have focused on in the text are based on divergences between cell pairs. Define *d_i_*(*X, Y*) to be zero if cells *X* and *Y* have the same nucleotide at site *i*, and one if they are different. The heterozygosity at site *i* – the probability that two cells will differ at that site – is just the average over pairs: *d̄_i_* ≡ ⟨*d_i_*(*X, Y*)⟩*_XY_*. The average heterozygosity over a set of sites studied is *π* ≡ ⟨*d̄_i_*⟩*_i_*. The divergence correlation function between two sites, *i* and *j*, is

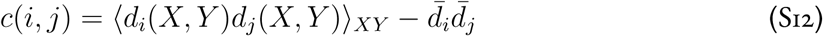

and the average correlation function of sites a distance *ℓ* apart along the genome is *C*(*ℓ*) = ⟨*c*(*i, i* + *ℓ*)⟩*_i_*. Kussell et al. (*21*) use a slight variation of this correlation function to measure linkage disequilibrium within genes – not on longer scales – as a function of *ℓ*, averaging over genes and normalizing by the average heterozygosity, calling this *P* (*ℓ*). The combination of the remarkably well-conserved gene order within HLII (Figure S7), and the use of SAGs allows us to calculate the linkage decrease over much longer distances compared to their study.

If each site has only two alleles – *A* and *a* at site 1, *B* and *b* at site 2 – the average distance at site 1 is *d̄*_1_ = 2*f_a_f_A_* = 2*f_a_*(1 − *f_a_*) (up to the difference between including a cell and itself which does not matter substantially for a large sample) and similarly for site 2. The first term in the correlation function is the probability that *both* sites differ between a pair of cells: *P*_2_(*i, j*) = 2(*f_ab_f_AB_* + *f_Ab_f_aB_*). In figure Figure S31, the two parts of the correlation function averaged over pairs of sites a distance *ℓ* apart are plotted: *P*_2_(*ℓ*) (blue points) and ⟨*d̄_i_d̄_i_*_+_*_ℓ_*⟩*_i_* (estimated by a random shuffle of cell labels, orange points), both normalized by *π*^2^. The ⟨*d̄d̄*⟩ − *π*^2^ part probes the *frequency-frequency* correlations and the difference between these – blue minus orange – is the divergence correlation function. As both measures are expected – at least naively – to go to *π*^2^ at large separations, the figure shows both these with *π*^2^ subtracted: for the entire quasi-random HLII dataset (left) and the quasi-equidistant subset (right). Note again the substantial decay within a gene. The residual correlation at long distances – the difference between the blue and orange – should approach the fractional variance of the genome-wide divergence in the cell group: (*std/mean*)^2^ which is (1.7%*/*7.3*/*%)^2^ ^∼^= 0.05 for HLII and (0.5%*/*5.8%)^2^ ^∼^= 0.007 for the quasiequidistant group. Note that the residual value of the random shuffle (orange dots) is ≈ 1*/S̃* with *S̃* the effective sample size. This is because if one cell has, e.g., an *A* at a site, the other cell is slightly more likely to have an *a*. The effective sample size, *S̃*, is the sample size, *S*, reduced by the probability that a cell is covered at both sites: with average coverage of 0.7, *S̃* ≈ 0.5*S*. The number of cells in the quasi-equidistant subset is about a quarter of the total, accounting for the higher large-separation values of the data.

**Figure S31:**
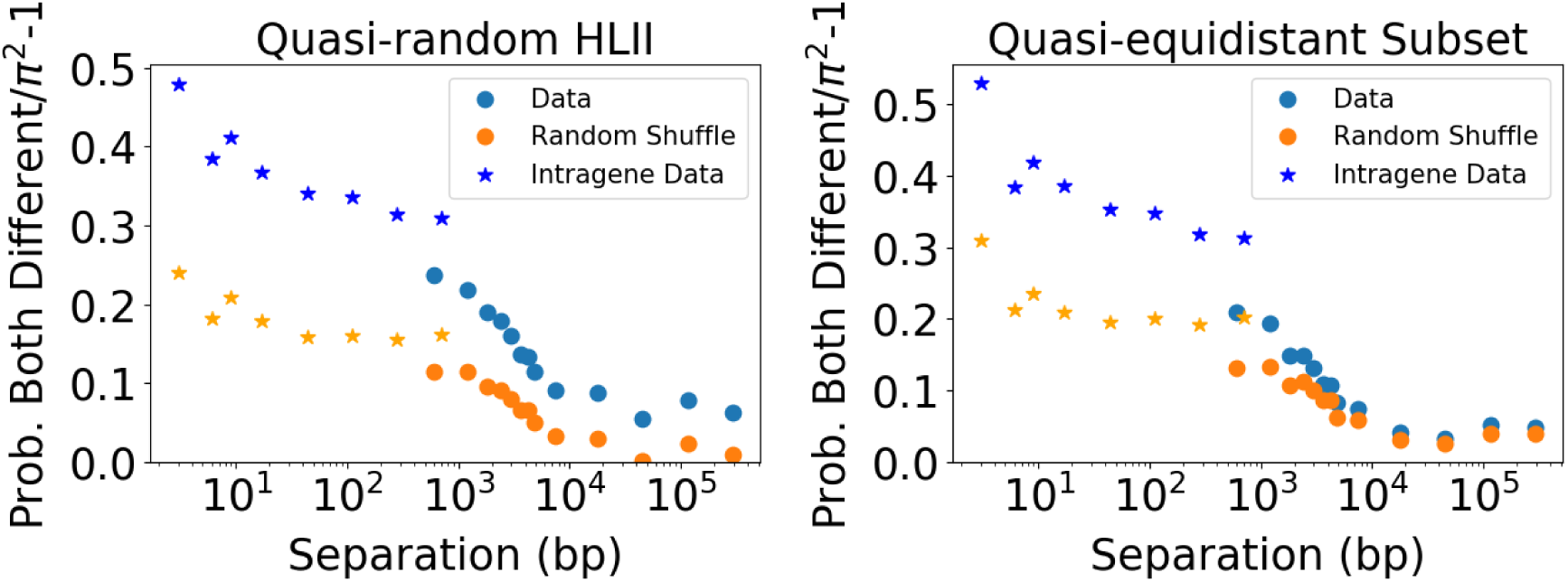
The probability that both sites of a pair separated by distance *l* are different, *P*_2_(*l*), normalized by *π*^2^, less 1, using twofold degenerate core sites for the entire HLII quasi-random sample (**left**) and the quasi-equidistant subset (**right**) is shown in blue. The random shuffle null (orange) is ⟨*d̄_i_d̄_i_*_+_*_l_*⟩*_i_*, the expectation in the absence of any linkage (i.e. where each site is independent). The difference between the blue and orange curves is the normalized divergence-divergence correlation function. The value of the orange data at long separations along the genome is set by 1*/S̃*, with *S̃* the number of cells in the set (or subset) that are covered at both sites: ^∼^= 0.01 for HLII and ^∼^= 0.03 for quasi-equidistant group.

Most of our analyses involve distance between genes rather than single sites. The distance between a pair o<u>f</u> cells at gene *G* with *n_sites_*(*G*) is *D_G_*(*X, Y*) = ∑*_i_*_∈_*_G_ d_i_*(*X, Y*)*/n_sites_*(*G*), with average over cell pairs *D_G_*. The Pearson correlation between genes *G*1 and *G*2 that we analyze is

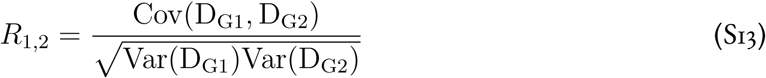

which involves the sum over the correlations between divergence at all sites *i* in *G*1 and *j* in *G*2. The numerator of this is proportional to ∑*_i_*_∈_*_G_*_1_ ∑*_j_*_∈_*_G_*_2_ *c*(*i, j*). If the variations in the variance of divergences from gene to gene are not large then averaging the numerator and denominator separately over pairs of genes is not much different, and the averaged divergence correlation function would behave similarly to *R*. This is a reasonable approximation to the data as the distribution of cell pair divergence variances across genes looks largely tightly peaked: across genes (using all HLII, twofold+fourfold sites), the mean variance is 0.004 and the standard dev of variances is 0.002. Thus *R* is expected to behave similarly to the mean correlation function.

The scaled correlation, *R̃*, which we also analyze, is the Pearson correlation of the gene divergence scaled by the genome-wide average divergence for the cell pair: comparing cells *X* and *Y*, *D̃_G_*_1_(*X, Y*) = *D_G_*_1_(*X, Y*)*/*⟨*D_G_*(*X, Y*)⟩*_G_*. Note that if the variances of cell pair divergences were approximately the same across genes then *R̃* averaged over all gene pairs would be equal to zero, by definition. As noted above, this scenario is well-approximated in the data. In Figure 3 (e) in the main text, we plot the average *R̃* value for various gene-pair-separation bins. The furthest separation bin contains many gene pairs and should be close to zero as it indeed is.

#### B. Statistics of gene-level divergence correlations

Comparisons of the Pearson correlations of the unscaled (*R*) and scaled (*R̃*) cell-pair divergences for pairs of genes indicate various behaviors. Figure S32 shows correlations of cell pair divergences (*R*) and whole-genome-normalized cell pair divergences (*R̃*) for three pairs of long genes chosen for illustration. All three examples have relatively high divergence correlations (*R* ∼ 0.5) but show qualitatively different behaviors for *R̃*. In the first case, the high divergence correlation can be explained simply by the large genome-wide divergence between cells so the normalized divergence correlation is relatively low (*R̃* ≈ 0.1). A similar conclusion holds for the second case. In the last case, the normalization by the whole-genome divergence leads to a decrease in the correlation, but the normalized correlation still remains high (*R̃* ≈ 0.7), suggesting these genes are tightly linked.

**Figure S32:**
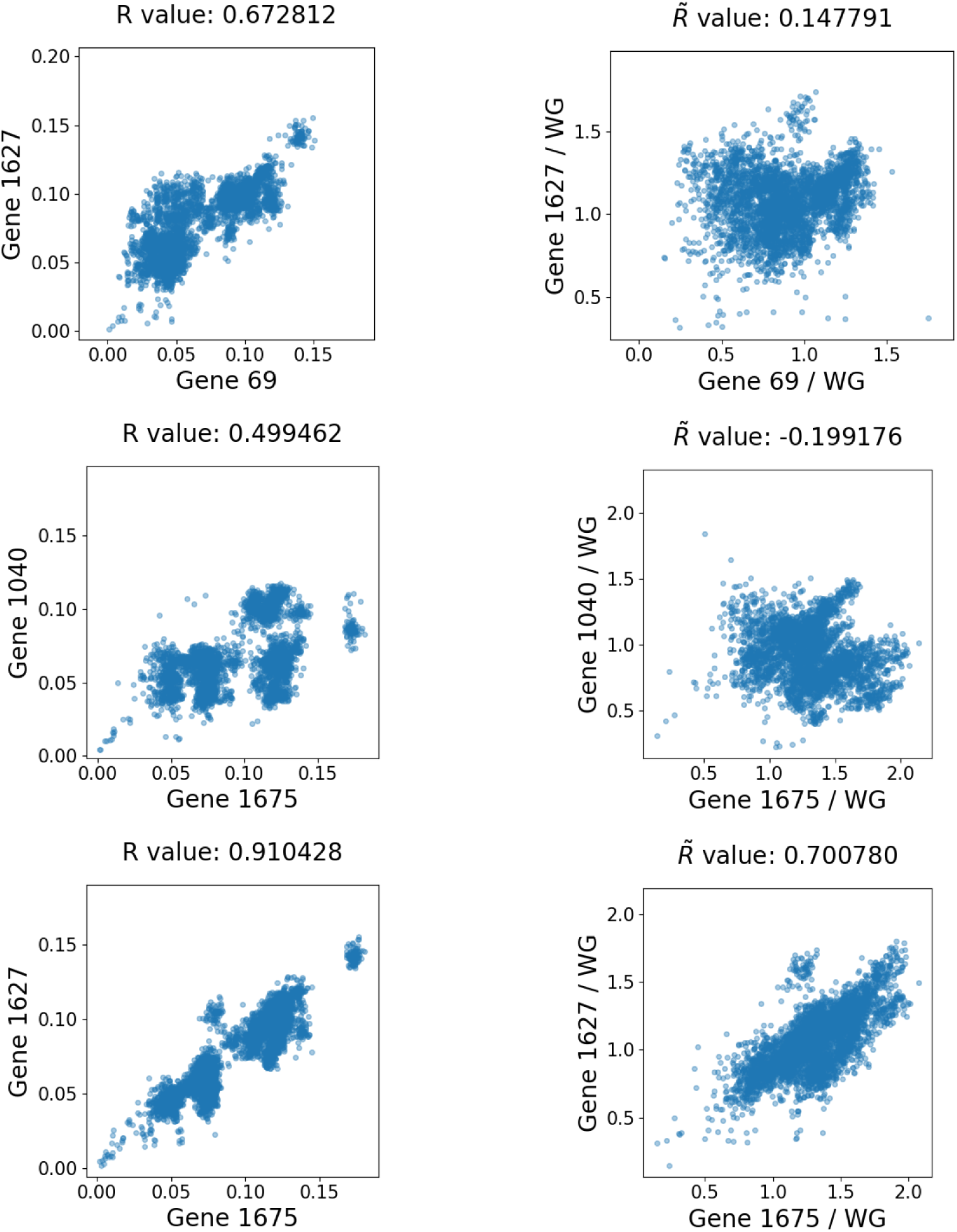
The correlations *R* and *R̃* for select pairs of the four genes whose PhyML-constructed trees are visualized in Figures S50, S51, S52, and S53. The top two pairs of genes have substantial correlation *R* (left column), but the scatter plot of normalized divergences *R̃* (right column), with indicate very weak residual correlation, showing that almost all the correlations between these genes is accounted for by the whole-genome divergences. In contrast, in the bottom row much of the correlation is not accounted for by the whole-genome divergences.

When averaged over longer genomic distances, both *R* and *R̃* vary over a wide range across different genome locations (Figure S33). Both correlations show distinct peaks and troughs over distances of ∼ 10 − 50 genes, suggesting significant local correlations. This is confirmed by the examining the variation of *R* and *R̃* with gene separation shown in Figure 3 (e) in the main text, which shows a notable decrease over a distances of ∼ 10 genes. We also observed that *R* has a residual positive value even on the longest genomic distances.

**Figure S33:**
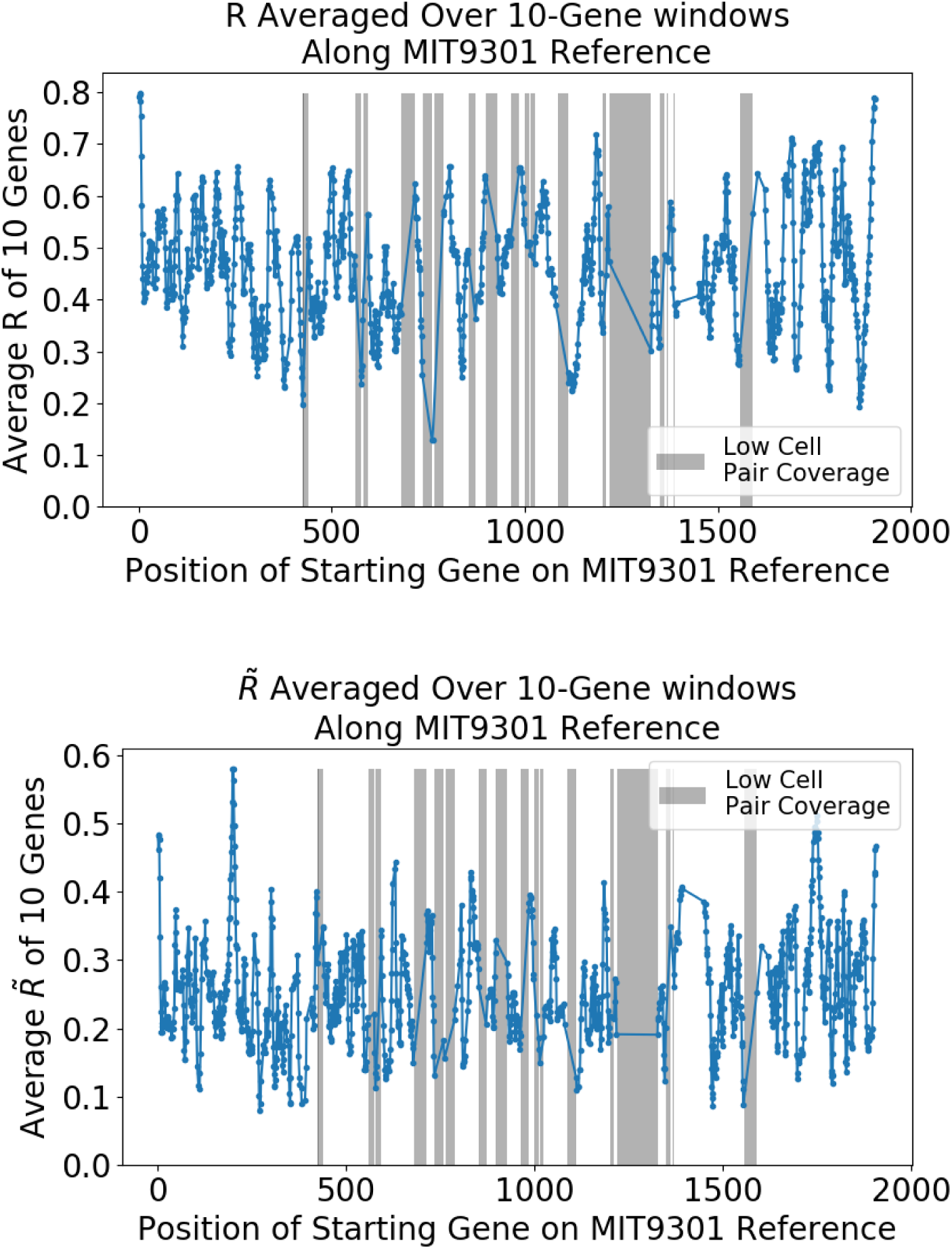
The gene pair divergence correlations along the MIT9301 reference genome. Each point represents an average over all pairs in a 10-gene sliding window of core genes. The two panels show the unnormalized correlation *R* (**top**) and normalized correlations *R̃* (**bottom**), calculated using twofold and fourfold degenerate synonymous sites for all covered cell pairs in the quasi-random dataset (see text for precise definitions of *R* and *R̃*).

To better understand the decrease and residuals of the correlations, we examined the distribution of *R* for different genomic distances. The distribution was unimodal across a range of gene separations from 1 to *>*100 genes and was broader than a fully unlinked null model (Figure S34). For the longest distances, normalizing the gene divergences by the genomewide divergences results in a shift in the distribution of *R̃* to mean close to zero, as expected (Figure S35). However, we did not find a noticeable decrease in the variance of *R̃*.

**Figure S34:**
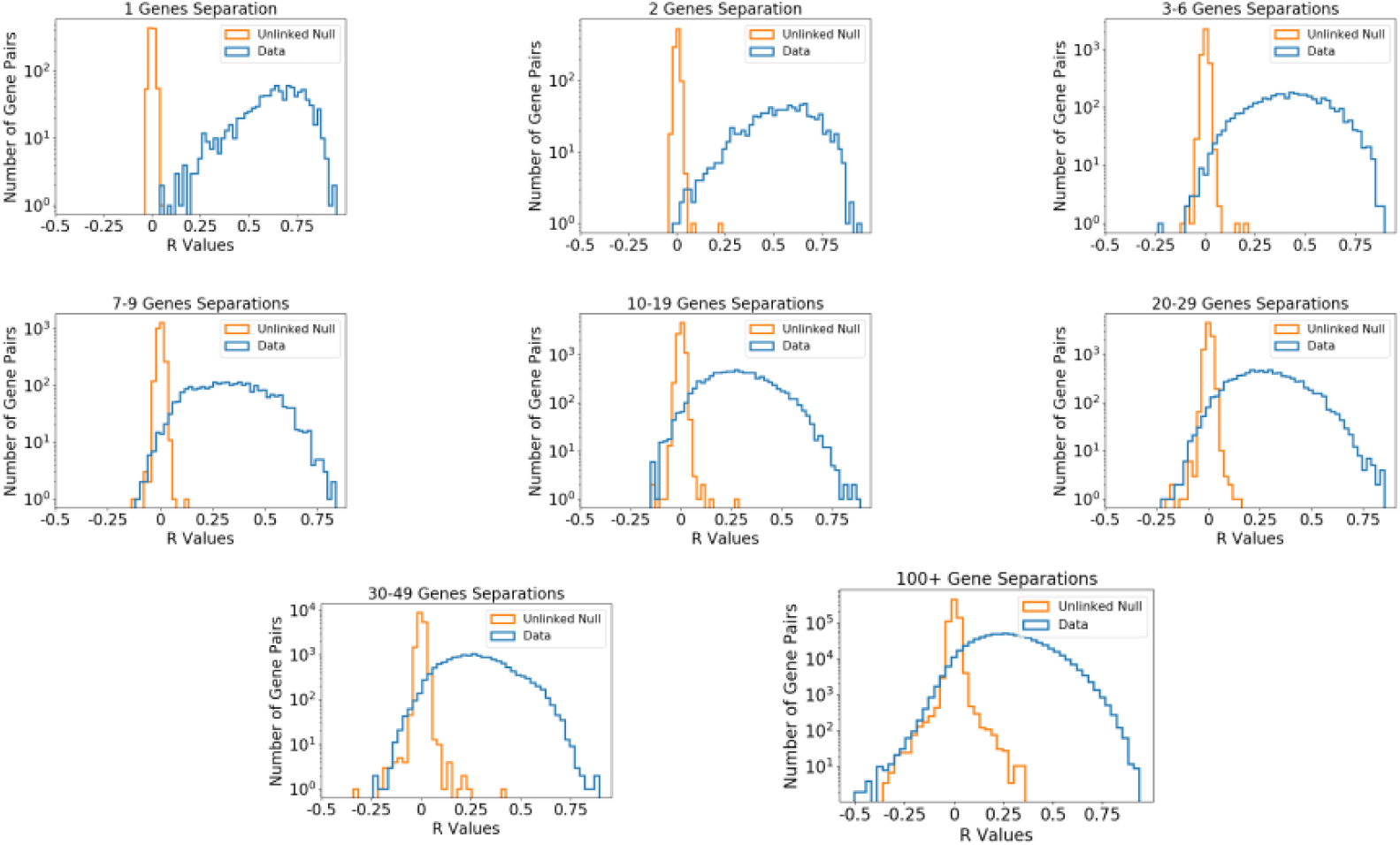
Histograms of gene pair correlations *R*, binned by separation between gene pairs for the quasi-random HLII dataset (using synonymous divergences). The linkage decays uniformly with separation and distributions are unimodal even for nearby gene pairs. This suggests that there are not many anomalously-highly-linked gene pairs. The uniform decay of *R* along the genome suggests that there have been recombination events at the boundaries of most gene pairs, rather than a few hotspots of homologous recombination along the genome (Figure S33). Note that the data in the last panel is the same as the left panel in Figure S35.

**Figure S35:**
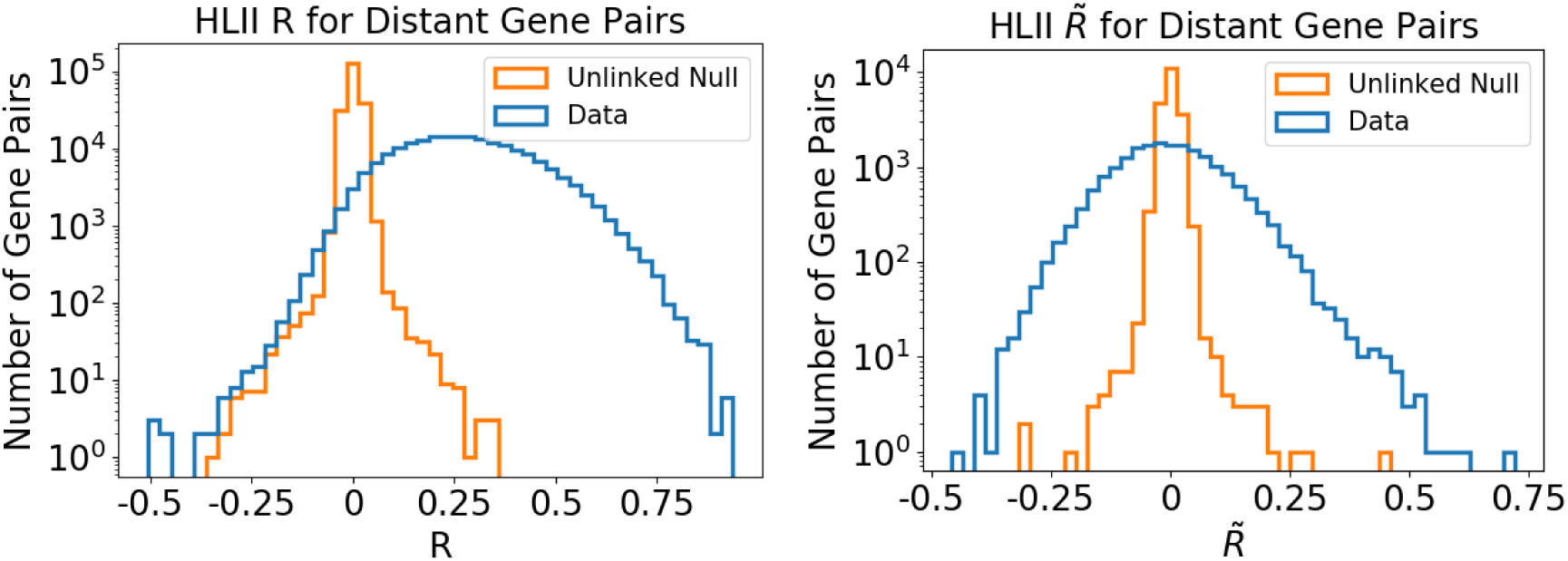
Histograms of *R* and *R̃* values for all core gene pairs separated by at least 100 genes along the reference genome, using twofold and fourfold degenerate synonymous sites, for the quasi-random sample of HLII. The data is compared to a random null in which cell pair divergences are shuffled at one gene, representing a completely sexual, unlinked population. The distributions are unimodal, consistent with residual linkage being due to whole genome weak epistasis rather than a few highly-linked outlier gene groups that would be expected if multiple discrete ecological niches were occupied by subgroups of HLII cells.

As a separate test of the effect of genome-wide divergences on the divergence correlation, we repeated the analysis from Figure 3 (e) in the main text on the quasi-equidistant dataset, for which genomewide divergences are sharply peaked around ∼ 6% (Section 1.). Contrary to the quasi-random cells, here we find that *R* decreases to a value close to zero (Figure S36). Below we consider two simple scenarios that could help explain the distributions of linkage correlations.

**Figure S36:**
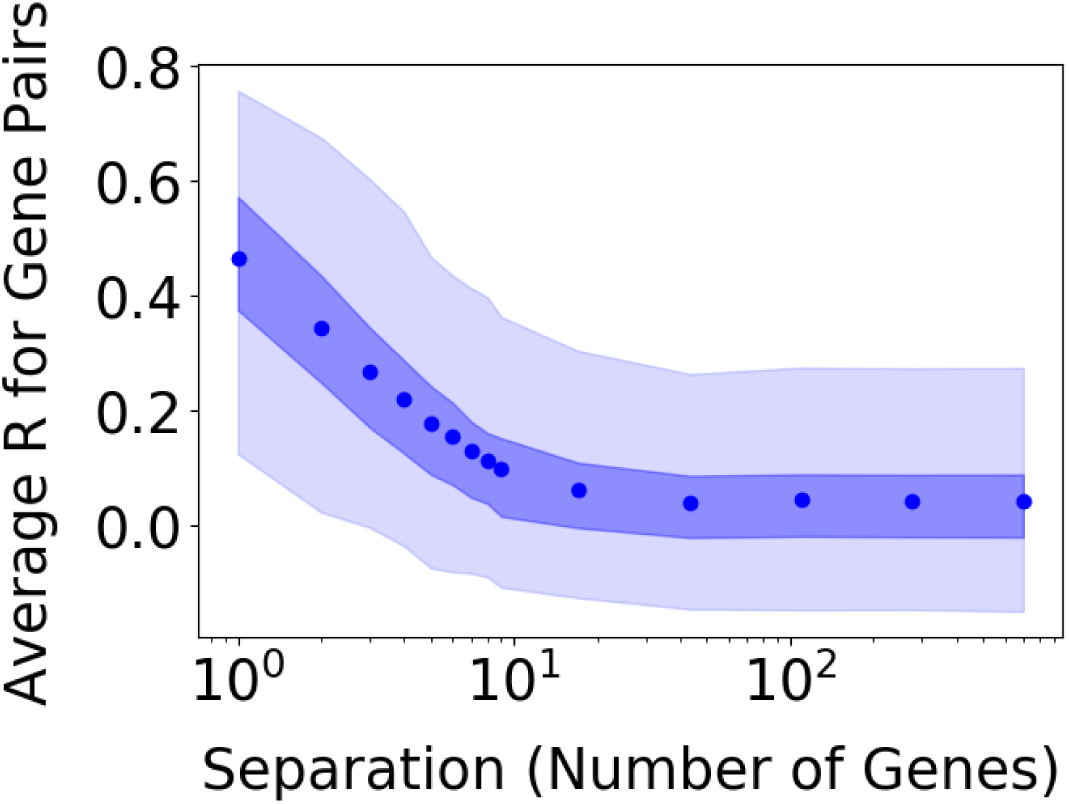
Correlation *R* of cell pair synonymous divergences at core genes as a function of gene separation along the genome in the quasi-equidistant subset of HLII (see Section A. 2.). Shaded bands show 1 and 2-sigma intervals for each separation bin. Note that the correlation decays by more than a factor of 10 over distances of 10 genes to *R* ≈ 0.04. The low value of *R* at large separations is consistent with a quasi-sexual population as explained in the text.

##### B.1. Divergence correlations under simple evolutionary scenarios

We consider two scenarios that could explain the results above. The first scenario is that the positive average *R* is the result of a small number of highly-linked genes scattered along the genome. This scenario could arise during the initial stages of evolution of new ecotypes, and has been observed previously in highly-recombining bacteria from the ocean (*22*). The second scenario is that the positive average *R* is due to weak correlations between the majority of genes. Such weak correlations could be the result of population structure or could be a residual signal of previous blooms, such as the inferred large ancient bloom. We simulate gene divergences in both models, calculate the correlations *R* and *R̃*, and compare their qualitative features to the data.

We first consider a two-ecotype scenario, though the results likely generalize to multiple ecotypes. In this model, there is some fraction of genes *f_genes_* that are clustered. These genes have 2 clusters of size *n*_1_ and *n*_2_ cells respectively. Clustered genes have average divergences *d_intra_*within each cluster and *d_inter_* between the two clusters. All clustered genes are perfectly linked. All other genes on the genome have cell pair divergences randomly distributed according to a Poisson with mean divergence *d_normal_* and gene length *l_gene_*.

We chose *l_gene_* = 170bp to approximately match the number of twofold and fourfold degenerate sites per HLII core gene, excluding six-fold degenerate codons. The remaining parameters are chosen to roughly match the mean and variance of the whole genome twofold and fourfold degenerate site divergence distribution within the quasi-random HLII dataset. Specifically, we use *d_intra_* = 0.01, *d_inter_* = 0.30, *d_normal_* = 0.14, *f_genes_* = 0.2, *n*_1_ = 34, *n*_2_ = 136, and simulate 100 genes. The results are shown in Figure S37 (left). The distribution of *R* across gene pairs is bimodal, depending on whether or not both of the genes in the pair have linked clusters. The distribution of *R̃* is also bimodal and has a larger average value than *R*. Note the high reproducibility of the mean correlation coefficient in both the normalized and non-normalized measures indicate that 100 genes is sufficient to minimize noise.

**Figure S37:**
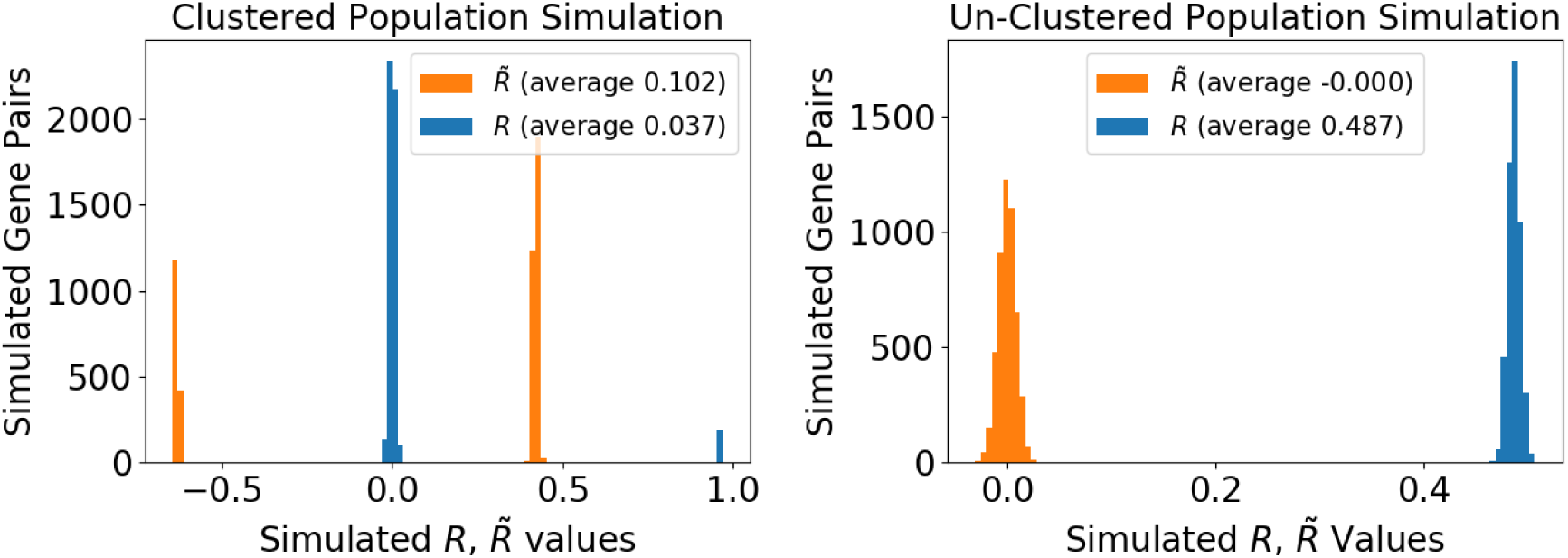
Simulations of gene pair divergence correlation coefficients in two evolutionary scenarios described in the text. **(Left)** An ecotype model in which distinct sub-clusters exist within HLII at certain highly-linked genes which determine ecological niches. **(Right)** An unclustered scenario in which all residual linkage is due to weak correlations between gene pair divergences along the entire genome.

We next consider a scenario in which there are weak correlations between cell pairs along the entire genome. To directly compare these results to the toy ecotype model above, we use the same whole genome divergence distribution generated from the first model. We then simulate 100 genes by drawing gene-level cell pair divergences from a Poisson distribution with mean equal to the gene length multiplied by the whole genome divergence of each pair. The results are shown in Figure S37 (right). The average *R* value is relatively high due to structure at the whole genome level. *R̃* is on average several orders of magnitude lower than *R*, and both distributions are unimodal.

Comparing the results in the two scenarios with the data, we see that the second qualitatively reproduces the unimodal distributions of the correlations (Figures S34, S35) and the positive residual at long genomic distances shown in Figure 3 (e) in the main text. These results show that the groups of highly-linked core genes found in Section 8. cannot account for the positive *R* at long genomic distances in the data. If this were the case, we would expect clear outliers in the distribution of *R*, which we do not observe in the data (Figure S35). Nevertheless, there are some features of the data that are not captured by our crude model. First, the distributions of both *R* and *R̃* appear much broader compared to both the model and unlinked controls, even at large distances. The broader distribution may be due to differences in the population structure of HLII from the one assumed in our model or may be a sign of more complex evolutionary dynamics (see Section 4.). Second, our model does not include recombination directly and therefore cannot capture the decrease in correlations with genomic distance found in Figure S36 and Figure 3 in the main text. We leave both of these as interesting topics for future work.

### 7. Flexible Gene Divergences

In this section we examine cell pair divergences at flexible orthologous gene groups which are found in at least 4 SAGs and at most 14 SAGs (for the deep clustered dataset) or 34 SAGs (for the quasi-random dataset). This upper cutoff is imposed to remove genes that are part of the core genome or are present in most genomes but have low coverage. We verified that our results are not sensitive to the choice of cutoff.

Within the quasi-random dataset, typical flexible genes have nucleotide divergence diameters similar to the HLII whole genome. For these genes, there are multiple evolutionary scenarios that are difficult to distinguish. For example, the genes may have been introduced into HLII from other clades once or multiple times and then subsequently swept through HLII, or they might have been core genes that have been partially lost in some genomes. We therefore focused our analysis on flexible genes in which the diversity is atypical relative to the deep clustered or quasi-random core genome. We find a few percent of flexible gene groups have diameters at least twice as high as the whole genome diameter and a similar percentage with very low divergences across all or most cells. The latter are consistent with recent gene sweeps.

We perform a similar analysis for the C1 SAGs from the deep clustered dataset. Within C1 the clonal backbone diversity is too low to identify sweeps of individual genes from the reduction in diversity. But we again find a few percent of flexible gene groups have diameters at least twice as high as the C1 whole genome divergence diameter. We find similar results when using only synonymous sites to calculate the diameter, suggesting these genes were likely transferred more than once independently from cells not in C1.

#### A. High-divergence flexible genes

We performed several checks to verify that the high-divergence flexible genes we observe are not due to errors in the construction of orthologous gene groups. First, we verified that each gene copy within the same group had the same annotation. In all cases, we found that the annotations were identical to each other. Second, we checked that each SAG contained a single copy of each orthologous gene group. We found only a small number of groups in which some SAGs had multiple copies of the gene. The few multiple copies we observed could be due to sequences being split into multiple reading frames during the processing of the raw assemblies.

Within the quasi-random dataset, some flexible genes have diameters greater than or equal to that between HLI and HLII, suggesting that transfers across clades, while rare, may still occur. This is consistent with our core-genome analysis, which revealed transfers from HLI into HLII in 10 core genes. In other cases, we find that flexible genes only rarely form distinct clusters. When we do observe clusters, intra-cluster divergences are less than the typical HLII core-genome divergences, but inter-cluster divergences are up to several times higher than typical. This suggests most of these flexible genes are the result of separate transfer events from outside HLII.

Within C1, high-divergence flexible genes have diameters comparable to the HLII core genome, several times larger than the C1 diameter. Such high diameter genes always have two or more clusters of cells each, with intra-cluster divergence consistent with typical C1 divergences. This suggests a scenario in which either a single allele from a non-C1 HLII cell was transferred into C1 and then spread as a partial gene sweep, or two alleles from a closely-related group of cells were separately transferred into C1.

#### B. Low-divergence flexible genes

Low-divergence flexible genes within the quasi-random dataset can be divided into two categories, depending on the number of distinct clusters within the othologous gene group. 15 of the 730 flexible genes in HLII we analyzed contained a single cluster with diameter less than 1% divergence, covered by on average 7 SAGs (range: 3-32 SAGs). Three gene groups have diamater less than or equal to typical C1 divergences (covered by 3, 3, and 10 SAGs respectively). This pattern is consistent with these genes spreading through some part of HLII by a relatively fast gene sweep. In other cases we find two of more tight clusters. In some cases clusters contain SAGs from both the Atlantic or Pacific Oceans, but not always. We also find some clusters with SAGs exclusively from the Atlantic (see also Section C. C.2. for discussion of flexible genes that are specific to the Atlantic and Pacific). This suggests that flexible genes are transferred over a range of time scales, with some spreading across different oceans and others specific to the Atlantic.

### 8. Searching for Ecotypes: Linkage and Ocean Correlations of Core and Flexible Genes

In this section we investigate genetic signatures that could indicate the presence of distinct ecotypes within HLII. We used the linkage metrics introduced in Section 6. to identify groups of highly linked core genes and analyze the presence or absence of flexible genes.

#### A. Functional annotations of highly linked genes

We identified groups of core genes with high values of *R̃* between them – here defined as *R̃* ≥ 0.6, comparable to typical pairs of genes 2-genes apart (see Section 6. for details). Figure S34 Most of these groups of genes lack a clustered structure that would indicate well-separated, distinct alleles. And for most groups, the gene annotations did not suggest any clear functional pattern. Nevertheless, the enhanced correlations suggest there may be epistatic interactions among them.

Among the groups of highly linked core genes, one contains rpoC1, which has been used as a proxy for whole-genome genome divergence (*23*, *24*). It is noteworthy that rpoC1 does tend to have higher correlation with whole genome divergence than ITS (Figure S39) and appears to be linked to some other core genes. The other genes in this group include phosphate and phosphatase-related genes (Protein phosphatase 2C domain-containing protein, carbamoyl-phosphate synthase small chain and large subunit, phosphoribosylglycinamide synthetase); nitrogen-related genes (dCTP Deaminase, Global nitrogen regulatory protein); GroEL protein; and a few tens of others. Thus, rpoC1 may be a useful proxy for alleles at these linked genes, but not for the majority of core genes along the HLII genome.

**Figure S38:**
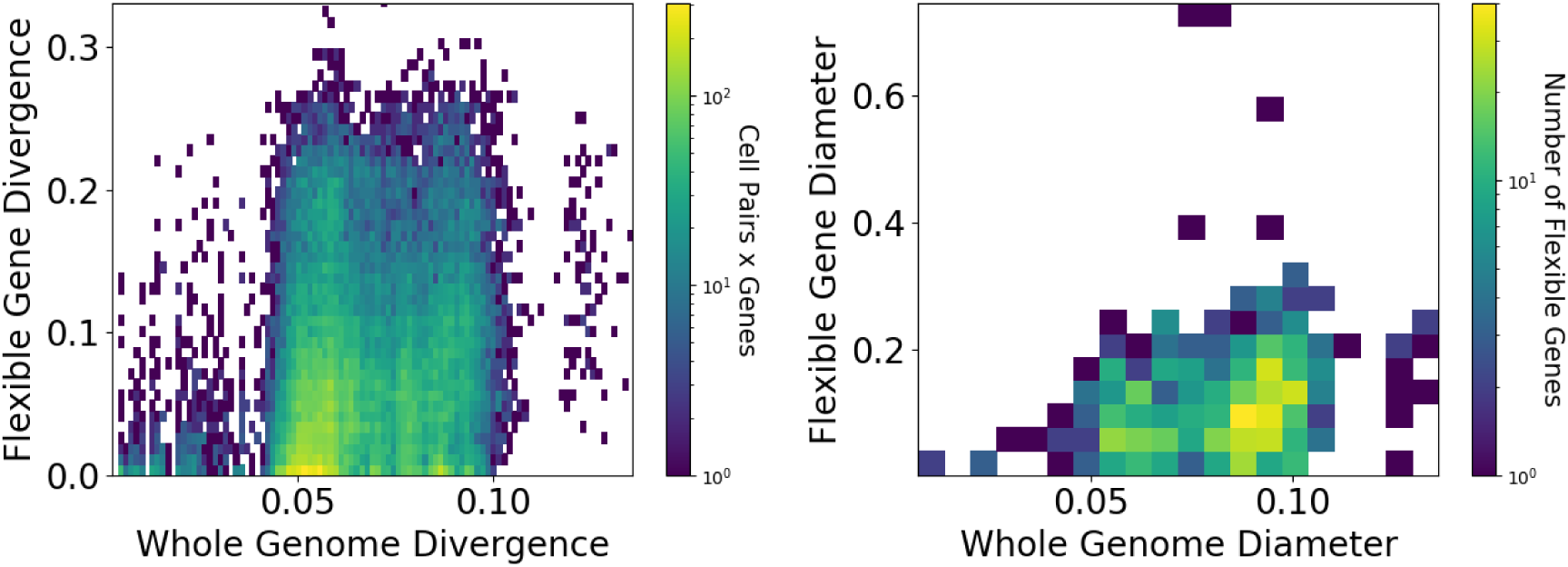
Within HLII, flexible genes come in multiple flavors. Most flexible genes have divergences similar to typical HLII core gene divergences, but others show evidence of recent partial sweeps (more recent than the advent of closely-related cell group C1, for example), and yet others may have been transferred into HLII from cells outside the clade (such as from other *Prochlorococcus* clades like HLI, or beyond the species), possibly multiple times. Here, flexible gene orthogroups with a gene found in 4-34 of the SAGs of the HLII quasi-random sample are plotted, using nucleotide divergence (730 such groups). On the left, each point represents a single cell pair at a flexible gene. The pair’s divergence at that gene is plotted on the y axis and its mean core genome divergence is plotted on the x axis. An enrichment of points at low flexible gene divergence is seen. This is sometimes due to short genes which are identical by change, but other times due to recent flexible gene sweeps. There are also many pairs with flexible gene divergence much higher than their whole genome divergence. A small number of outlier points at even higher flexible gene divergence are excluded. On the right, each point represents one flexible gene orthogroup. On the y axis, the nucleotide divergence diameter of the gene group is plotted, and on the x axis the whole genome divergence diameter of the SAGs with an ORF in the corresponding orthogroup is plotted. Typical flexible genes have diameters consistent with the cells’ whole genome diameter, but there are exceptions.

**Figure S39:**
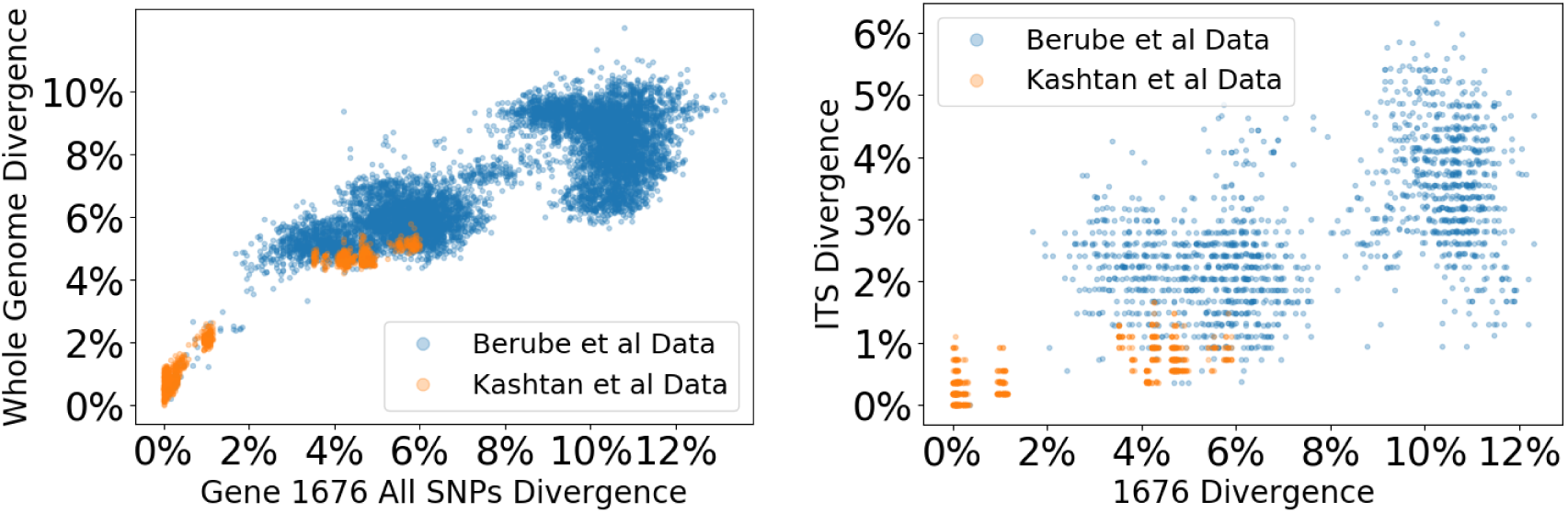
The gene rpoC1 correlates reasonably well with core genome divergence when closely-related SAGs are included (orange points), though this correlation decreases if only quasi-randomly sampled HLII SAGs are considered (blue points). (a) Correlation between the gene rpoC1 (which is used as a whole genome proxy in (*23*)) and the mean core genome nucleotide divergence for SAG pairs within the quasi-random HLII dataset and the deep sampling dataset consisting of 5 closely-related clusters. Including all SAGs gives a correlation coefficient *R* ≈ 0.88. When only the quasi-random sample of HLII cells, representative of typical HLII divergences, is considered, *R* ≈ 0.8. Note, however, that the relationship for typical HLII cells is sub-linear. (b) Correlation between rpoC1 and the ITS gives *R* ≈ 0.77 when all SAGs are considered, but *R* ≈ 0.57 when only the quasi-random HLII cells are included. In Figure 2 of the main text, for comparison. the correlation between ITS and whole genome divergence is 0.8 when all SAGs are used, and 0.56 when only the quasi-random sample of SAGs is included.

#### B. Linkage of presence/absence of flexible genes

Quantifying the correlations in the presence/absence of flexible genes is challenging because the true absence of a flexible gene from a SAG is difficult to distinguish from lack of coverage at that locus. To address this issue, we calculate the ⟨*r*^2^⟩ linkage coefficient between pairs of flexible genes using the *observed* flexible gene frequencies and compare the results to the expectation ⟨(*r_*ab*_^max^*)^2^⟩ for a fully linked pair of genes with *true* frequencies *f_a_* = *f_b_* = *f_ab_* ≡ *f*. Note that because SAG coverage is inherently random, the *observed* frequencies of the two genes *f_a,o_*, *f_b,o_*, and the genotype *f_ab,o_* will generally be different from the true frequency *f*and will vary between pairs of genes even when the true frequency is the same. Thus, while it is possible to observe pairs of fully linked flexible genes, the expectation of the linkage coefficient ⟨(*r_*ab*_^max^*)^2^⟩ will be less than one and will depend on the probability that a gene is covered, *c*, and the observed frequency of each gene. To estimate ⟨(*r_*ab*_^max^*)^2^⟩ we assume that each gene is independently covered with a probability *c* ≈ 0.7. The assumption that SAG coverage is independent across genes quantitatively reproduced correlations in presence/absence of core genes in (*17*) and should be a good approximation. Using this, we can calculate the expected ⟨(*r_*ab*_^max^*)^2^⟩ in terms of the observed gene frequencies as

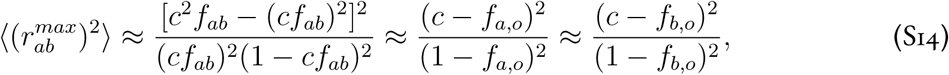

*max* 2

where we used *f_ab,o_* ≈ *c*^2^*f_ab_* ≈ *cf_a,o_* ≈ *cf_b,o_*. Note that for low-frequency fully-linked gene pairs, *f_ab_* ≪ 1, the above expression is approximately the square of the coverage probability, and hence ⟨(*r_*ab*_^max^*)^2^⟩ ≈ *c*^2^ simply from the coverage being roughly independent at each site. But at high pair frequencies, it is very likely that some genotypes lacking both genes will be produced and ⟨(*r_*ab*_^max^*)^2^⟩ becomes small. However for the criteria we have used to define flexible genes, most of the flexible gene frequencies are low enough that the reduction factor from *c*^2^ is roughly independent of the observed frequencies (Figure S41).

**Figure S40:**
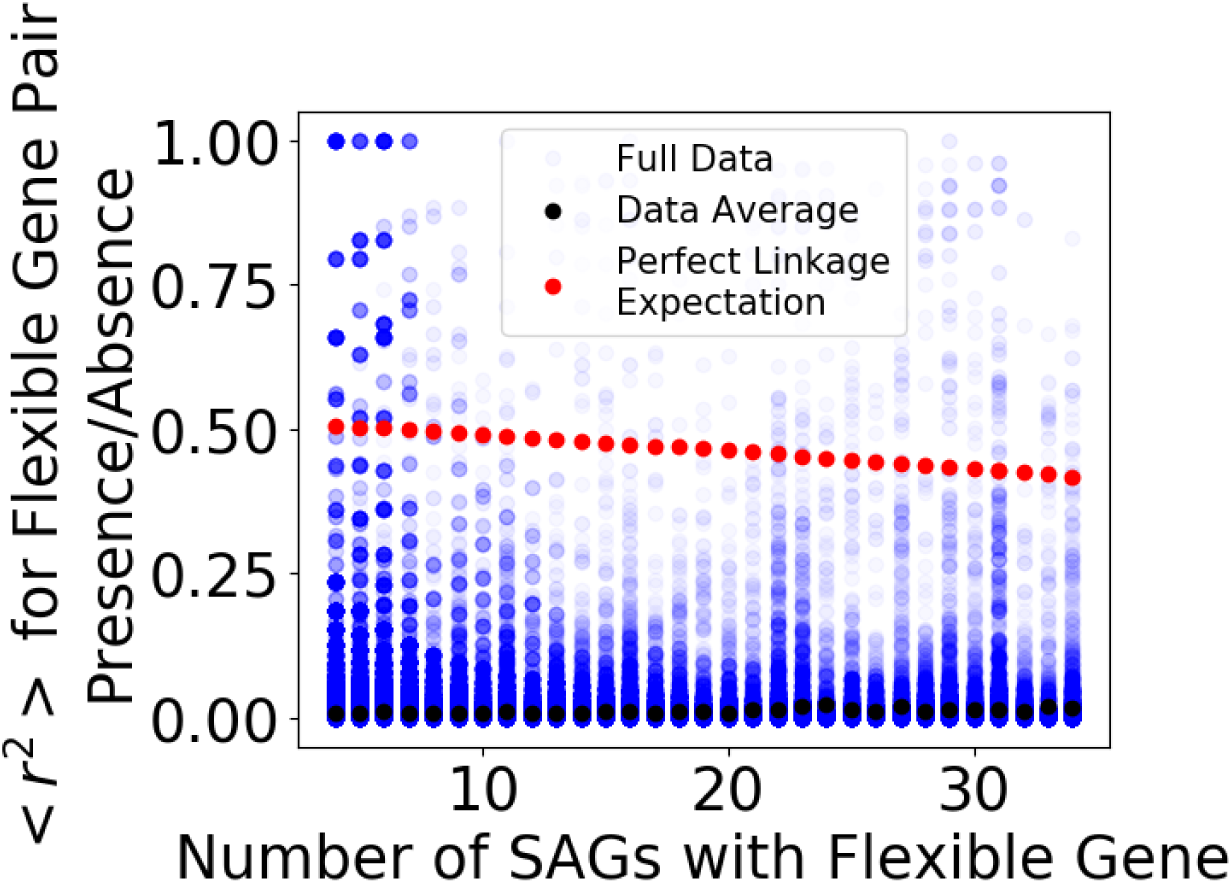
Comparison of flexible-gene pair linkage coefficients ⟨*r*^2^⟩ to expectation, ⟨(*r_*ab*_^max^*)^2^⟩, if pairs were fully linked, shown as a function of the individual gene frequencies in each pair. On the x axis is the number of SAGs in the quasi-random dataset (of 167 total) which have a given flexible gene covered, such that the observed gene frequency, *f_a,o_*, is the x axis coordinate divided by 167. For the data point s shown, each pair of flexible gene is represented as two points with the same observed *r*^2^ value, but with the two x-axis values equal to the number of SAGs observed with each of the two genes in the pair. The perfect linkage expectation was calculated from Eq. S14 with *c* = 0.7, for all possible observed gene frequencies we consider (4*/*167 ≤ *f_a,o_, f_b,o_* ≤ 34*/*167). Note that some data points reach or exceed the value expected from the rough model for perfect linkage.

**Figure S41:**
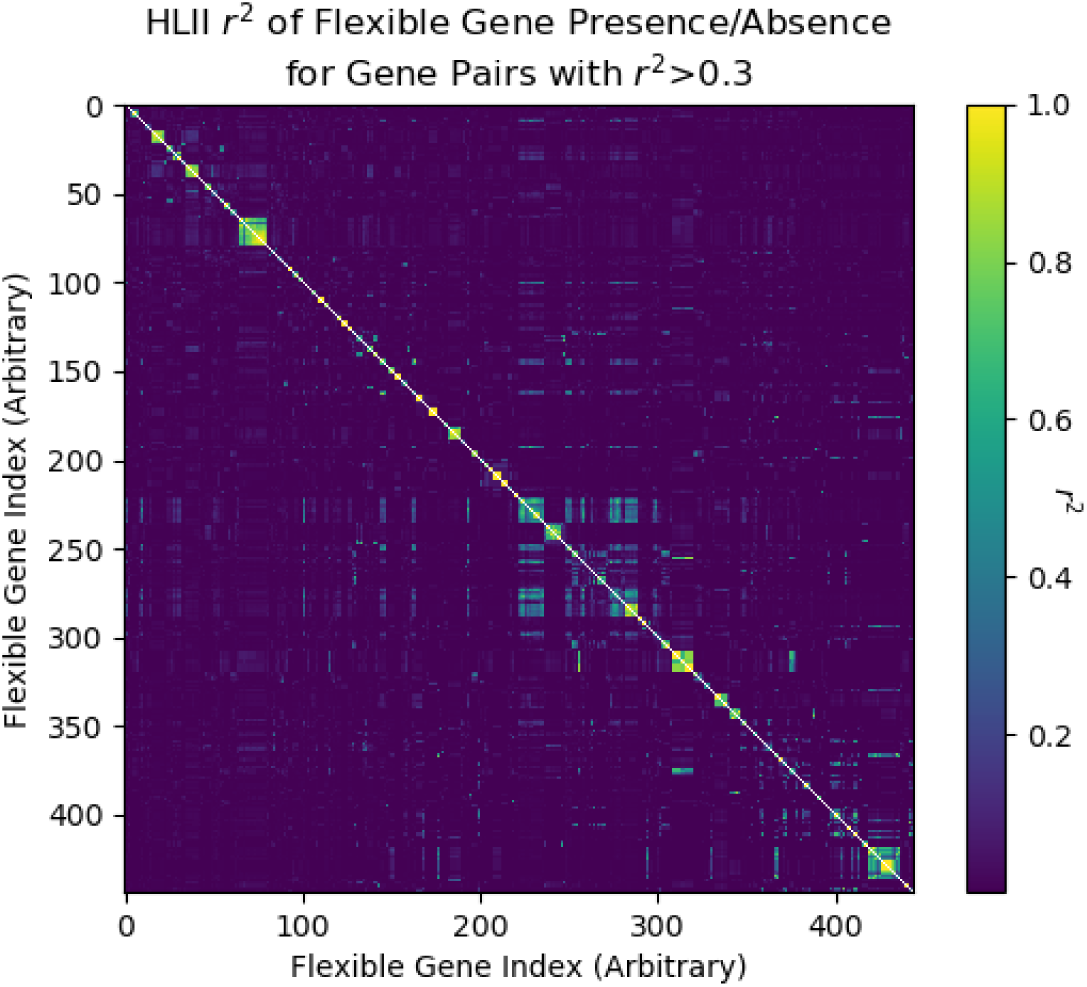
A small number of flexible genes have strong presence/absence linkage in quasi-random HLII. Matrix shows the *r*^2^ value (Equation S10) for different pairs of flexible gene orthogroups. Only orthogroups present in 4 - 34 SAGs in the quasi-random HLII sample and having *r*^2^ ≥ 0.3 with at least one other orthogroup were considered. Note the small clusters of strongly-linked genes. Compare to expectation for ⟨*r*^2^⟩ from perfectly-linked genes shown in Figure S40.

We searched for tightly-linked pairs of flexible genes in the 421 groups of flexible genes with intermediate frequencies range of 4-34 SAGs within the quasi-random HLII sample (Section B. B.3.). We compared the expectation from (S14) with the *r*^2^ calculated from the data and found that most gene pairs had *r*^2^ much less than ⟨(*r^max^*)^2^⟩ (Figure S41). In addition, we also observed a small number of pairs with *r*^2^ comparable to or greater than ⟨(*r^max^*)^2^⟩. Some of the higher values are likely due to statistical fluctuations not accounted for in our estimate, but others may be due to correlations in the coverage probability of nearby genes. However, because flexible genes do not align to the HLII references, the ordering of these genes along the genome is not easily known and we did not attempt to distinguish between these hypotheses.

Figure S41 shows the matrix of *r*^2^ values for all pairs of flexible genes with *r*^2^ *>* 0.3 with at least one other gene. The cutoff for *r*^2^ was chosen to reduce noise but still be somewhat smaller than *c*^2^ to retain intermediate frequency gene pairs. The results show a few blocks of highly-linked gene groups, which may be co-located on the genome. We find that blocks are due to positive correlations in the presence of a group of genes and did not find genes that were linked to the absence of other genes. 75 pairs of genes with *r*^2^ = 1 exist (0.01% of examined flexible gene pairs; and which comprise 1 gene pair found in 11 cells, 12 gene pairs found in 7 cells, 17 gene pairs found in 5 cells, and 45 gene pairs found in 4 cells), most of which are unannotated, though some include nitrate/nitrite transport system genes (a groups of three genes); iron III transport system gene pair; phosphonate transport binding gene pair cyanate lyase gene pair; and a transcriptional regulator protein linked with a major facilitator protein. Groups of cells that have highly-linked flexible genes tend to span both oceans rather than being ocean-specific.

#### C. Ocean-specific genes

##### C.1. Core genes with ocean-specific alleles

Core gene alleles are typically well-mixed between oceans within the quasi-random sample of HLII SAGs, which includes SAGs from a variety of locations and depths. Figure S42 shows the distributions of cell pair divergences at core genes, binning cell pairs as Atlantic-Atlantic, Pacific-Pacific, and Atlantic-Pacific according to the sampling location of each cell in the pair. The distributions are highly similar, with only a slight enrichment for closely-related alleles among pairs with both cells from the Atlantic. When we restrict to only the BATS site from the Atlantic and the HOT sampling site from the Pacific, this enrichment is more evident, but remains modest. The majority of alleles are well-mixed between oceans.

**Figure S42:**
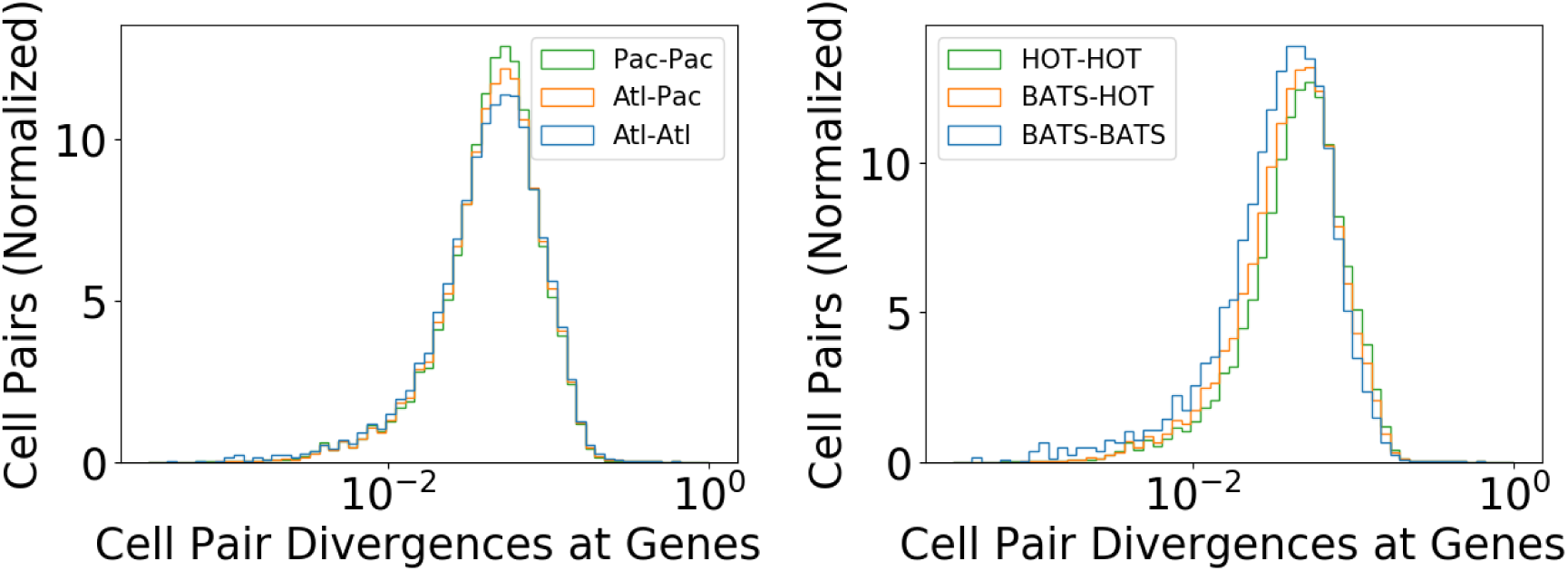
The distribution of cell pair divergences at HLII core genes reflects well-mixed oceans. We first remove closely-related cell pairs (clusters of cells with less than 4% whole genome divergence, which are strongly correlated with the ocean (Section 9.) and then divide cell pairs into three groups. **(Left)** Distributions for pairs in which both cells are from the Atlantic (blue), both cells are from the Pacific (green), and in which one cell is from the Atlantic and the other from the Pacific (orange) are shown. **(Right)** Instead of using all quasi-random SAGs from the Atlantic and Pacific oceans, we confine to just the BATS sampling site for the Atlantic and just the HOT sampling site for the Pacific. The distributions are highly similar, with some enrichment for closely-related alleles among Atlantic pairs.

For a more detailed comparison, we analyzed the divergences for each core gene separately (Figure S43). For most genes, typical cell pairs from the same ocean are just as diverged as pairs from different oceans, as shown by the high density of points along the diagonal in Figure S43. We also find a small number of genes with tight clusters in one ocean. We define such tightly-clustered outlier genes as genes at which the intra-ocean median divergence is less than 0.66 times the inter-ocean divergence. We sometimes refer to these as *ocean-specific genes*, although they are only partially specific.

**Figure S43:**
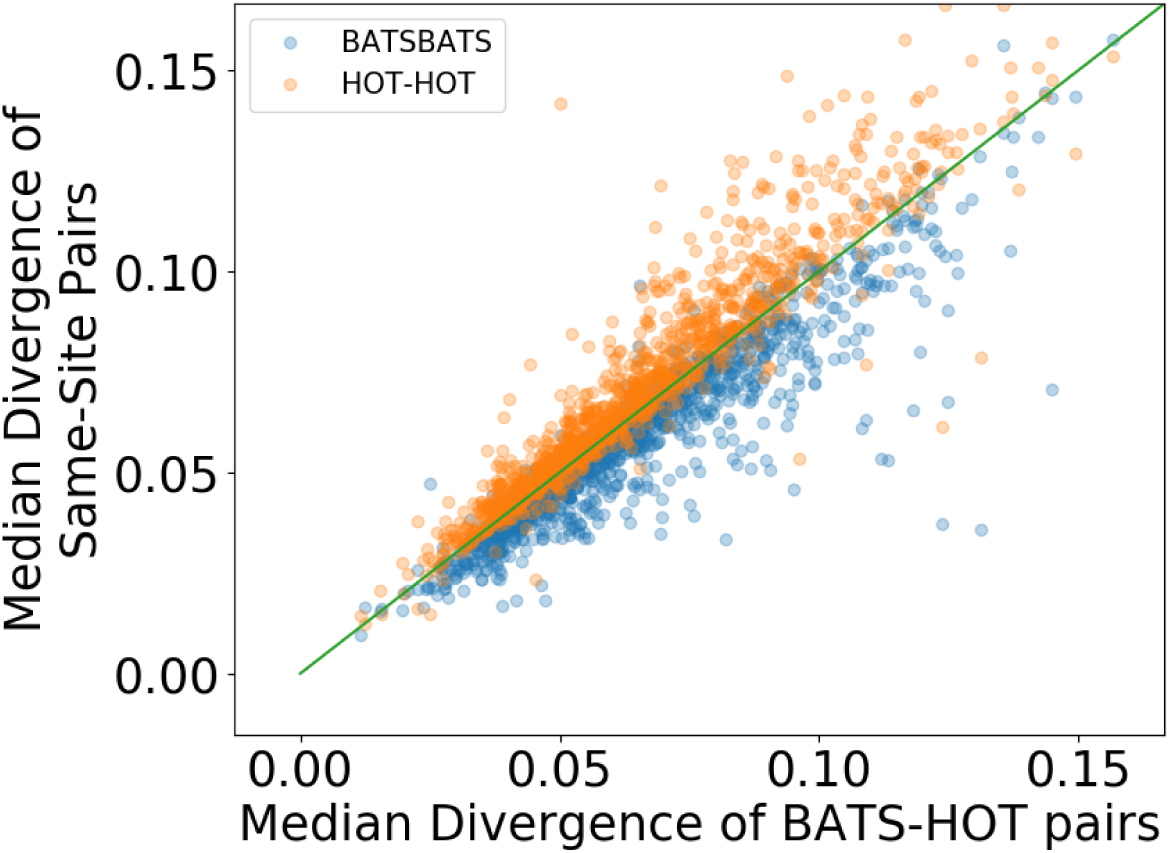
At most core genes, typical cell pairs divergences from the same ocean are similar to those from different oceans. Each gene in the HLII core genome (Section B. B.2.) is represented as one blue point and one orange point. On the x-axis is the median nucleotide divergence across all pairs of cells from the quasi-random sample in which one cell is from BATS (Atlantic ocean) and the other from HOT (Pacific ocean), at a given gene. On the y-axis is the median divergence at that gene, taken over cell pairs with both cells drawn from the BATS sampling site (blue) or both from the HOT sampling site (orange). The line *y* = *x* is shown in green for reference. Note that most genes cluster near the *y* = *x*, consistent with being well-mixed between BATS and HOT, with a small number of outliers below the diagonal.

We find about 40 core genes that are relatively tightly-clustered among BATS sample site cell pairs, and 7 genes that are relatively tightly-clustered among HOT sample site cell pairs. Many of the core genes that have ocean-specific alleles are related to phosphate utilization, including one pair of phosphate ABC transporter genes that form tight clusters in both BATS and HOT. In total, of the 47 core genes in the MIT9301 reference genome that contain “phosphate” in their annotation, 7 have ocean-associated clusters, including all three phosphate ABC transporters. In addition, two of the 15 genes with annotations containing “phosphatase,” and three of the eight genes with annotations containing “iron” or “ferric” are ocean-associated genes. But all together, ocean-associated genes represent less than 4% of HLII core genes. Thus, phosphate, phosphatase, ferric, and iron-related genes are significantly enriched among ocean-associated core genes.

Of the 47 ocean-associated genes, ∼ 15 have two clear clusters based on hierarchical clustering of nucleotide divergences, using average-linkage clustering. None of these genes have clusters that perfectly separate Atlantic and Pacific SAGs, but five have clusters comprising SAGs from predominantly one ocean, with a small minority of ∼ 5 − 10 SAGs from the other ocean. Three of these five genes are adjacent on the reference genome and are annotated as putative phosphate ABC transporters. About 600 genes away on the reference genome is another phosphate ABC transporter at which cells form largely ocean-specific clusters. Between these two locations on the reference, the final ocean-specific core gene observed is annotated as a ferric uptake regulator.

##### C.2. Flexible genes with ocean-specific presence/absence

Flexible genes that are ocean-specific are identified among the 421 flexible genes with intermediate frequencies (Section 1. B.) using a simple cutoff. Specifically, genes where at least 80% of cells are from the Atlantic or the Pacific, are identified. In total, 114 flexible genes meet this threshold, including 40 which are present in SAGs sampled exclusively from a single ocean. A few gene annotations stand out as enriched among ocean-specific genes. Three iron (III) transport system genes are listed in the annotated flexible gene groups, all of which are found exclusively in SAGs from the Atlantic. The list also includes ten phosphonate-related genes, nine of which are found mostly in the Atlantic, including six which are found exclusively in the Atlantic. Annotations of flexible genes found mostly in Pacific SAGs are not as easily tied to known chemical differences between oceans. The enrichment of ocean-specific flexible genes as compared to a well-mixed null is shown in Figure S44 and Figure S45.

**Figure S44:**
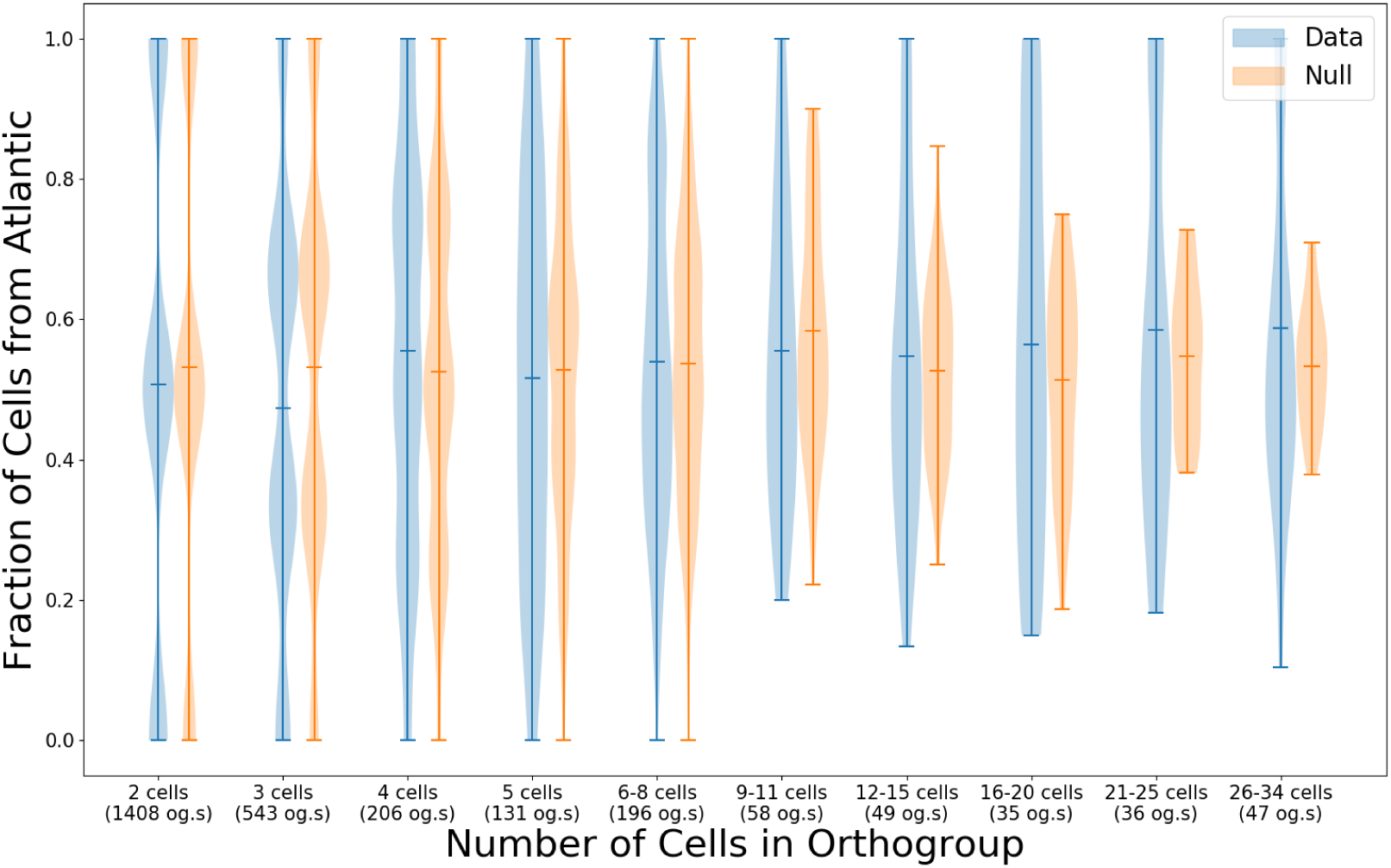
The distribution of flexible genes across the Atlantic and Pacific oceans is similar to expectation from a null model in which the two oceans are perfectly mixed. Flexible orthologous genes groups found in quasi-randomly sampled SAGs from the Atlantic or Pacific are binned according to the number of SAGs in which they are detected (shown on the x-axis). The fraction of cells from the Atlantic is shown as a blue violin. A null distribution is determined by drawing cells randomly from the Atlantic and Pacific, for each gene group in the bin: shown in orange. Only flexible gene orthologous gene groups that were found in ≤ 20% of quasi-randomly sampled cells were included. The 20% cutoff was chosen to minimize the number of poorly-covered core genes and is close to the trough in the distribution of orthologous gene groups abundances shown in Figure S8. Note the small but systematic discrepancy between the data and the null model for bins containing more than 12 SAGs.

**Figure S45:**
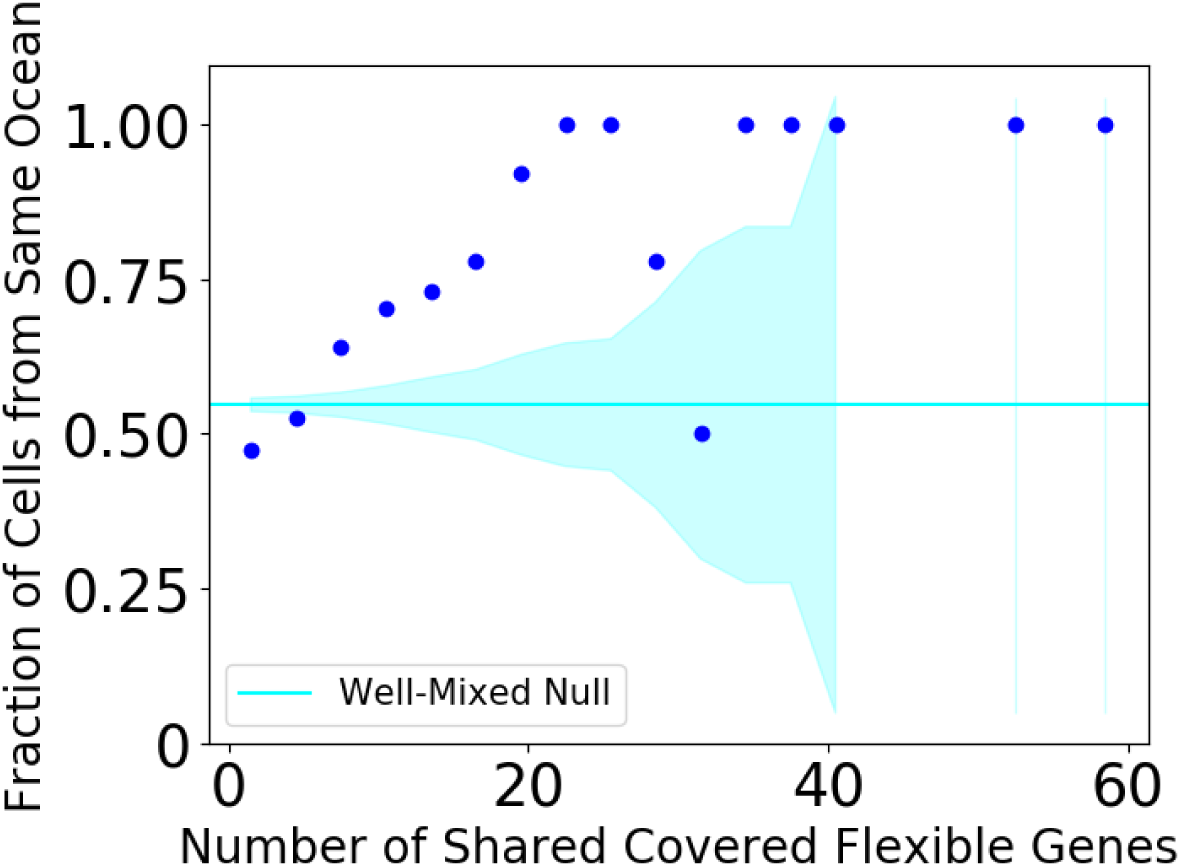
Cell pairs sharing relatively many flexible genes tend to be from the same ocean. The number of shared flexible genes for different quasi-random HLII cell pairs is calculated and binned values are shown on the x-axis. For each bin, the fraction of pairs from the same ocean is calculated and shown on y-axis. Close pairs with whole genome divergences ≤ 4% we excluded. The probability that a SAG pair was sampled from the same ocean increases with the number of flexible genes the pair has in common, reaching nearly 100% probability by the time SAGs share 20 covered flexible genes.

### 9. Ocean Mixing: Closest Genomes and Genes Across Oceans and Methods for Null Models

In this section we use nucleotide divergences to compare the HLII populations in the Atlantic and Pacific oceans.

#### A. Divergences across oceans

We analyzed correlations between whole-genome divergences at different locations to understand how much spatial structure contributes to the patterns we observed. Within the quasi-random dataset, divergences between SAGs from the same sample were not significantly different from those between samples. Similarly, whole genome divergences of inter-ocean cell pairs are similarly-distributed to those of cell pairs with both cells from the Pacific (Figure S46). The one notable exception was one sample from the BATS location, which contained several dozen closely-related cells. Figure S47 shows the mean core genome divergence of close pairs in the quasi-random dataset, binned by which ocean the cells in the pair are sampled from. Consistent with the findings above, the Atlantic ocean contains the closest pairs in the dataset, and inter-ocean close pairs are similarly diverged as pairs with both cells from the Pacific.

**Figure S46:**
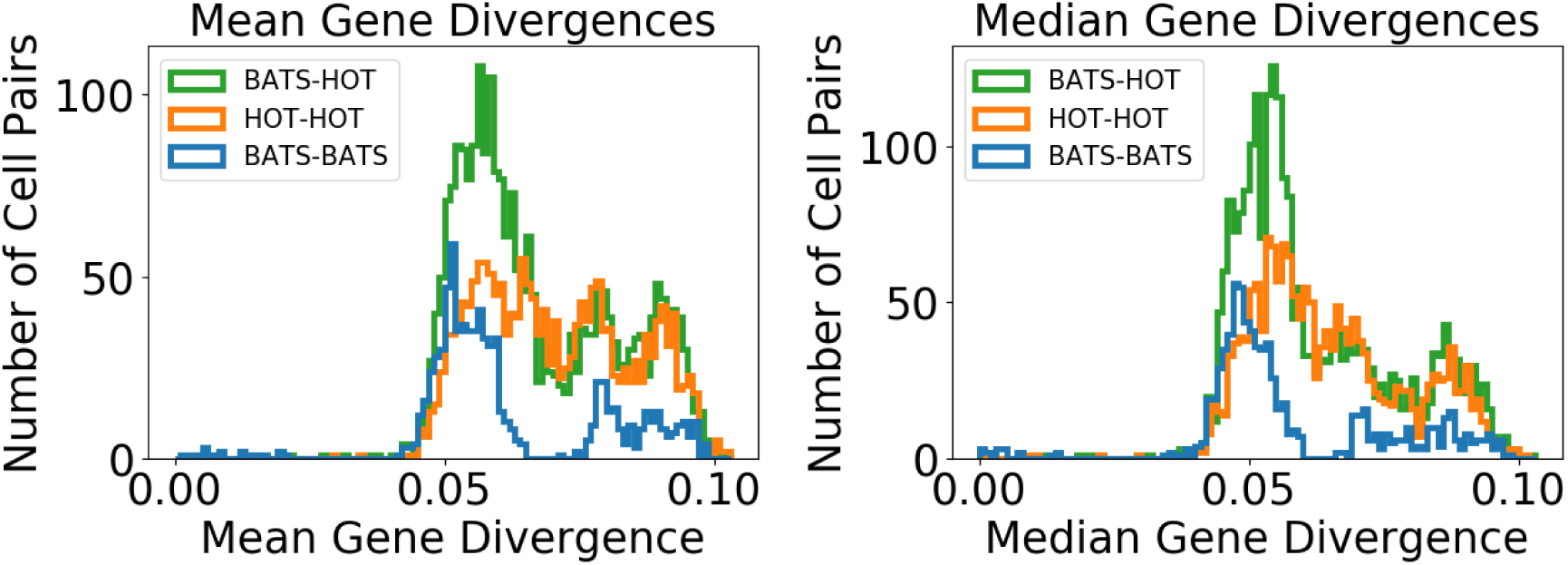
Whole genome divergence distributions of cell pairs sampled from BATS (Atlantic) show an enrichment of closely-related cell pairs compared to HOT (Pacific). Mean (**left**) and median (**right**) core-genome divergences between all HLII cell from either BATS or HOT from the quasi-random dataset are shown. Note the gap at just over 6% divergence in the BATS data. Cells just below this gap form the quasi-equidistant cell group which includes Pacific cells.

**Figure S47:**
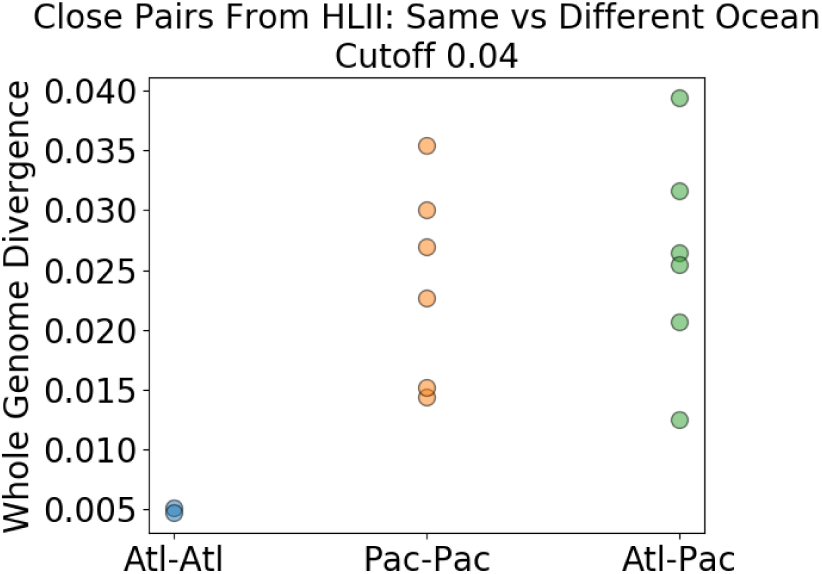
The closest pairs from the quasi-random dataset are found in the Atlantic. Using a 4% divergence cutoff, close pairs of cells are categorized by ocean and their mean core genome divergence plotted. Consistent with other results, we see that the closest cells here are from the Atlantic, and that divergences within the Pacific are similarly distributed to divergences across the Atlantic and Pacific.

We compared the divergences of close groups from the Atlantic to those containing SAGs from both the Atlantic and Pacific. The lowest divergence between SAGs from the Atlantic and Pacific depends on the number of mutually covered core genes for the pair. If we require that at least several hundred core genes be mutually covered, we find a cell pair that is part of a group of three closely-related SAGs, with 712 mutually covered core genes that has median core gene divergence of 1.3% and mean core gene divergence of 2.1%, using total nucleotide divergence. Another pair, with 1024 mutually covered core genes, has median core gene divergence of 2.1% and mean core gene divergence of 3.0%. This cell pair is part of the mixed-ocean 7-cell cluster described in Section B. B.4..

At the gene level, there is evidence of more rapid mixture between oceans. To probe the smallest divergences, we search for strings of consecutive identical and nearly-identical genes in inter-ocean cell pairs. We find two inter-ocean cell pairs which each have strings of three consecutive identical or 1-SNP genes, equating to nucleotide divergences of order 10^−3^ in both cases. Neither of these pairs are part of the 7-cell inter-ocean close cluster mentioned above, and they have median core gene divergences of order 5.5%. This suggests that the nearly-identical gene strings are the results of relatively recent transfer events. One of these gene trios has annotations of: phosphonate ABC transporter, phosphonate binding protein for ABC transporter, and aminotransferase (while the other gene trio consists of unannotated proteins). For comparison, the longest nearly-identical gene stretch among cell pairs with both cells from the Pacific ocean is 4 genes, while for cell pairs from the Atlantic ocean it is 50 genes. This again highlights the similarity in divergence of inter-ocean cell pairs compared to Pacific cell pairs, but the enrichment for close cells in the Atlantic.

If we include the close groups of SAGs identified above in the quasi-random dataset, we find 846 out of 1253 HLII core genes have at least 1 identical cell pair with both cells from the Atlantic, 169 genes have at least 1 identical inter-ocean cell pair, and 141 genes have at least 1 identical cell pair from the Pacific (using total nucleotide divergence, for covered genes only). If we exclude close cells, we find 102 genes have at least 1 identical cell pair with both cells from the Atlantic, 109 genes have at least 1 identical inter-ocean cell pair, and 98 genes have at least 1 identical cell pair from the Pacific. Note that more cells are excluded from the Atlantic than from the Pacific when close cells are excluded (Atlantic cells decrease from 82 to 52 while Pacific cells decrease from 83 to 65).

#### B. Null models for inter- and intra-ocean closest relatives in Figure 4 (main text)

##### B.1. Null model for the fraction of core genes per cell at which the closest relative is from a different ocean

To construct the null histograms from Figure 4 (a) in the main text, we used the following procedure. For each SAG at each gene we randomly sampled an allele from Atlantic and Pacific SAGs only and assigned it as the closest relative. We then calculate the fraction of genes among all covered core genes at which the closest relative assigned using this null model is from the same ocean or a different ocean. The histogram represents the distribution of these gene fractions across all SAGs.

##### B.2. Null model for the fraction of cell pairs per gene from the same ocean, binned by nucleotide divergence

The null model for the scenario in which the Atlantic and Pacific oceans constitute a single, perfectly-mixed population is shown in Figure 4 (b) in the main text. In this case, the probability that two cells are from the same ocean is assumed independent of their divergence at any gene. The mean value of the null model is the fraction of cell pairs that are from the same ocean. The error band represents one standard deviation using a binomial distribution for the number of cell pairs from the same ocean per bin. We only consider genes longer than 700 bp – approximately the median length of core genes – to exclude genes that might be identical by chance.

### 10. Problems with Oft-used Methods: PhyML and Metagenomic Recruitment to References

In this section we highlight two practical consequences of our results for future analysis of sequencing data from *Prochlorococcus*. In the first part we show that the high rates of homoplasies within HLII make inferring phylogenies using standard algorithms unreliable. In the second part we benchmark methods for inferring the abundances of clonal subpopulations using recruitment profiles of metagenomic reads to reference genomes. Such methods are especially successful at identifying individual strain abundances in the context of the human gut microbiome, where many species consist of clonal subpopulations descended from individual colonizers (*25*, *26*). By contrast, in *Prochlorococcus*, we find that the combination of the small number of reference genomes available and recombination leads to metagenomic reads mapping nonspecifically to reference genomes, even when the entire sample consists of a single genome.

#### A. Problems with attempts to construct gene phylogenies

##### A.1. Gene trees constructed using distance based methods and with PhyML

We first use distance-based metrics to construct gene trees of the quasi-random HLII sample using an agglomerative clustering algorithm as implemented in SciPy (*20*). Here, we report the results using the average linkage criterion, but we verified that our conclusions do not change when using single or complete linkage criteria. We choose four genes among the longest ones to maximize the number of mutations used to infer the phylogenies. The resulting trees are characterized by long terminal branches and closely-spaced branch mergers (Figures S48). These features are somewhat similar to star-like trees and make inferring the tree topology challenging. Figure S49 shows the nucleotide pairwise distance matrices and the divergence distributions for the genes shown in Figures S48). Consistent with our previous conclusions, we find pairwise distances are very similar across a large fraction of the population, resulting in sharp peaks in the divergence distributions.

**Figure S48:**
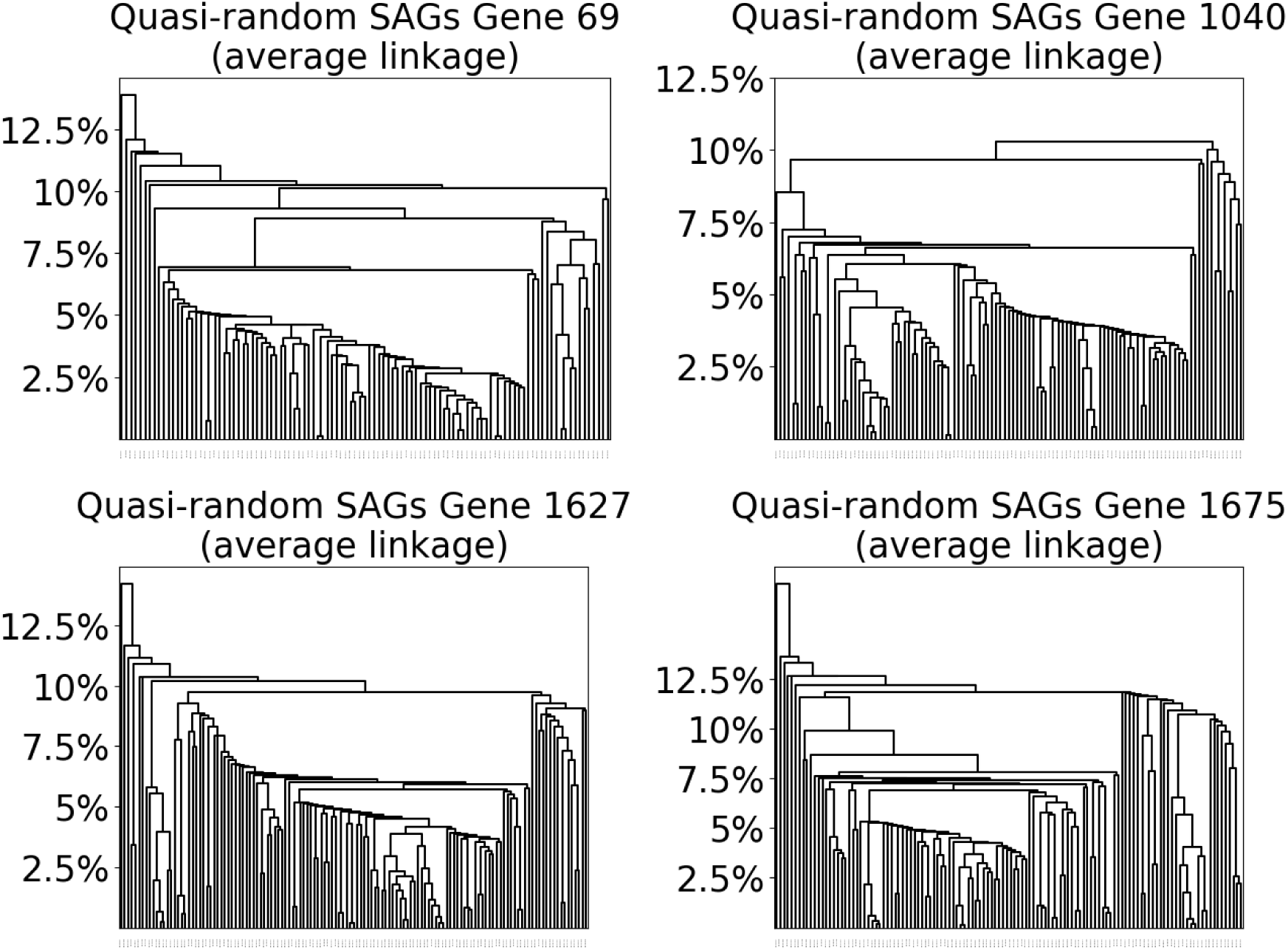
Dendrograms obtained from hierarchical clustering of nucleotide divergences of the four genes in Figures S50, S51, S52, and S53. All dendrograms were obtained using agglomerative average linkage clustering.

**Figure S49:**
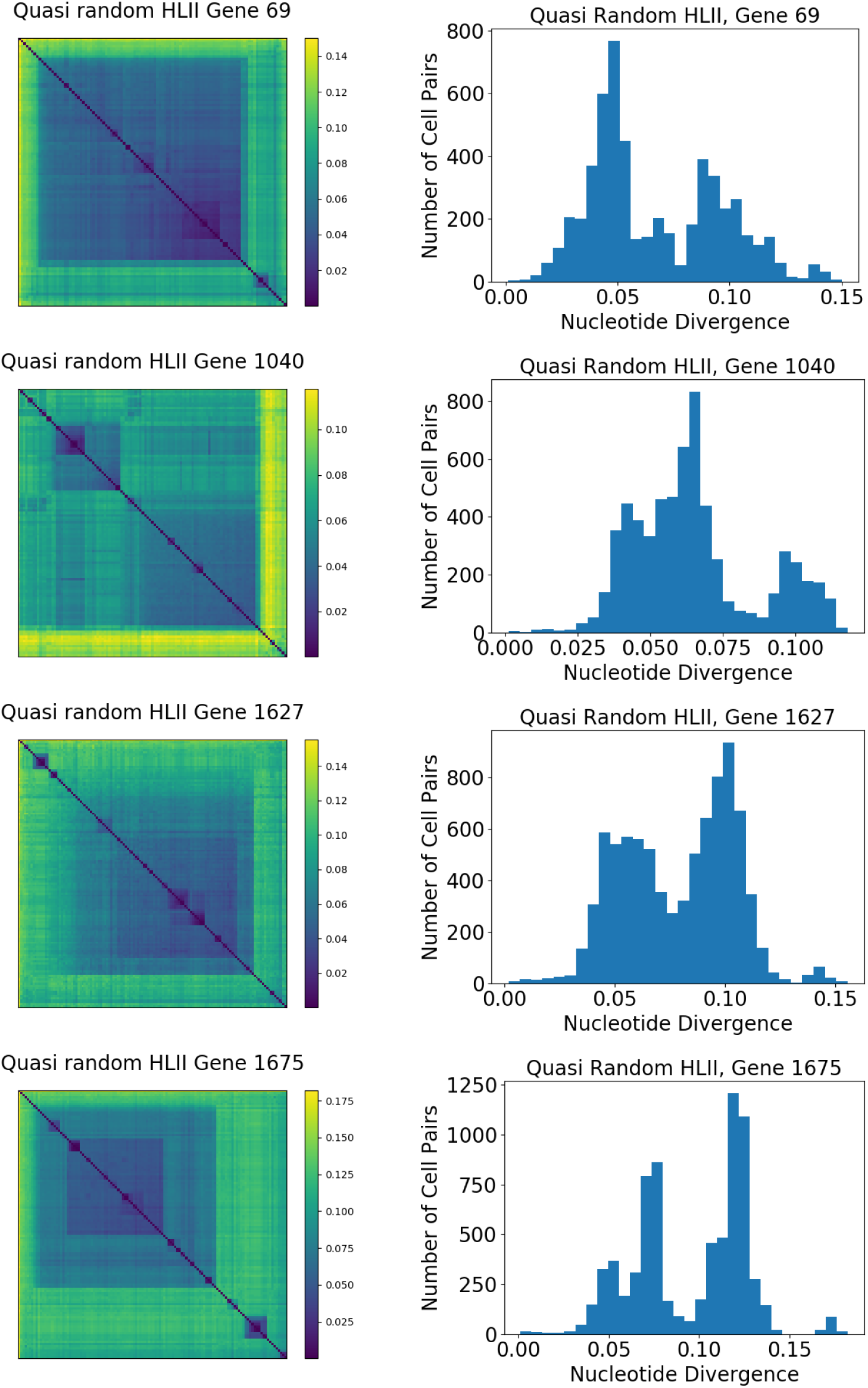
Distance matrices and cell pair divergence histograms for the four genes whose PhyML-constructed trees are visualized in Figures S50, S51, S52, and S53 are shown for comparison. Nucleotide divergence within the quasi-random HLII sample is shown, with each distance matrix having its own (linear) color scale.

We also compared the previous results to the phylogenies inferred by PhyML (*27*). The trees are shown in Figures S50, S51, S52, and S53. Default parameters were used and the program selected a mutation model that it found optimal for each alignment. Plots were generated by PhyML’s Tree Visualizer and bootstrap values are displayed at each branch (*27*). We calculated the total length of the resulting phylogenies to estimate the typical number of times a site mutated in the population. We found the lengths were between 2 and 5, indicating that multiple mutations at a given site (homoplasies) cannot be ignored. We also found many internal branches, including ones deep in the trees (close to the root), with low bootstrap values, including a some with 0.00 bootstrap support.

**Figure S50:**
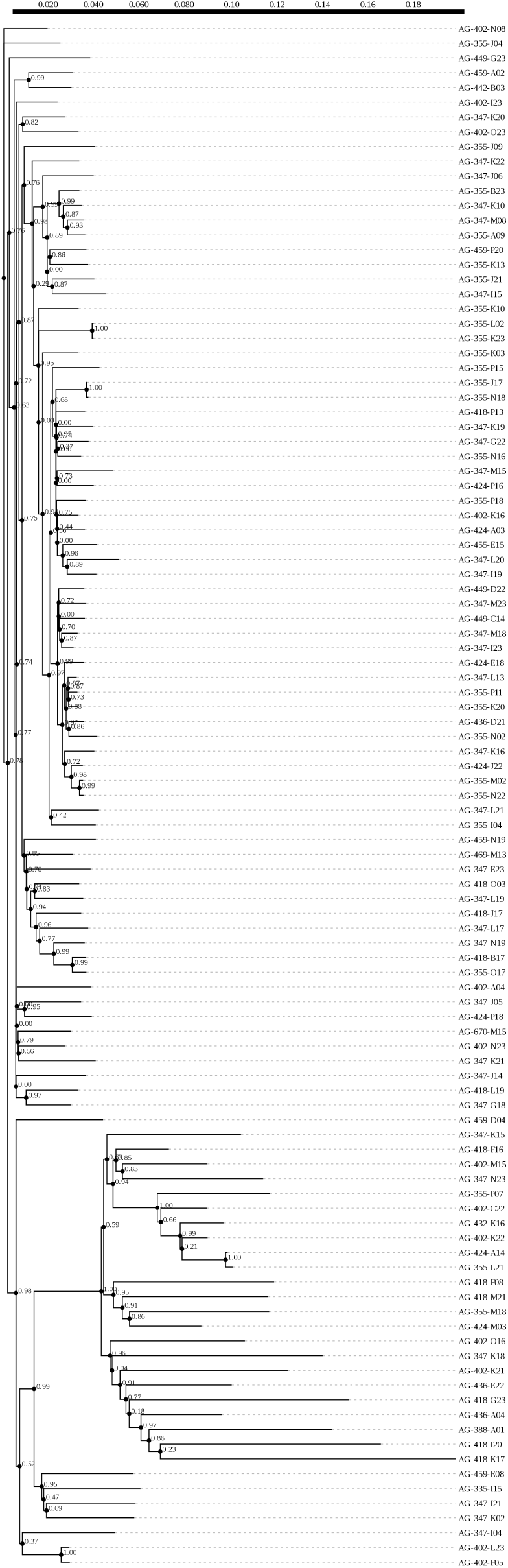
Gene 69 on the MIT9301 reference genome, 3591 bp in length and annotated as putative chromosome segregation protein, SMC ATPase superfamily. PhyML selected the “GTR +R” mutation model.

**Figure S51:**
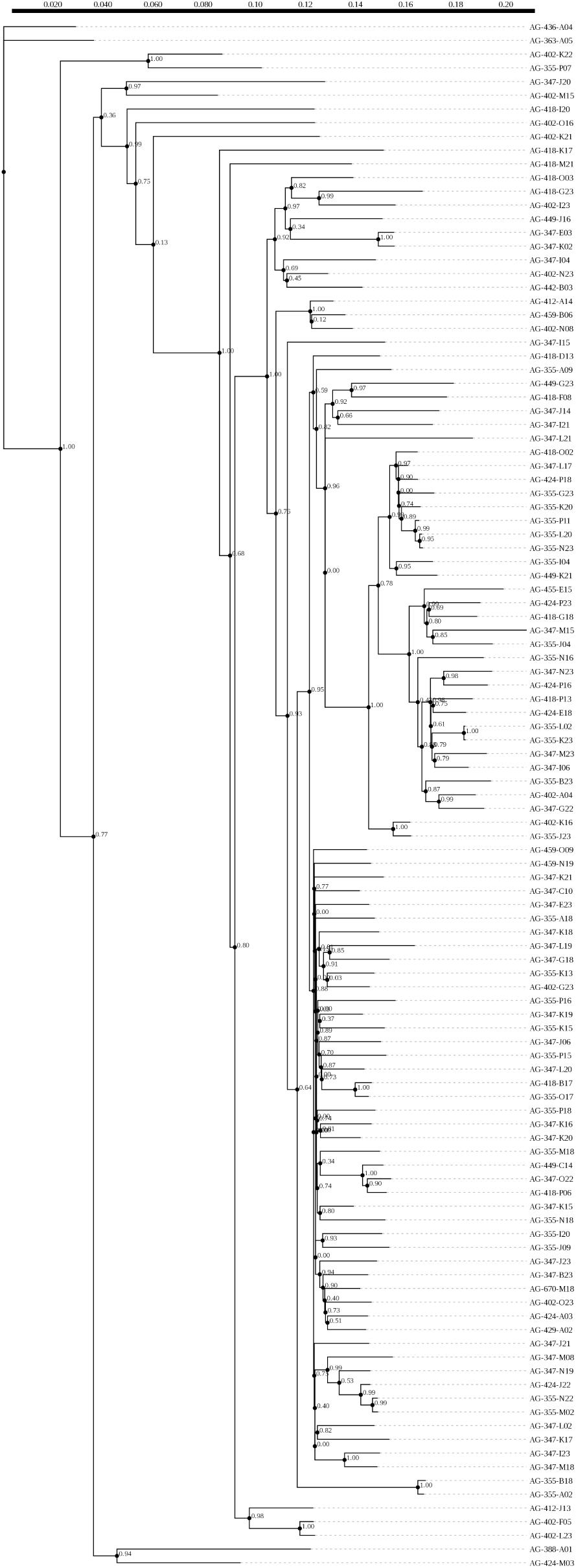
Gene 1040 on the MIT9301 reference genome, 4011 bp in length and annotated as protoporphyrin IX magnesium chelatase, subunit chlH. PhyML selected the “TN93 + R” mutation model.

**Figure S52:**
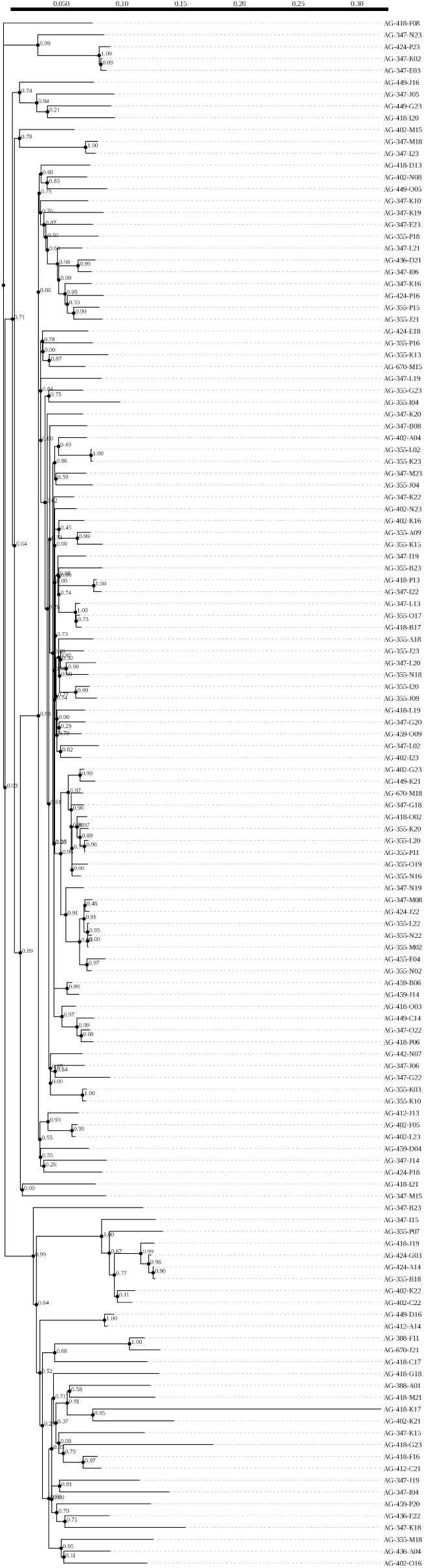
Gene 1627 on the MIT9301 reference genome, 1638 bp in length and annotated as GroEL protein (Chap-eronin cpn60). PhyML selected the “GTR +R” mutation model.

**Figure S53:**
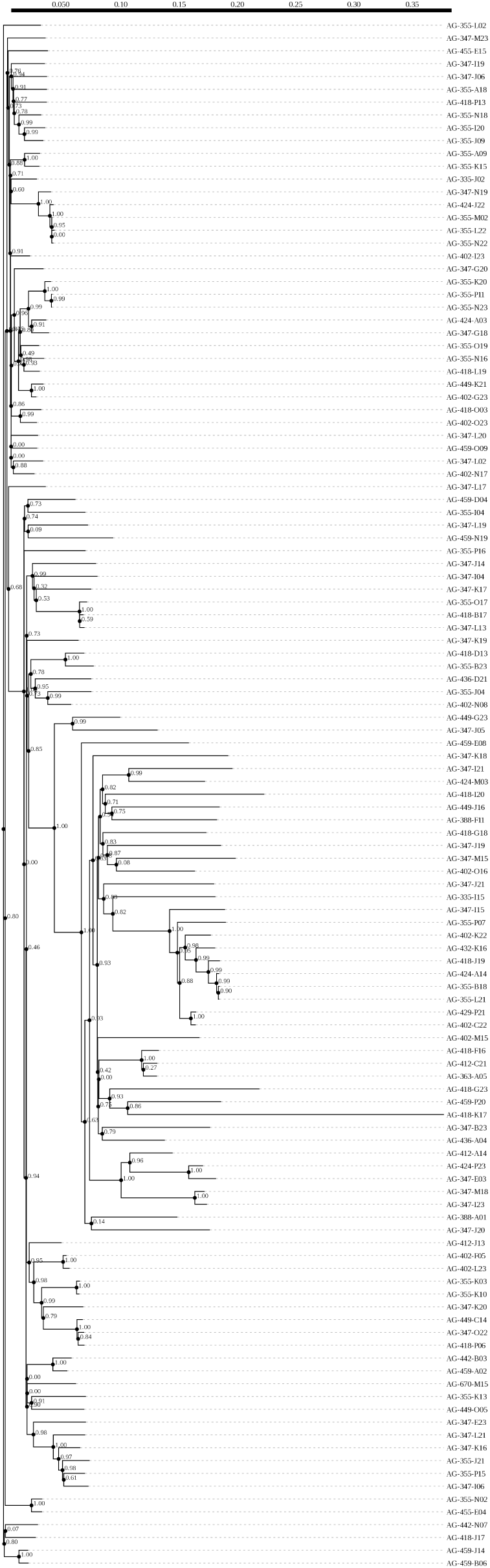
Gene 1675 on the MIT9301 reference genome, 4101 bp in length and annotated as, RNA polymerase beta prime subunit, and one gene downstream from gene rpoC1 (mentioned throughout this paper). PhyML selected the “GTR +R” mutation model.

##### A.2. SNP pair sites are incompatible with a single phylogeny on short length scales due to homoplasy

In addition to homoplasies, recombination, even if rare, can also result in errors in inferring phylogenetic trees. To investigate the likely effect of recombination, we calculate the fraction of SNP pairs that are incompatible with a single, homoplasy-free phylogeny as a function of nucleotide separation between the SNPs. Pairs of dimorphic SNPs are incompatible if all four possible combinations of alleles (e.g., AB, Ab, aB, ab) are present within the sample. Such a scenario could not have occurred without a recombination breakpoint between the SNPs or multiple mutations at at least one of the SNP sites in the sample. We use twofold degenerate synonymous dimorphic SNPs, and exclude singletons which cannot have incompatibilities by definition. SNP incompatibility rate at minimal SNP separation (3 bp here) gives an estimate of the rate of homoplasy because there is low probability of a recombination event at such short separations. An increase in the rate of incompatibility with SNP separation indicates effects of homologous recombination. The overall SNP statistics of a sample of cells determine the saturation value at large separations, expected if each SNP site represented a different tree, keeping SNP frequencies intact.

Within the whole quasi-random HLII sample, the fraction of incompatible synonymous SNP pairs does not appear to change with separation between sites. This result suggests that synonymous homoplasy is so pervasive that the rate of SNP pair incompatibilities can not be further increased by homologous recombination, even in a group as small as 80 cells used here. By comparison, nonsynonymous SNPs have lower rates of incompatibility than synonymous SNPs and are further separated from the random shuffle expectation than synonymous sites.

In the quasi-equidistant subset of HLII cells the synonymous SNP pair incompatibility rate increases from 0.37 for neighboring SNPs and increases on very slightly to 0.39 for distant SNPs, again reaching the saturation value for the dataset. Note the differences in saturation values between the different groups of cells due to differing SNP statistics and sample sizes. These results indicate that homoplasy is extensive within HLII to the extent that parsimony-based methods cannot yield an accurate tree independent of homologous recombination, which additionally contributes to SNP pair incompatibilities. For the closely-related cell group C1, the SNP pair incompatibility rate is about 0.15 for neighboring SNPs and saturates to ∼ 0.3 over 10s of kb separation. The saturation value is about 50% of the value of the unlinked null model, but the increase over short genomic distances is consistent with significant recombination even within a gene.

The several-kb lengthscale over which the incompatibility rate increases before saturating in the quasi-equidistant subset is similar to the lengthscale of SNP linkage (shown in Figure S29) for this cell group. A surprising feature is that the incompatibility rate for C1, a similar size cell group comprised of much more closely-related cells, seems to increase and roughly saturate on the same several-kb lengthscale. Why are the length scales of saturation similar for the two groups despite their significantly different divergences? Naively we might expect the incompatibility rate for C1 to increase on a longer length scale because fewer recombination events have occurred in the history of that cell group compared to the quasi-equidistant subset. We leave the resolution of this question as a topic for future work.

#### B. Recruiting metagenomic reads to reference genomes does not reflect local strain abundances

Recently, several groups have tried to infer the abundances of clonal subpopulations (strains) within *Prochlorococcus* by mapping metagenome reads to *Prochlorococcus* reference genomes (*5*, *28*, *29*). These methods assume that the fraction of reads that map to a given reference genome is proportional to the abundance of subpopulations that are asexually diverged from the reference. Below, we use synthetic metagenomes constructed from representative HLII SAGs to show that this assumption is unjustified for HLII. Specifically, we show that the fraction of reads mapping to a given reference approximates true strain abundances only when strains are ≲ 0.03 diverged from one of the references (Figures S57, S58). Comparing 168 randomly-sampled SAGs with 31 reference genomes, we find that such low divergences occur in only 5 SAG-reference pairs and are thus very unlikely in natural samples (Figure S56 left). For typical SAG-reference divergences of *d* ≈ 0.06, we find that reads map non-specifically to reference genomes (Figures S57, S59), which leads to large error in the inferred fraction of the closest reference genome (Figure S62) and to the spurious inference of the presence of a large number of references even in samples containing a single genome (Figure S60).

**Figure S54:**
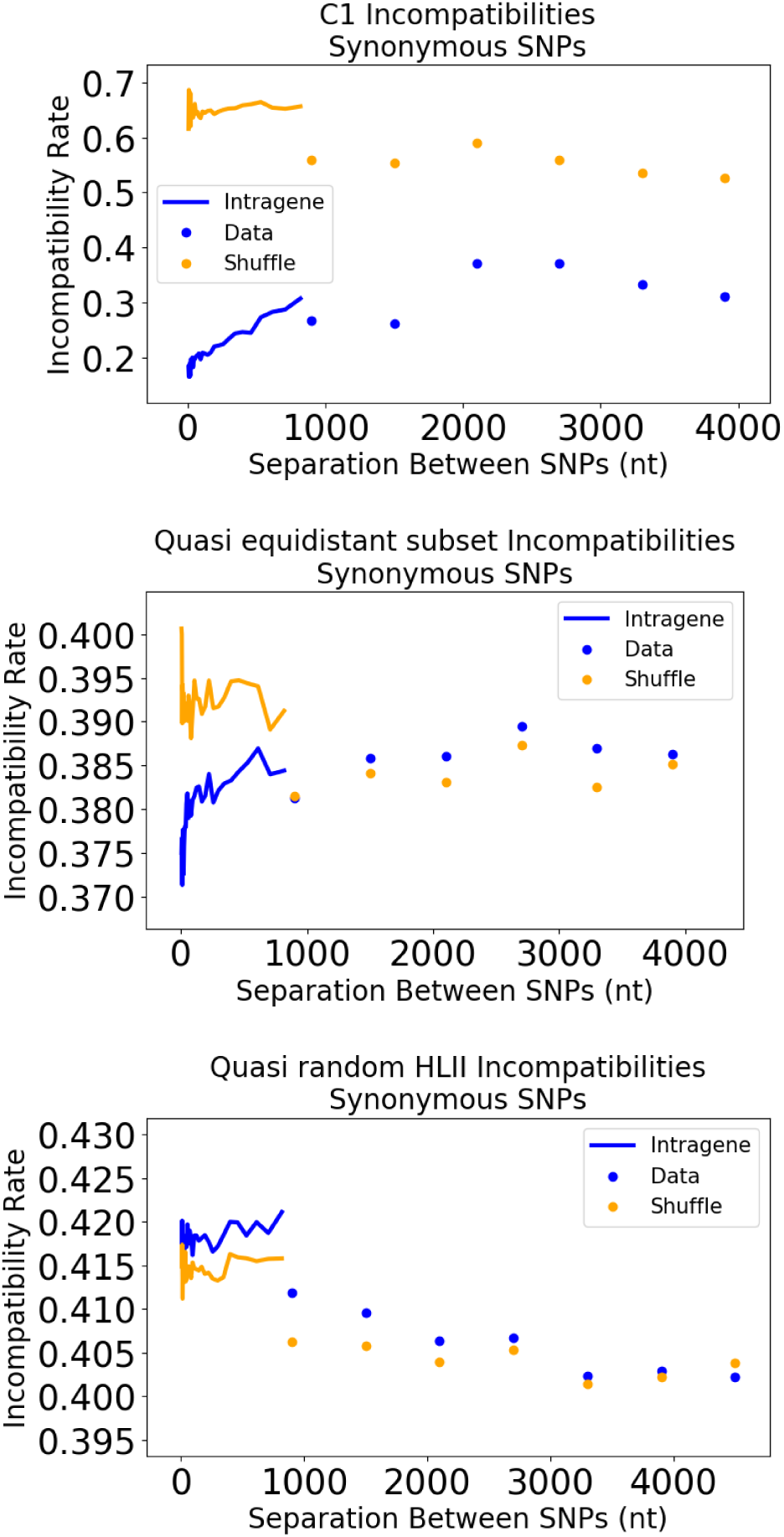
Incompatibilities of core SNP pairs (excluding singleton SNPs) at twofold degenerate sites as a function of separation between the SNP pairs on the MIT9301 reference genome, for three cell groups studied in this paper: C1 from the deeply sampled dataset (**top**), the quasi-equidistant subset (**middle**), and all quasi-random HLII SAGs (**bottom**). The continuous line at short separations represents intra-gene separations, while points represent intergene averages.

**Figure S55:**
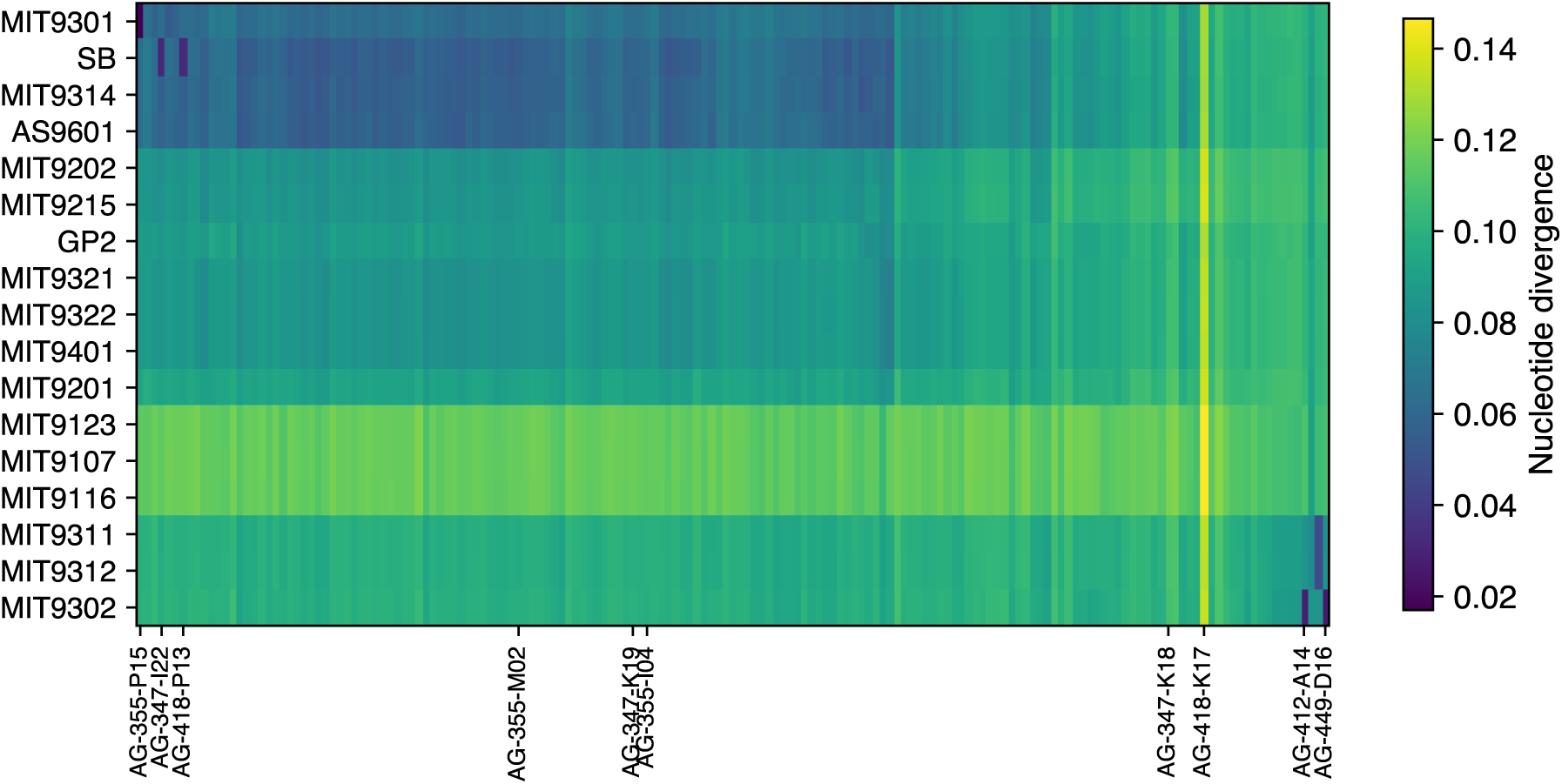
Pairwise distance matrix between 168 randomly-sampled HLII SAGs and 17 HLII reference genomes. The ten representative SAGs chosen to construct synthetic metagenomes are marked on the x-axis. Five of these chosen SAGs (AG-347-I22, AG-355-I04, AG-355-P15, AG-412-A14, AG-449-D16) have a clear closest relative among the reference genomes, with divergences low enough to share residual clonal backbones with the corresponding references (d ≲ 0.03). The remaining five SAGs span the range of divergences between SAGs and references.

**Figure S56:**
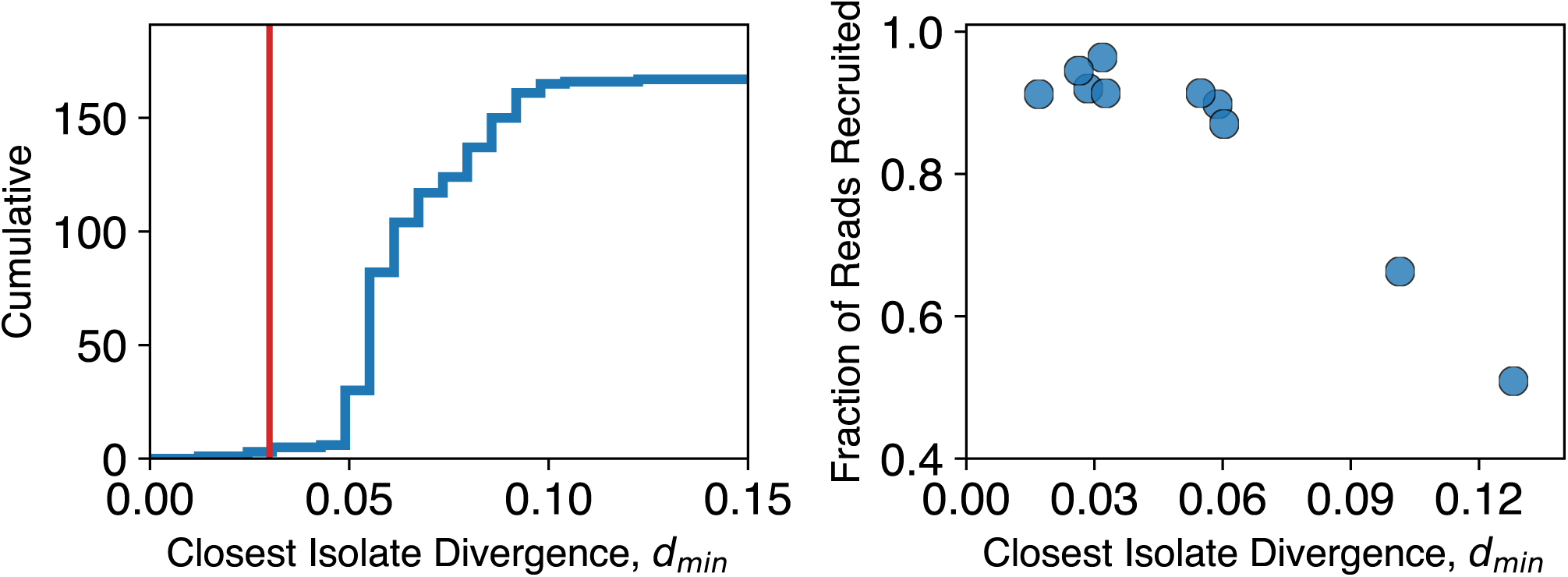
(Left) Approximately 3% of SAGs (5/168) form close pairs with one of the references (*d* ≲ 0.03). The blue line shows the unnormalized cumulative distribution of the distance to the closest reference *d_min_*, calculated from Figure S55. The red vertical line shows the empirical divergence threshold found earlier (*d* ≈ 0.03) beyond which reads from the SAGs map indiscriminantly to reference genomes. (Right) The fraction of synthetic reads recruited to *any* reference across representative SAGs decreases with divergence to the closest reference. The rest did not map to any of the references in spite of being *Prochlorococcus* reads. Synthetic metagenomes were generated from the SAGs shown in S55 as described in the text. Reads were mapped to 31 *Prochlorococcus* reference genomes using anvi’o (v8) as described in (*5*). The fraction of reads mapped was calculated from the output of bowtie2 ran with default parameters.

To understand how closely related typical *Prochlorococcus* cells from natural samples are likely to be to current reference genomes, we calculated the pairwise distances between the 168 quasi-randomly-sampled HLII SAGs and a collection of 31 reference genomes obtained from isolate strains (referred to as reference or isolate genomes interchangeably) which were used in previous metagenome studies (*5*). Distances were calculated by cutting the SAG contigs into 1kbp segments and aligning the segments to contigs from the reference genomes using BLAST (`-task blastn -word_size 12 -evalue 1E-50`). For each 1 kbp segment, the best hits that aligned over at least 750 bp were kept and averaged to obtain the mean nucleotide divergence. We found that this simple procedure agreed very well with the averages obtained from core genes calculated previously. In all cases, we found that the vast majority of reads (*>* 98%) obtained from HLII SAGs mapped to HLII references. These results show that abundances of different *Prochlorococcus* clades can reliably be inferred from metagenome read recruitment. We therefore focus on recruitment patterns to just the 17 HLII reference genomes for most of the analysis (Figure S55).

The pairwise divergences between the quasi-random SAGs and the 17 HLII references are shown Figure S55. Most SAGs were at least 6% diverged from the closest reference, which is consistent with the typical nucleotide divergences within HLII (Figure S12). We found 5 SAGs with divergences to the closest reference between 0.017 and 0.033 (Figure S56 left), that were noticeably closer to a reference than typical SAGs, although still considerably more diverged than C1 SAGs (Figure S12). Note that the small fraction of SAGs (∼ 3%) closely-related to a reference is consistent with the fraction of HLII cells in tight clusters in the quasi-random dataset shown in Figure S14. We chose the 5 SAGs closely-related to a reference together with 5 other typical SAGs and used them to create synthetic metagenomes to test the read mapping methods used previously.

We generated synthetic reads by randomly cutting SAG contigs into 150 bp segments and assigning a constant quality score of 40 to all bases, corresponding to an error rate of 10^−4^. These numbers were chosen to be comparable to data from the Tara Oceans expedition, which contains 2 × 101 bp pairedend reads with an average quality score of 38 (*30*). Note that we do not include sequencing errors or any sampling biases in our synthetic read construction so our tests represent a best case scenario. The synthetic reads were combined into a single FASTQ file for each SAG, with the number of reads chosen such that the average coverage at each site along the genome was 100. We call these samples single-SAG metagenomes. Metagenomes containing reads from multiple SAGs were constructed by sampling single-SAG metagenomes in different proportions, as explained below. We recruited reads to a collection of 31 *Prochlorococcus* reference genomes using anvi’o (v8), following Ref. (*5*). In most cases we found ∼ 90% of reads aligned to at least one reference, but we found that this fraction decreased rapidly once the distance to the closest reference from the SAG approached *d_min_* ≈ 0.1 (Figure S56 right).

To verify the accuracy of the mapping, we compared the fraction of reads mapped to each reference genome for the different single-SAG metagenomes. If individual strains can reliably be identified from short reads, we expect the fraction of reads recruited across the reference genomes to be sharply peaked around the closest reference to the strain. For the 5 SAGs that were closely-related to one of the reference genomes, we indeed found such a peak (Figure S57). But even in these cases, we found up to 40% of reads mapped to other reference genomes. For example, reads from AG-418-P13 mapped to three other references, each recruiting approximately 10% of reads.

**Figure S57:**
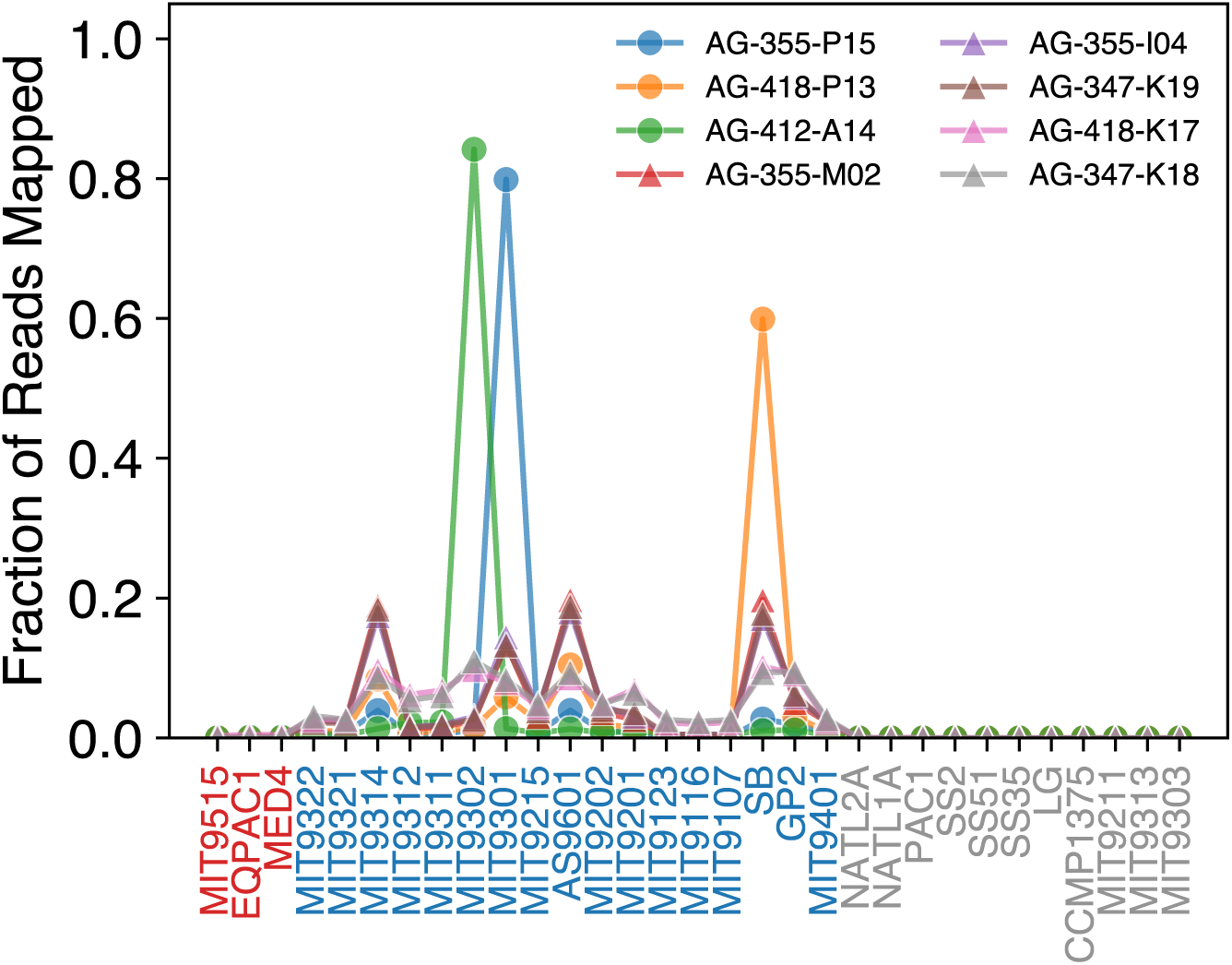
Recruitment of synthetic metagenome reads shows distinct patterns for SAGs with and without close references in the database. Synthetic metagenomes were generated from single SAGs as described in the text. Reads were mapped onto a collection of 31 reference genomes, consisting of 3 HLI (red), 17 HLII (blue), and 11 LL (gray) isolates, using anvi’o (v8) as described in (*5*). The fraction of reads mapped to each reference out of the total number of reads mapped, as reported by anvi’o, is shown. Recruitment profiles shown for 8 of the 10 single-SAG metagenomes. Only comparisons to HLII references, which account for *>* 98% of recruited reads, are shown. Metagenomes from SAGs with *d* ≲ 0.03 from one of the reference genomes are shown with circles while other SAGs are shown with triangles.

The reads mapped to other reference genomes could be due to a small number of flexible genes not shared with the closest reference. Alternatively, they could be due to short core gene segments that map better to other references, possibly as a result of recombination in the population. To distinguish between these two scenarios, we investigated the coverage across individual genes for each reference. The coverage across individual genes showed multiple references, other than the one closest to the SAG, recruited reads over a large fraction of their genome, with coverages comparable to the target coverage (Figure S58). These results are consistent with these reads belonging to core genome regions that recombined with other strains since their common ancestor with the closest reference. We can estimate the fraction of the genome that recombined for these pairs from our analysis of C1 from the main text. The divergence between genome pairs that share an asexual backbone is approximately *d* ≈ *f_rec_d_rec_* + (1 − *f_rec_*)*d_asex_*, where *f_rec_* is the fraction of the genome that is recombined, *d_rec_* is the typical divergence between recombined segments and *d_asex_*is the divergence of the asexual backbone. For C1, we found *f_rec_*increased approximately linearly with *d_asex_*, with a slope of approximately 30 (Figure 2c, main text). We also found that *d_rec_* was close to the typical HLII divergence of 6%. Using these values and the SAG-reference divergences for the close pairs of *d* ≈ 3%, we obtain *f_rec_* ∼ 0.5, which is quite close the observed fraction of reads that map to other genomes. These results suggest that the closest SAG-reference pairs may share an asexual backbone, but have still undergone substantial recombination since their common ancestor.

**Figure S58:**
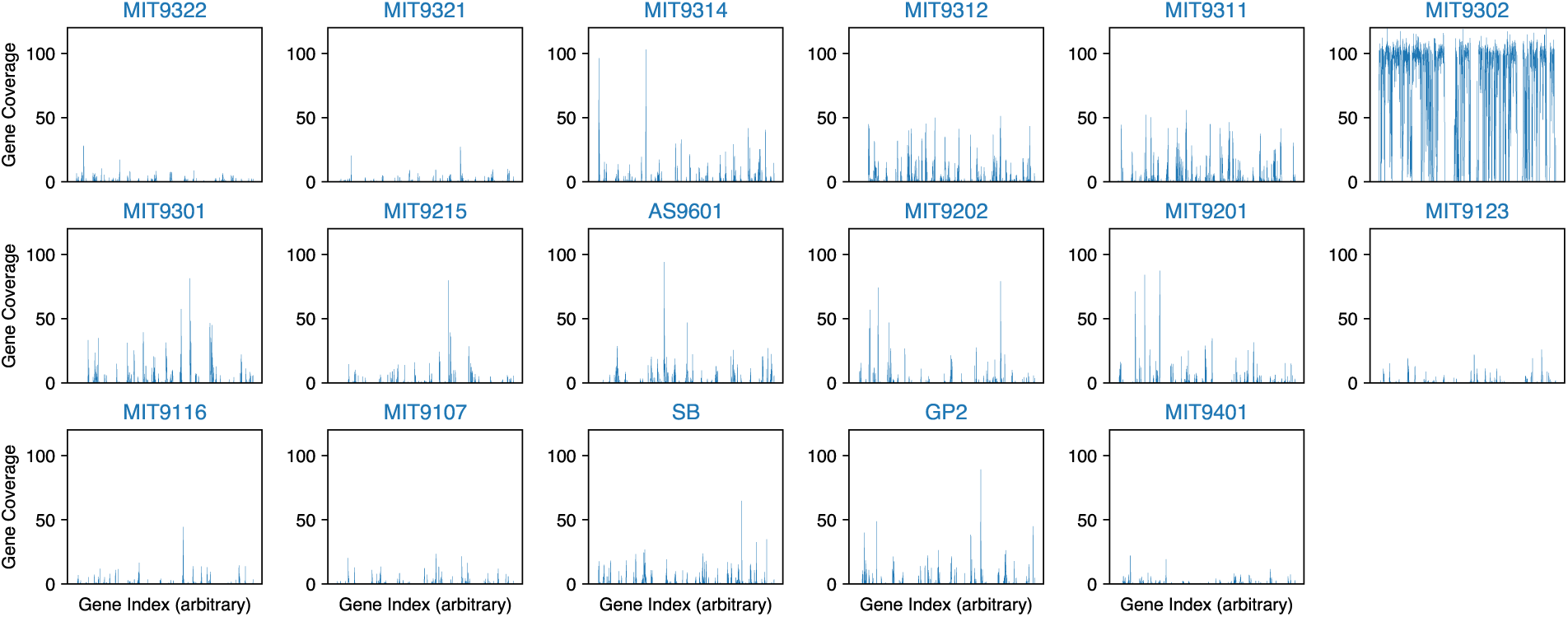
Read coverage across core genes for each reference from the AG-412-A14 (*d* ≈ 0.03 from MIT9302) synthetic metagenome. The 17 HLII reference genome out of 31 total are shown in different panels. In each panel, the gene coverage as reported by anvi’o is shown on the y-axis and the gene index assigned by anvi’o is show on the x-axis. The synthetic metagenome was generated with a target coverage of 100.

By contrast, reads from typical SAGs mapped non-specifically to a large number of reference genomes. We found that no single reference recruited more than ≈ 20% from the typical SAG chosen. The coverage profiles over individual genes from the reference genomes confirmed the high coverage over a large fraction of the genomes of multiple references (Figure S59). These results suggest that even for a single SAG, multiple references genomes would be inferred in the sample. We verified this conclusion by estimating the fraction of each reference in the sample using the mean Q2-Q3 coverage reported by anvi’o. Specifically, for each reference, the average coverage between the 25 and 75 percentiles (second and third quartiles) of the gene coverage distribution is calculated. We found that this method provided an accurate predictor of frequency for SAGs with low divergences to one of the references (Figure S60 left). By contrast, for typical SAGs we found between 5-16 references were detected with fraction *>* 0.01 (Figure S60). The total number of references detected increased with the distance to closest reference, consistent with genomes being roughly random mixtures of segments from population on scales larger than 10 genes (Figure S60 right).

**Figure S59:**
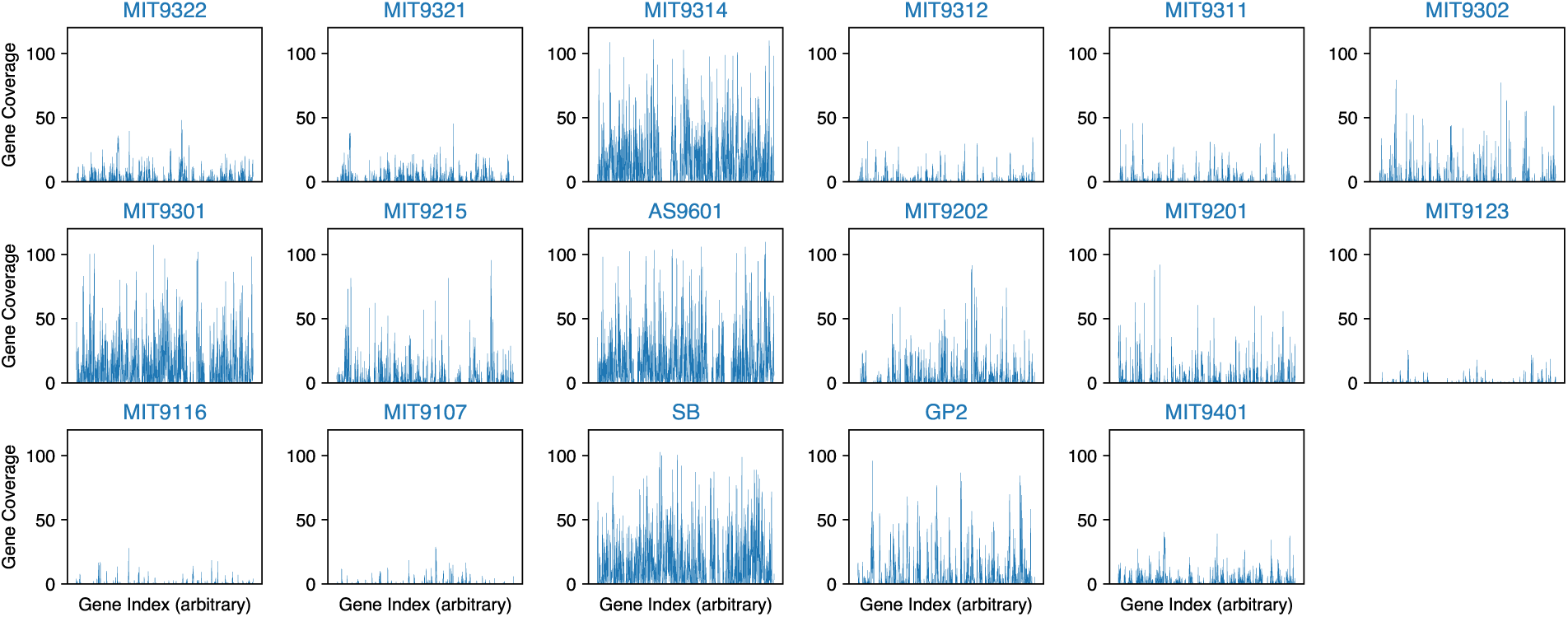
Read coverage across core genes for each reference from the AG-355-I04 (typical SAG) synthetic metagenome. Panels were made as in S58. The synthetic metagenome was generated with a target coverage of 100. Note the high coverage across a large fraction of most HLII genomes.

**Figure S60:**
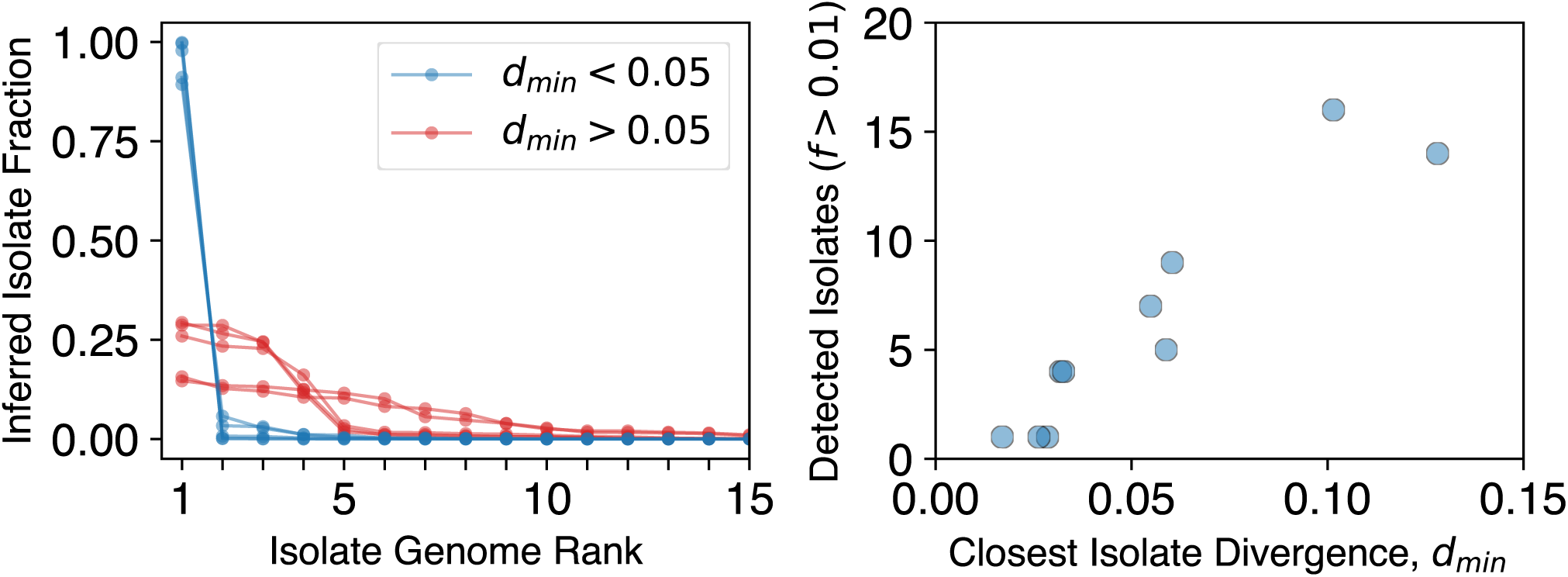
Between 1/3 and 2/3 of HLII reference genomes are detected at significant abundances in typical single-SAG synthetic metagenomes. **(Left)** The inferred fraction of reference genomes ranked according to their fraction in single-SAG synthetic metagenomes. The inferred fraction of each reference was estimated from the Q2-Q3 coverage reported by anvi’o as described in the text. The five metagenomes generated from SAGs with unusually low divergences from one of the references are shown in blue, with the other five shown in red. **(Right)** The number of reference genomes with an inferred fraction *>* 0.01 as a function of the distance to the closest reference genome for the ten single-SAG metagenomes. Note that for typical distances to the closest reference (0.05 ≲ *d* ≲ 0.1), between 5 and 16 out of 17 HLII references appear to be detected.

Next, we checked how recruitment profiles were affected by the addition of multiple SAGs to the metagenomes. We generated multi-SAG metagenomes by randomly sampling a total of 500,000 reads from the single-SAG metagenomes in different proportions. We chose three SAGs that were closely related to MIT9301, MIT9302, and SB and created synthetic metagenomes of all pairs with the fraction of reads of 0.01, 0.1, 0.5 for each SAG. In addition, we constructed metagenomes with all 3 SAGs plus 0, 1, 2, or 5 other SAGs, for a total of 19 synthetic metagenomes. The fraction of reads recruited to MIT9301, MIT9302, and SB were similar to the proportions of the 3 closely-related SAGs in the metagenome, consistent with our earlier results for single-SAG metagenomes (Figure S61). However, we did not observe any qualitative differences between metagenomes with 3, 4, 5, or 10 SAGs. Thus, even in cases where a significant fraction of the sample contains genomes closely-related to the reference, read recruitment can severely underestimate the number of distinct genomes present.

**Figure S61:**
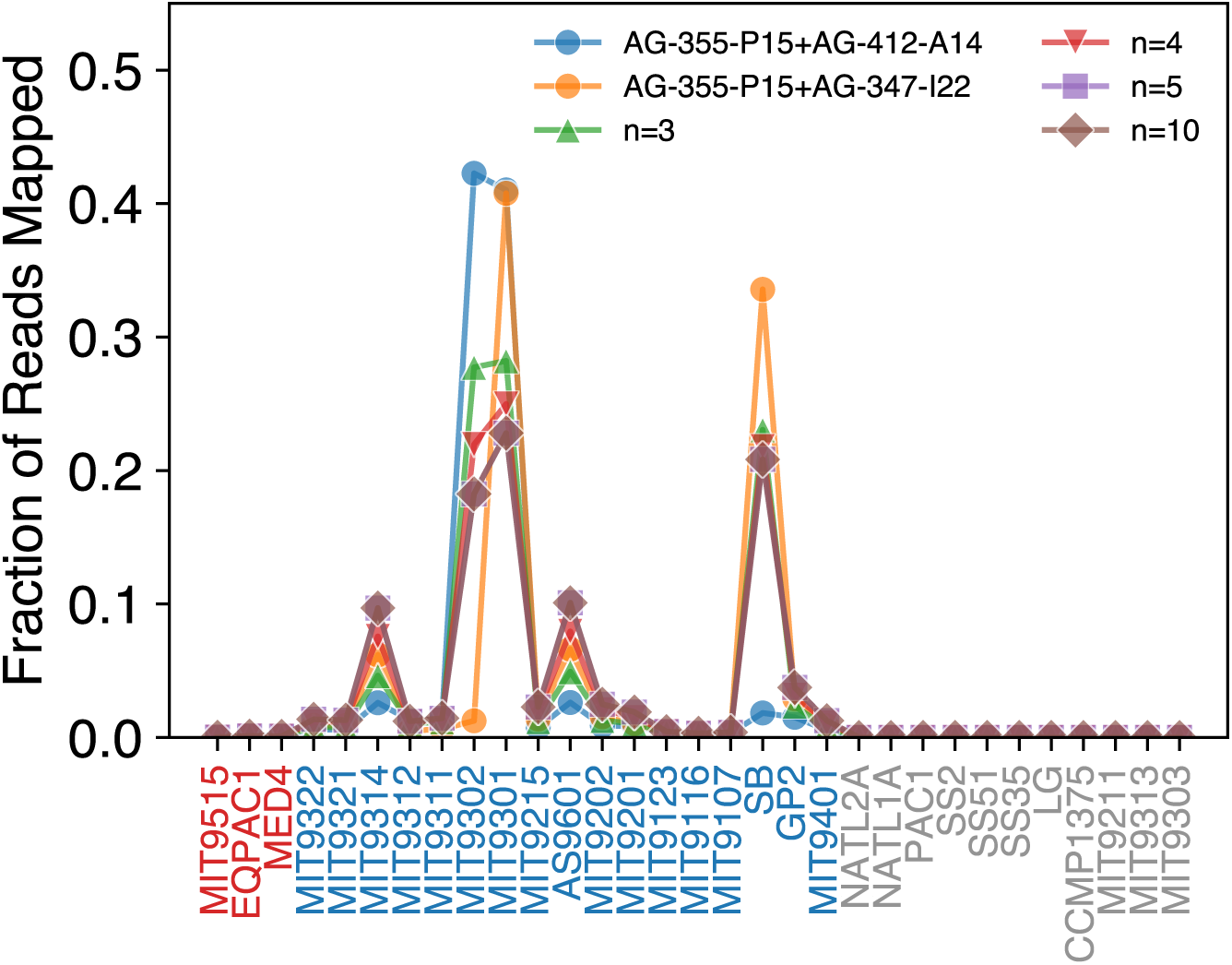
Read recruitment patterns cannot resolve the number of genomes in the sample. Same as Figure S57, but using synthetic metagenomes with reads from multiple SAGs. Metagenomes from pairs of SAGs with *d* ≲ 0.03 from one of the references are shown with circles. Example metagenomes with 3 (upward triangle), 4 (downward triangle), 5 (square), and 10 (diamond) SAGs are also shown for comparison. An equal number of reads from each single-SAG metagenome were sampled to create metagenomes with *>* 1 SAG. All metagenomes with 3 or more SAGs contained AG-355-P15, AG-347-I22, and AG-412-A14.

Finally, we asked how well does the fraction of reads mapped to the closest reference for each SAG approximate the SAG abundance in the sample. For each SAG in the 29 synthetic metagenomes, we identified the closest reference and inferred its abundance using the mean Q2-Q3 coverage method described earlier. We found that inferred abundance of references with closely-related SAGs approximated fairly well the true value (Figure S62 left), consistent with our earlier results. But even here, the relative error could be quite high if SAGs were present at low abundances (Figure S62 right). For typical SAGs, the differences between the inferred and true relative abundances were close to one. These results show that inferred relative abundances from metagenome samples are only reliable when a reference that is closer than 3% nucleotide divergence is available and the strain has abundances close to 10% or higher in the sample. These circumstances never appear in our randomly-sample SAG dataset (Figure S4) suggesting that they are very unlikely in natural samples from the ocean.

**Figure S62:**
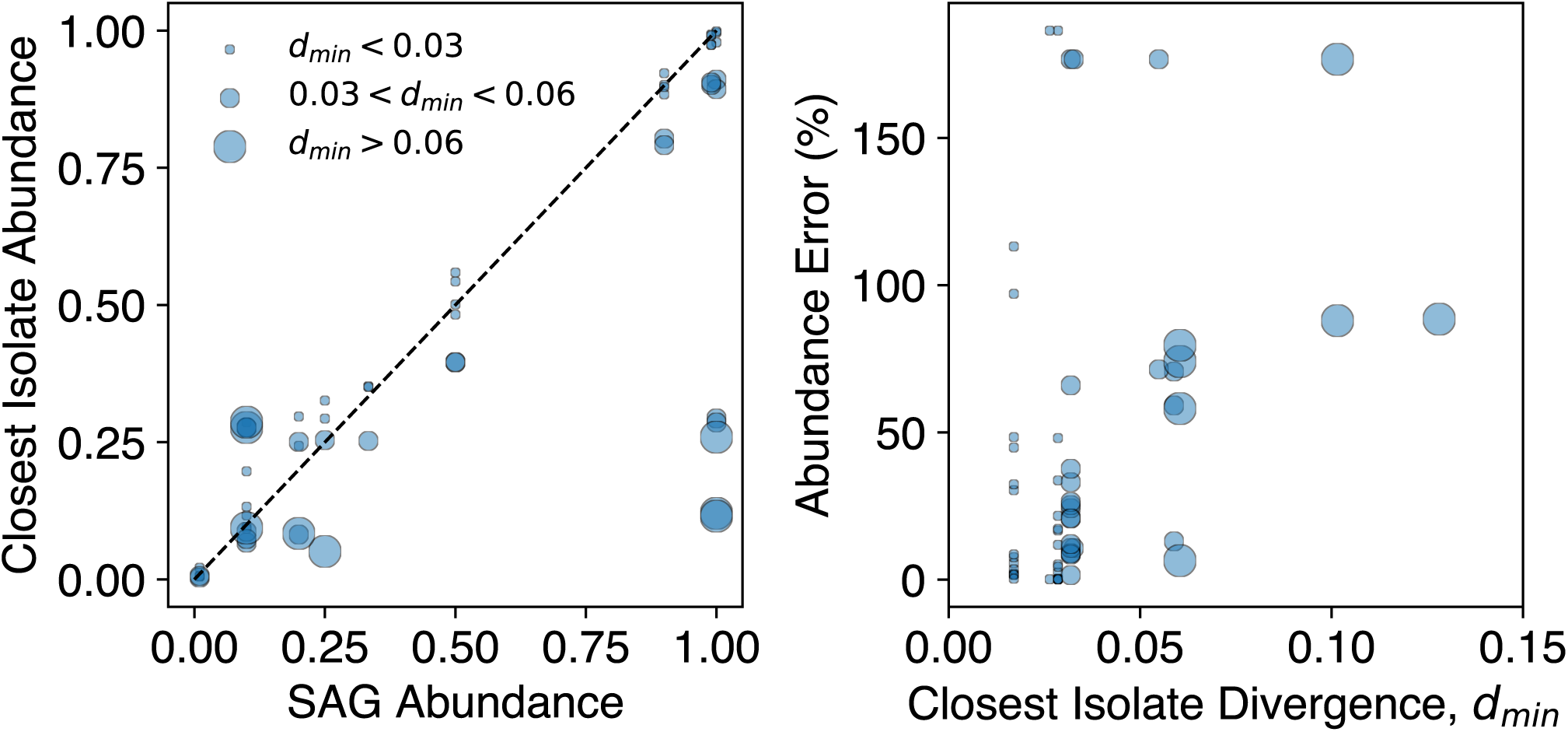
Relative abundance of closest reference is poorly correlated with SAG abundances in synthetic metagenomes for typical SAGs. **(Left)** A total of 29 synthetic metagenomes were generated containing reads from 1-10 SAGs with different relative abundances, as described in the text. For each SAG, the closest reference was determined and the relative abundance of the reference was calculated from the Q2-Q3 coverage reported by anvi’o. Symbols of different sizes denote SAGs with distances to the closest reference *d_min_*in different ranges. **(Right)** The relative error in the relative abundance of the closest reference shown in the left panel as a function of *d_min_*.

We note that in both single-SAG and multi-SAG metagenomes, we observed a fairly high correlation in the recruitment pattern of different SAGs that had typical divergences to other reference genomes (Figure S57, S61). This high correlation is consistent with the residual linkage correlation. *R*, at long genomic distances reported in the main text. But note that these SAGs do not form a cluster, as the divergences between them are comparable to the divergences to the references. Thus there is no simple mapping from the recruitment profiles to individual genomes.

